# Evaluating and improving heritability models using summary statistics

**DOI:** 10.1101/736496

**Authors:** Doug Speed, John Holmes, David J Balding

## Abstract

There is currently much debate regarding the best way to model how heritability varies across the genome. The authors of GCTA recommend the GCTA-LDMS-I Model, the authors of LD Score Regression recommend the Baseline LD Model, while we have instead recommended the LDAK Model. Here we provide a statistical framework for assessing heritability models using summary statistics from genome-wide association studies. Using data from studies of 31 complex human traits (average sample size 136,000), we show that the Baseline LD Model is the most realistic of the existing heritability models, but that it can be improved by incorporating features from the LDAK Model. Our framework also provides a method for estimating the selection-related parameter α from summary statistics. We find strong evidence (P<1e-6) of negative genome-wide selection for traits including height, systolic blood pressure and college education, and that the impact of selection is stronger inside functional categories such as coding SNPs and promoter regions.

---

Previously,^1^ we explained how the softwares GCTA^2^ and LD Score Regression^3^ (LDSC) derive their estimates from an underlying heritability model, and demonstrated how changing this model can lead to very different estimates of SNP heritability, confounding bias and heritability enrichments.^4^ Originally, both GCTA and LDSC assumed that each SNP is expected to contribute equal heritability, which we refer to as the GCTA Model.^5^ We showed that it is more realistic to assume the LDAK Model, where expected heritability varies with both linkage disequilibrium (LD) and minor allele frequency (MAF).^1,6^ Now the authors of GCTA recommend using the GCTA-LDMS-I Model,^7,8^ which partitions the GCTA Model based on MAF and LD, while the authors of LDSC recommend using the Baseline LD Model,^9,10^ which adds to the GCTA Model 74 SNP annotations based on LD, MAF and functional classifications (when used with this model, LDSC is referred to as S-LDSC).

Our earlier work^1^ compared the GCTA and LDAK Models based on the likelihood from restricted maximum likelihood^11^ (REML). However, this approach requires access to individual-level data and is only feasible for relatively simple heritability models. We now propose an approximate model likelihood that can be computed from genome-wide association study (GWAS) summary statistics and for highly complex heritability models.

## Results

For our main analysis, we use summary statistics from 31 GWAS to compare twelve heritability models (Table 1). We perform this analysis using SumHer, part of our software package LDAK (see URLs).^4,6^ Here we briefly describe the heritability models, our proposed model likelihood and the data we use; for full details see Online Methods.

**Table 1.**
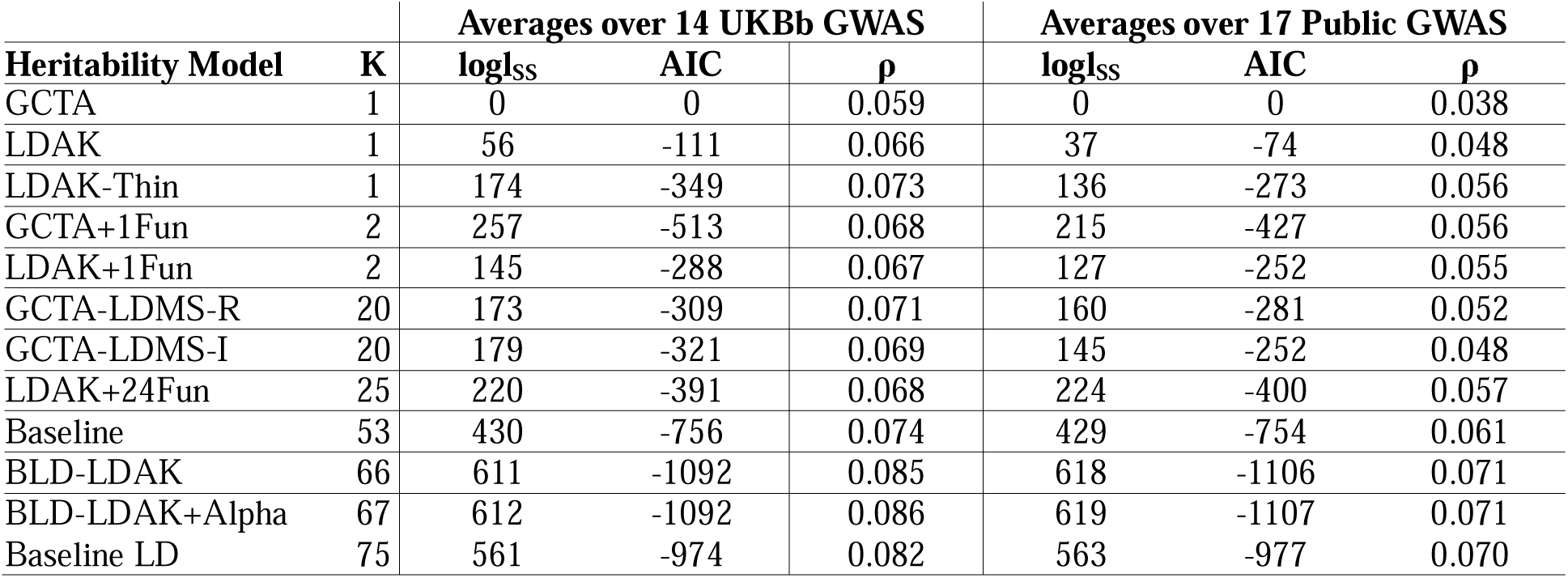
Performance of heritability models. K is the number of parameters. logl_SS_ is the approximate model log likelihood (relative to the GCTA Model), AIC is the Akaike Information Criterion (also relative to the GCTA Model), and ρ is the weighted correlation between observed and predicted test statistics from leave-one-chromosome-out prediction; values are averaged across either the 14 UKBb or 17 Public GWAS. The s.d. of average ρ is always 0.001. See main text for details of the heritability models. There are 24 versions of the GCTA+1Fun and LDAK+1Fun Models (one for each function indicator); values here correspond to the versions with highest average logl_SS_.

### Heritability models

The heritability model specifies how E[h^2^_j_], the expected heritability contributed by SNP j, varies across the genome. We consider nine existing heritability models. The one-parameter GCTA Model^5^ assumes E[h^2^_j_] is constant. The 20-parameter GCTA-LDMS-R^7^ and GCTA-LDMS-I^8^ Models both partition the genome based on MAF and LD, then assume E[h^2^_j_] is constant within each bin. The 53-parameter Baseline Model^9^ extends the GCTA Model by adding 52 functional annotations; these include 24 function indicators that can be used to estimate functional enrichments (the heritability enrichments of functional categories of SNPs). The 75-parameter Baseline LD Model^10^ adds to the GCTA Model six LD-related annotations, ten MAF indicators and 58 functional annotations (including the 52 functional annotations of the Baseline Model). The two-parameter GCTA+1Fun Model^12^ adds to the GCTA Model one function indicator from the Baseline LD Model (there are 24 versions of this model, depending on which indicator is added). The one-parameter LDAK Model^1,6^ assumes E[h^2^_j_] is proportional to w_j_[f_j_(1-f_j_)]^0.75^, where w_j_ is the LDAK weighting of SNP j (w_j_ tends to be higher for SNPs in low-LD regions) and f_j_ is its MAF. The two-parameter LDAK+1Fun^1^ and 25-parameter LDAK+24Fun^4^ Models extend the LDAK Model by adding either one or all 24 function indicators of the Baseline Model.

We construct three novel heritability models. The 66-parameter BLD-LDAK Model combines features of the Baseline LD and LDAK Models; first we add to the Baseline LD Model the LDAK weighting w_j_, then we remove the ten MAF indicators and scale the remaining 66 annotations by [f_j_(1-f_j_)]^0.75^. The 67-parameter BLD-LDAK+Alpha Model is the same, except it scales the annotations by [f_j_(1-f_j_)]^1+ α^, where α is estimated from the data.

The one-parameter LDAK-Thin Model is a simplified version of the LDAK Model, obtained by setting the LDAK weightings to either zero or one.

### Measuring model fit

Suppose we have summary statistics from a GWAS; let S_j_ denote the χ^2^(1) test statistic from regressing the phenotype on SNP j. Suppose also we have genotype data from an ancestrally-matched reference panel, from which we can estimate r^2^_jl_, the squared correlation between SNPs j and l. The authors of LDSC^3^ derived that the marginal distribution of each S_j_ is approximately gamma with shape ½ and scale 2E[S_j_], where E[S_j_] is the expectation of S_j_ (a function of the parameters of the chosen heritability model). Based on this, we propose the approximate joint log likelihood

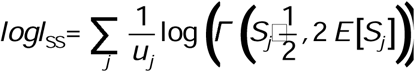

where

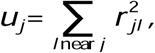

and Γ(X|a,b) is the probability density function of a gamma distribution with shape a and scale b. The weights 1/u_j_ are the same as those used by LDSC when regressing S_j_ onto E[S_j_], and are included to allow for correlations between local SNPs.^3^

We perform three analyses to support the use of logl_SS_ to compare heritability models. Firstly, Supplementary Fig. 1 shows that for scenarios where both logl_SS_ and the REML likelihood can be computed, they are concordant. Secondly, Supplementary Fig. 2 shows that when we add a non-informative annotation to a heritability model, twice the increase in logl_SS_ is approximately χ^2^(1) distributed (the expected distribution were logl_SS_ an exact likelihood). Thirdly, Table 1 shows that the ranking of heritability models based on logl_SS_ is consistent with the ranking based on leave-one-chromosome-out prediction of test statistics. Additionally, Supplementary Table 1 shows that the ranking of models is unchanged if we instead compute an unweighted version of logl_SS_ using only SNPs in approximate linkage equilibrium and u_j_=1.

### Data

We use summary statistics from two sets of GWAS. The first set are 14 traits from UK Biobank (UKBb):^13,14^ eight continuous (body mass index, forced vital capacity, height, impedance, neuroticism score, pulse rate, reaction time and systolic blood pressure), four binary (college education, ever smoked, hypertension and snorer) and two ordinal (difficulty falling asleep and preference for evenings). We performed these GWAS ourselves; after stringent quality control, 130k samples and 4.7M SNPs remained. We additionally use summary statistics from 17 Public GWAS:^4,15^ ten continuous (including anthropometric measures and psychiatric scores) and seven binary (mostly complex diseases). The average sample size is 141k (range 21-329k). As a reference panel, we use 489 European individuals from the 1000 Genome Project,^16^ recorded for 10.0M SNPs (MAF>0.005).

### Performance of heritability models

Table 1 reports logl_SS_ for the twelve heritability models, averaged across either the 14 UKBb or 17 Public GWAS (values for individual GWAS are in Supplementary Table 2). We rank models based on the Akaike Information Criterion^17^ (AIC), equal to 2K-2logl_SS_, where K is the number of parameters.

When we restrict to the nine existing heritability models, the Baseline LD Model performs best; it has average AIC 221 lower than the next best model and is the top-ranked model for 28 of the 31 GWAS. However, when we consider all twelve heritability models, the BLD-LDAK and BLD-LDAK+Alpha Models are the best; they both have average AIC 124 lower than the Baseline LD Model, and now these are the top two models for 28 of the 31 GWAS. These two models would remain the best if instead of the AIC, we ranked models based on -2 logl_SS_ or 4K-2logl_SS_ (i.e., either removed or doubled the penalty on parameters).

### Genetic architecture estimates

Figure 1, Supplementary Fig. 3 and Supplementary Tables 3-6 compare estimates of SNP heritability, confounding bias, and functional enrichments from the twelve heritability models (note that only the seven models that include function indicators can be used to estimate functional enrichments). As heritability models have become more complex, estimates have tended to converge (the more complex models produce estimates of SNP heritability and confounding bias intermediate between those from the GCTA and LDAK Models, and estimates of functional enrichments intermediate between those from the GCTA+1Fun and LDAK+1Fun Models). Based on model fit (Table 1), we consider the BLD-LDAK and BLD-LDAK+Alpha Models to be most reliable; their estimates of confounding bias and functional enrichments are close to those from the Baseline LD Model, while their estimates of SNP heritability tend to be between those from the LDAK Model and those from the Baseline LD or GCTA-LDMS-I Models.

**Fig. 1.**
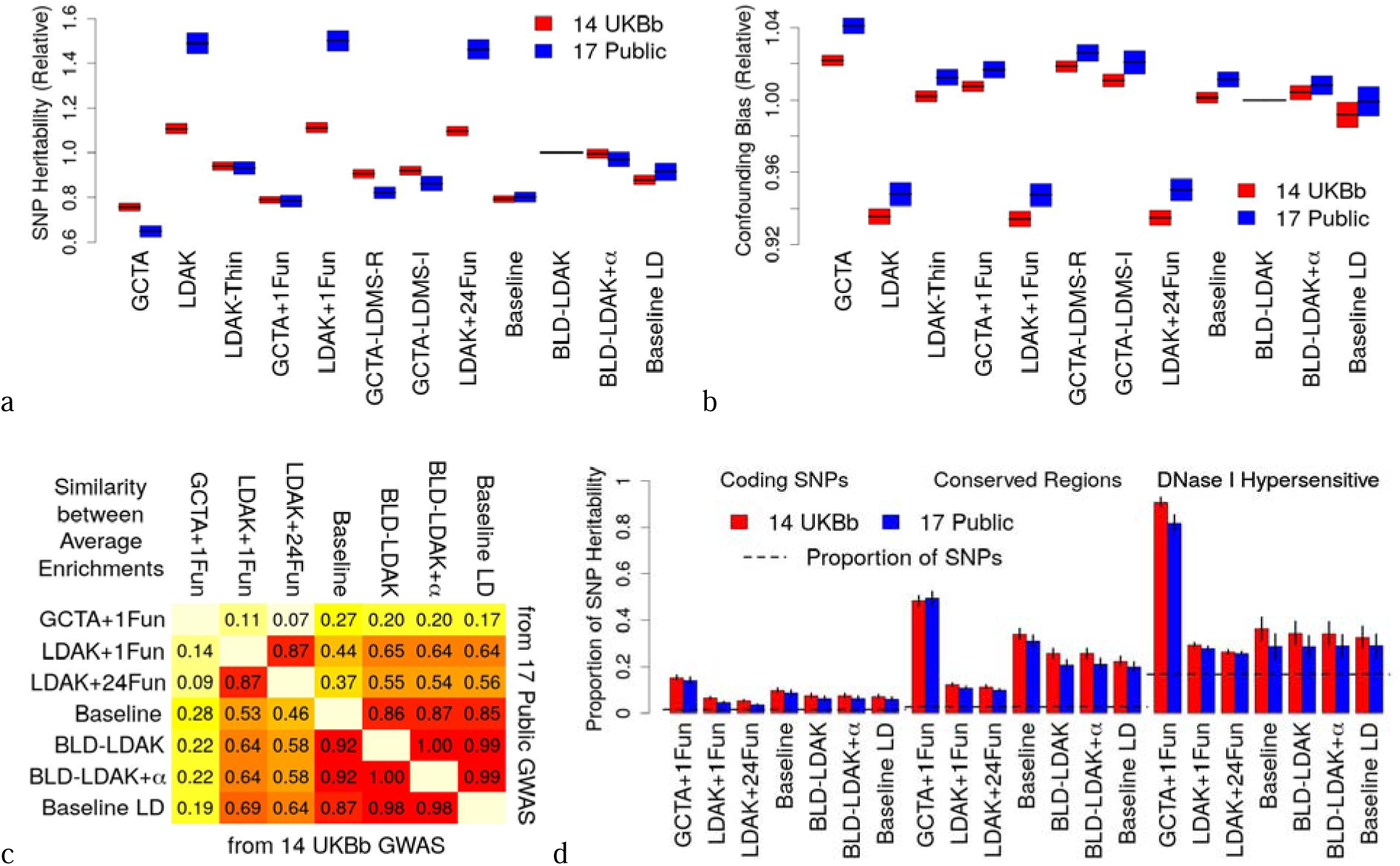
Genetic architecture estimates from different heritability models. **a** & **b**, Average estimates of SNP heritability and confounding bias from twelve heritability models; values are relative to those from the BLD-LDAK Model, and calculated using either the 14 UKBb or 17 Public GWAS (bar heights indicate 95% confidence intervals). There are 24 versions of the GCTA+1Fun and LDAK+1Fun Models (one for each function indicator); values here correspond to the versions with highest average logl_SS_. **c**, Concordance correlation coefficient between average estimates of functional enrichments from seven heritability models; values are calculated using either the 14 UKBb or 17 Public GWAS. **d**, Average estimates of the proportion of SNP heritability contributed by coding SNPs, conserved regions and DNase I hypersensitive sites (three of the 24 functional categories of SNPs) from seven heritability models; values are calculated using either the 14 UKBb or 17 Public GWAS (vertical segments indicate 95% confidence intervals). The estimates of functional enrichments are obtained by dividing these estimates by the proportion of SNPs in each category (dashed horizontal lines).

### Evidence of selection

Figure 2a and Supplementary Table 7 report estimates of α in the BLD-LDAK+Alpha Model. This parameter specifies the assumed relationship between heritability and MAF,^1,4^ and has been used to measure selection^10,18^ (negative α indicates that less-common SNPs tend to have larger effect sizes than more-common SNPs, and vice versa). Across the 31 GWAS, the average estimate of α is -0.25 (s.d. 0.01). Estimates of α are significantly negative (P<0.05/31) for seven of the 14 UKBb GWAS and three of the 17 Public GWAS; the three most significant are for height (α=-0.35, s.d. 0.03), college education (α=-0.43, s.d. 0.05) and systolic blood pressure (α=-0.33, s.d. 0.05).

**Fig. 2.**
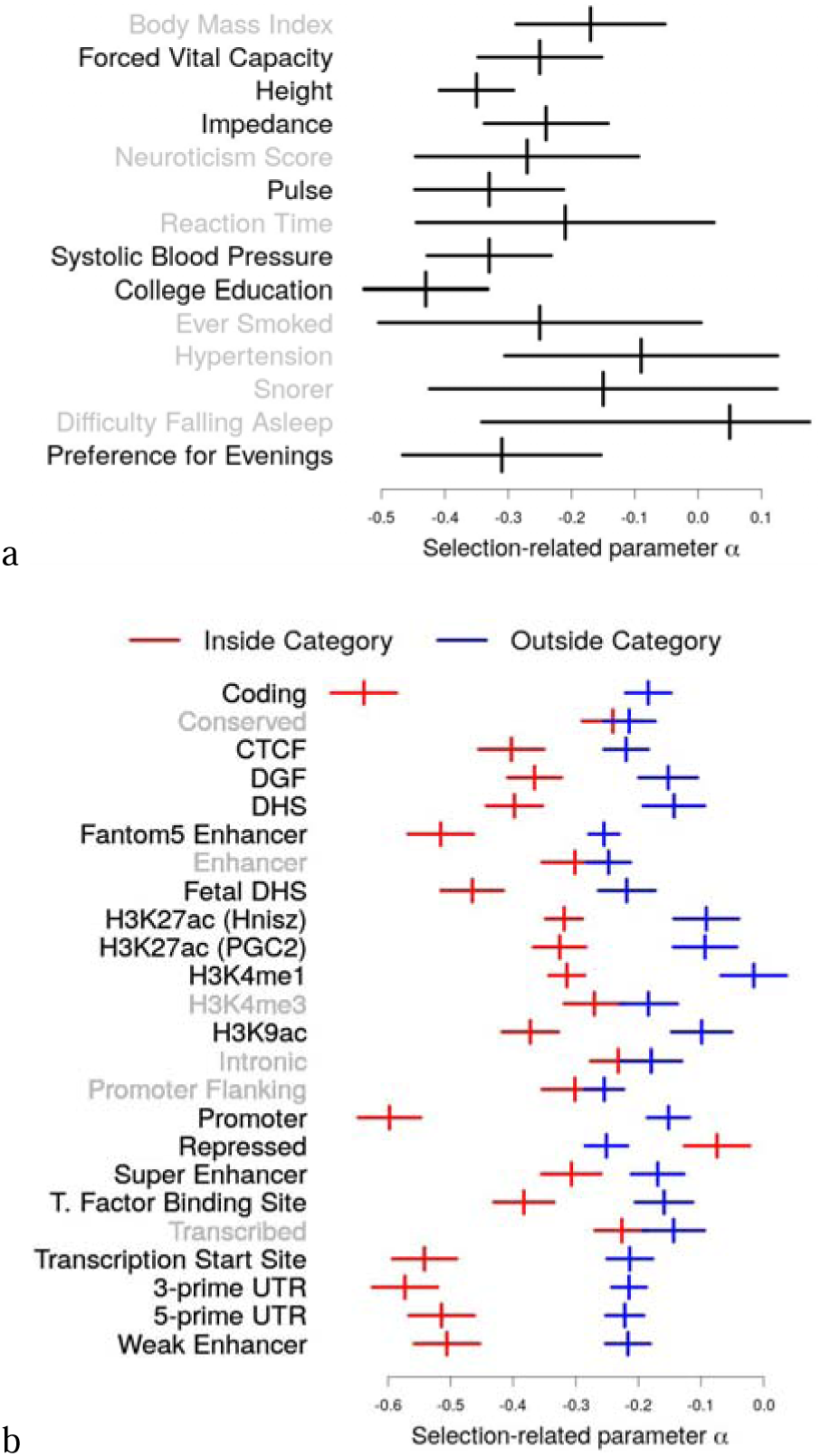
Evidence of selection. **a**, Estimates of α for the 14 UKBb GWAS, obtained using the BLD-LDAK+Alpha Model (horizontal lines indicate 95% confidence intervals). α is significantly negative (P<0.05/31) for the seven **bold** traits. **b**, Estimates of α for SNPs inside and outside of the 24 functional categories, obtained using the BLD-LDAK-Lite+Alpha Model and averaged across the 14 UKBb GWAS (horizontal lines indicate 95% confidence intervals). α varies significantly for the 18 **bold** categories.

We also investigate whether α varies across functional categories of SNPs. For computational reasons, we use the seven-parameter BLD-LDAK-Lite+Alpha Model, a reduced version of the BLD-LDAK+Alpha Model (Supplementary Table 8 explains how we constructed the BLD-LDAK-Lite+Alpha Model by identifying the most important annotations of BLD-LDAK+Alpha Model, while Supplementary Fig. 4 shows that estimates of α from the BLD-LDAK-Lite+Alpha Model are consistent with those from the BLD-LDAK+Alpha Model). Figure 2b and Supplementary Table 9 show that α differs significantly (P<0.05/24) for 18 of the 24 functional categories. The largest differences in α are observed for coding SNPs, promoter regions, 3’UTR and transcription start sites (inside each of these four categories, the average estimate of α is below -0.5).

## Discussion

When software for estimating SNP heritability were first developed, little attention was given to the heritability model, and it was standard to assume that all SNPs are expected to contribute equal heritability.^2,3^ It is now recognized that this assumption is sub-optimal,^1,4^ and the best way to model how heritability varies across the genome has become a topic of debate.^7,8,10,19,20^ Heritability models have previously been compared based on REML likelihood,^1,20^ prediction accuracy^4,20^ and performance on simulated data,^7,8^ however, all three approaches have shortcomings; the REML likelihood requires individual-level data and can not be computed for complex heritability models, to measure prediction accuracy requires two independent datasets for each trait (one for training and one for testing) and there is no consensus regarding the best prediction method, while comparisons of heritability models based on simulated data are sensitive to the assumptions of the simulation model.^1^

We have proposed logl_SS_, an approximate model likelihood that can be computed from summary statistics and for complex heritability models. logl_SS_ can be used both to evaluate heritability models (e.g., if two models produce contrasting estimates, then those from the model with highest AIC should be preferred), and to improve them (for example, by combining pairs of models when this significantly improves logl_SS_). Using logl_SS_, we have shown that the Baseline LD Model is the best of the existing heritability models, but that it can be substantially improved by incorporating the SNP weightings and MAF scaling used by the LDAK Model. Estimates of confounding bias and functional enrichments from the resulting BLD-LDAK Model are close to those from the Baseline LD Model, while its estimates of SNP heritability are between those of the Baseline LD, GCTA-LDMS-I and LDAK Models.

Our results support the conclusion of Gazal et al.,^20^ that estimates of functional enrichments from the Baseline LD Model are more accurate than those from the LDAK+1Fun Model. They provide partial support for Evans et al.,^8^ who argued that the GCTA-LDMS-I Model produces the most accurate estimates of SNP heritability (although we found that the GCTA-LDMS-I Model performs well compared to the other existing models, our analysis indicates that the BLD-LDAK Model should now be preferred). Our results support our previous finding^4^ that the LDAK Model is more realistic than the GCTA Model, but not that estimates of functional enrichments from the LDAK+24Fun Model should be preferred to those from the Baseline and Baseline LD Models. We discuss these three papers in detail in the Supplementary Note.

Recently, Hou et al.^21^ proposed GRE, a method for estimating SNP heritability without specifying a heritability model. While we agree with the benefits of performing heritability analysis without requiring a heritability model, this is not feasible for most analyses. For example, GRE can not be used on our Public GWAS, because it requires individual-level data, nor can it be used on our (full) UKBb GWAS, because it requires that the number of samples is larger than the number of SNPs on the largest chromosome. For Supplementary Fig. 5, we run GRE using a subset of our UKBb GWAS (we restrict to 623,000 directly-genotyped SNPs). Estimates of SNP heritability from the BLD-LDAK Model are consistent (P>0.05/14) with those from GRE for all 14 traits. However, this partly reflects that reducing the number of SNPs reduces the impact of the heritability model (the LDAK-Thin, GCTA-LDMS-R, GCTA-LDMS-I and Baseline LD Model also produce consistent estimates for all 14 traits).

The BLD-LDAK+Alpha Model is a generalization of the BLD-LDAK Model. The two models have similar fit and produce similar estimates of SNP heritability, confounding bias and functional enrichments. For computational reasons, we generally recommend the BLD-LDAK Model. However, the advantage of the BLD-LDAK+Alpha is that it provides estimates of the selection-related parameter α. Our results broadly agree with those of Zeng et al.;^18^ using the software BayesS, they found significantly negative α for 23 out of 28 UK Biobank traits, while their average estimate of α was -0.38 (s.d. 0.01). To our knowledge, SumHer is the only software to estimate α from summary statistics, and therefore can be viewed as a more computationally-efficient alternative to BayesS (for a full comparison of the two methods and their results, see Supplementary Fig. 6).

Although its shortcomings have been well documented, the GCTA Model continues to be widely used in statistical genetics. It remains the default model of both the GCTA and LDSC software, and is the model used by LD Hub.^22^ More widely, the GCTA Model is implicitly assumed by any penalized or Bayesian regression method that standardizes genotypes then assigns the same penalty or prior distribution to each SNP, or in simulations when causal SNPs are picked at random then their standardized effects sizes drawn from the same distribution. Ideally, the GCTA Model should be replaced by the BLD-LDAK or BLD-LDAK+Alpha Model whenever it occurs. However, we recognize that for many methods, introducing a multi-parameter heritability model would require substantial algorithmic changes and dramatically increase computational demands. When this is the case, we instead recommend using the LDAK-Thin Model; this is a one-parameter model, so computation demands should not be affected, that can be incorporated in any existing method simply by changing which predictors are included in the regression and how these are standardized.

We finish by highlighting three areas for future work. Firstly, we have only considered common SNPs; with the increasing availability of sequence data, it will be necessary to examine whether the BLD-LDAK and BLD-LDAK+Alpha Models remain the best performing model when rare SNPs are included. Secondly, the ability to measure model fit for very large sample sizes means that we now have sufficient power to construct heritability models specific to either individual traits or groups of traits (Supplementary Table 2). Thirdly, we have only considered the genomic annotations contained within the Baseline LD Model. We expect it will be possible to find new annotations predictive of how heritability varies across the genome (i.e., whose inclusion in the heritability model significantly increases logl_SS_). Identifying these will both improve the performance of the heritability model and our understanding of the genetic architecture of complex traits.

## Online Methods

Let h^2^_j_ denote the heritability contributed (uniquely) by SNP j. Suppose we have summary statistics from a GWAS of n individuals; let S_j_ denote the χ^2^(1) test statistic from regressing the phenotype on SNP j. Suppose also we have access to an ancestrally-matched reference panel, from which we can estimate r^2^_jl_, the squared correlation between SNPs j and l (genotypes coded additively).

### Linear heritability models

The heritability model describes how the expectation of h^2^_j_ varies across the genome. We first assume the model takes the form

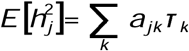

where the a_jk_ are pre-specified SNP annotations and the parameters τ_k_ are estimated in the analysis. Assuming no confounding bias^3^

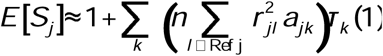

where Ref j indexes the reference panel SNPs ‘near’ SNP j (a working definition of near is within 1cM^4^). The term within the parentheses is known, and thus we can estimate the τ_k_ by regressing S_j_ on E[S_j_] (details below). To allow for confounding bias, the authors of LDSC^3^ recommend increasing each E[S_j_] by A, the average amount each test statistic is inflated additively (it is standard to then report 1+A, referred to as the intercept), while we^4^ instead recommended scaling each E[S_j_] by C, the average amount each test statistic is inflated multiplicatively. Allowing for confounding bias (whether additive or multiplicative) results in a revised form for Eq. (1), but it remains that E[S_j_] can be expressed as a linear combination of the model parameters^4^ (now the τ_k_ plus either A or C), and so we can continue to estimate the parameters by regressing S_j_ on E[S_j_].

### Approximate model likelihood

The authors of LDSC^3^ derived that each S_j_ approximately follows a scaled χ^2^(1) distribution with scale factor E[S_j_] (or equivalently, a gamma distribution with shape ½ and scale 2E[S_j_]). Let

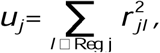

where Reg j indexes the regression SNPs (those used when regressing S_j_ on E[S_j_]) near SNP j. We propose the approximate log likelihood

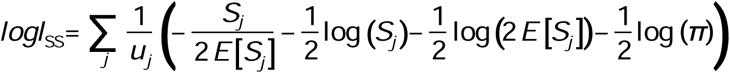

The term within the large parentheses is the log likelihood for a single SNP; therefore logl_SS_ computes a weighted sum of these, where the weights 1/u_j_ reflect local correlations. Supplementary Figs. 1 & 2 show that logl_SS_ is concordant with the exact likelihood computed from REML and can be used for likelihood ratio testing.

### Estimating parameters

LDSC estimates the parameters of the heritability model using weighted least-squares regression,^3^ with regression weights 1/u_j_ x 1/(2E[S_j_]^2^); weighting by 1/u_j_ allows for correlations between nearby SNPs (motivating our use of 1/u_j_ in the definition of logl_SS_), while weighting by 1/(2E[S_j_]^2^) allows for heteroscedasticity (2E[S_j_]^2^ is the variance of a gamma distribution with shape ½ and scale 2E[S_j_]).

When we proposed SumHer, we followed LDSC and estimated parameters using weighted least-squares regression.^4^ However, now that we have an expression for the model likelihood, we can instead use maximum likelihood estimation. To identify the values that maximize logl_SS_, we use (multi-dimensional) Newton-Raphson.^23^ Let θ denote the vector of parameters (the τ_k_ and, if allowing for confounding, either A or C). Starting from the null model (τ_k_=0, A=0, C=1), we update θ iteratively until convergence using θ→θ-E^-1^D, where the vector D and matrix E contain, respectively, the first and second derivatives of logl_SS_ evaluated using the current parameter values (the required derivatives can be computed using the chain rule; for example, δlogl_SS_ /δθ_k_= δlogl_SS_ /δE[S_j_] x δE[S_j_]/δθ_k_). Sometimes, a move causes a (non-negligible) reduction in logl_SS_. When this happens, we cancel the move, then for the next iteration (only) update each parameter once individually using (one-dimensional) Newton-Raphson.

Except where stated, our analyses use the maximum likelihood solver, regardless of the heritability model. In Supplementary Fig. 7, we compare the impact of changing the solver. This shows that for simple heritability models, the weighted least-squares and maximum likelihood solver result in identical logl_SS_, but that for complex models, the maximum likelihood solver often results in substantially higher logl_SS_

### Non-linear heritability models

To date LDSC and SumHer have required that the heritability model is linear (this ensures that E[S_j_] can be expressed as a linear combination of the model parameters). However, SumHer can now accommodate a small number of non-linear parameters. We first crudely estimate the non-linear parameters using a grid-search, selecting the values that result in highest logl_SS_. We then increase resolution and obtain standard deviations by fitting a Gaussian likelihood to the realizations of logl_SS_. Supplementary Fig. 8 demonstrates how we use this approach to estimate the selection-related parameter α.

### Leave-one-chromosome-out prediction of test statistics

For each SNP we compute E[S_j_] in Eq. (1) using parameter estimates obtained from the other 21 chromosomes. In Table 1 we report a weighted correlation between predicted and observed test statistics

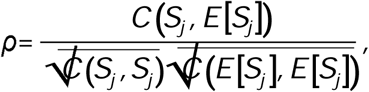

where

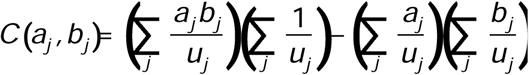

We estimated the standard deviation of ρ using block jackknifing with 200 blocks.^3,24^ We consider it appropriate to include the weights 1/u_j_, as otherwise ρ will overweight high-LD regions, however, Supplementary Table 1 shows that the ranking of models is the same if we instead compare unweighted correlations.

### GWAS

We accessed UK Biobank^13,14^ (UKBb) data via Project 21432. In total we identified 20 phenotypes that were recorded for the majority of individuals: the 14 we retained were body mass index (data field 21001), forced vital capacity (3062), height (50), impedance (23106), neuroticism score (20127), pulse rate (102), reaction time (20023), systolic blood pressure (4080), college education (6138), ever smoked (20160), hypertension (20002), snorer (1210), difficulty falling asleep (1200) and preference for evenings (1180); the six we discarded were asthma, wears glasses, handedness, any mouth problem, basal metabolic rate and diastolic blood pressure (each either had estimated heritability less than 0.1 or was highly correlated with one of the retained phenotypes). The imputed dataset contains 487k individuals recorded for 93M SNPs. However, after quality control, which included filtering individuals based on ancestry and relatedness (for the latter, we ensured that no pair remained with allelic correlation >0.02), and excluding SNPs with MAF<0.01, info score <0.99 or within the major histocompatibility complex (Chr6:25-34Mb), only 130,080 individuals and 4,725,151 SNPs remained.^4,5,25^ As we have individual-level data, we could use our previously-published protocols to confirm that confounding due to residual population structure, relatedness or genotyping errors was slight^1^. For the association analysis, we tested each SNP using linear regression (regardless of whether the phenotype was continuous, categorical or binary), having first regressed the phenotype on 13 covariates: age (data field 21022), sex (31), Townsend Deprivation Index (189) and ten principal components. For more details, see Supplementary Figs. 9 & 10.

The 17 Public GWAS are coronary artery disease,^26^ Crohn’s Disease,^27^ ever smoked,^28^ inflammatory bowel disease,^27^ rheumatoid arthritis,^29^ schizophrenia,^30^ type 2 diabetes,^31^ bone mineral density,^32^ body mass index,^33^ depressive symptoms,^34^ height,^35^ menarche age,^36^ menopause age,^37^ neuroticism,^34^ subjective well-being,^34^ waist-hip ratio^38^ and years education.^39^ These are a subset of the 24 GWAS we considered previously;^4^ we excluded the remaining seven GWAS as the authors of LDSC^9,40^ recommend only using traits with a heritability Z-score above seven. For these GWAS we have to rely on the quality control choices of the original authors (Supplementary Table 10), which are generally less strict than ours, and without access to individual-level data, we could not test for confounding due to population structure, relatedness or genotyping errors.

### Software settings

When running an analysis using LDSC or SumHer it is necessary to choose the heritability and confounding models, provide a reference panel, select the regression and heritability SNPs, and if estimating enrichments, specify the expected proportion of SNP heritability contributed by each category (for an explanation of each option, see Supplementary Table 11). We describe the different heritability models we consider below. For the other options, our main analysis follows the recommendations of LDSC.^3,9^ When analyzing the UKBb GWAS, we assume there is no confounding bias (the exception is for Figure 1b, when we allow for additive confounding bias, then report the estimate of 1+A); when analyzing the Public GWAS, we always allow for additive confounding bias. Our reference panel is the 1000 Genome Project^16^ dataset provided on the LDSC website (see URLs), which contains 489 European individuals recorded for 10.0M autosomal SNPs with MAF>0.005. When analyzing the UKBb GWAS, the regression SNPs are all 4.7M GWAS SNPs; when analyzing the Public GWAS, the regression SNPs are the GWAS SNPs present in HapMap3^41^ but not in the major histocompatibility complex (on average 1.1M SNPs per GWAS). The heritability SNPs are the 6.0M reference panel SNPs with MAF≥0.05. When computing estimates of enrichments, we divide the estimated proportion of SNP heritability contributed by a category by the proportion of SNPs it contains.

### Sensitivity analyses

Supplementary Table 1 shows that the ranking of heritability models is the same if we use a UK Biobank^13,14^ reference panel (instead of the 1000 Genomes Project panel), if we reduce the reference panel to the 4.7M SNPs in our UKBb GWAS, or if we reduce the regression SNPs to a subset in approximate linkage equilibrium. Supplementary Table 12 shows that enrichment estimates from the BLD-LDAK Model are similar if the heritability SNPs are all 10.0M reference panel SNPs (rather than just the 6.0M with MAF≥0.05).

### Heritability models

Full details of all heritability models are provided in Supplementary Tables 13 & 14. The one-parameter GCTA Model^2^ (the default model of both the GCTA and LDSC software) assumes E[h^2^_j_]=τ_1_. The GCTA-LDMS-R^7^ and GCTA-LDMS-I^8^ Models both take the form

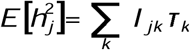

where I_jk_ indicates whether SNP j is in Bin k. The bins are obtained by first dividing the genome four ways based on LD, then M ways based on MAF, (when dividing based on LD, the GCTA-LDMS-R Model ranks SNPs based on regional LD scores, while the GCTA-LDMS-I Model ranks based on per-SNP LD scores). We opt for M=5, with the boundaries at 0.1, 0.2, 0.3 and 0.4 (in total 20 bins); this choice is based on the first application of GCTA-LDMS-R,^7^ which used seven MAF tranches with boundaries at 0.001, 0.01, 0.1, 0.2, 0.3 and 0.4 (we exclude the bottom two tranches as we only consider common SNPs). Supplementary Table 15 shows that this version performs better than using M=2, with the boundary at 0.05 (in total 8 bins), a choice based on the first application of GCTA-LDMS-I,^8^ which used four MAF tranches with boundaries at 0.0025, 0.01 and 0.05.

The 53-parameter Baseline Model^9^ adds to the GCTA Model 52 functional annotations; 24 of these are function indicators (e.g., indicating SNPs within coding regions), while 28 are ‘buffer’ indicators (e.g., indicating SNPs within 500bp of a coding region). The 75-parameter Baseline LD Model^10^ adds to the GCTA Model 74 SNP annotations: six are LD-related annotations (e.g., estimated allele age), ten are MAF indicators (e.g., indicating SNPs with 0.05 ≤ MAF < 0.07) and 58 are functional annotations (including the 52 used in the Baseline Model). The two-parameter GCTA+1Fun Model^12^ adds to the GCTA Model one function indicator from the Baseline Model (there are 24 versions of this model, one for each indicator).

The one-parameter LDAK Model^1,4^ assumes E[h^2^_j_]=w_j_p_j_^0.75^τ_1_, where w_j_ is the LDAK weighting of SNP j, f_j_ is its MAF and p_j_=f_j_(1-f_j_). We recommend that the weightings are only computed over high-quality SNPs^1,6^ (so low- and moderate-quality SNPs automatically get w_j_=0). We do not have SNP info scores for the 1000 Genome Project reference panel, so when computing weightings, we restrict to the 4.7M SNPs in our UKBb GWAS (i.e., we assume that SNPs well-genotyped in the UK Biobank are well-genotyped in the 1000 Genome Project). The two-parameter LDAK+1Fun Model^1^ assumes E[h^2^_j_]=w_j_p_j_^0.75^(τ_1_+b_ji_τ_2_), where b_ji_ is the ith function indicator from the Baseline LD Model (there are 24 versions of this model, one for each indicator), while the 25-parameter LDAK+24Fun Model^1^ assumes

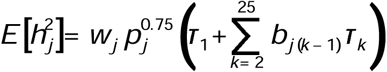

The novel 66-parameter BLD-LDAK and 67-parameter BLD-LDAK+Alpha Model both take the form

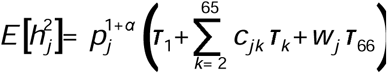

where the c_jk_ are the 64 LD-related and functional annotations from the Baseline LD Model;^10^ the BLD-LDAK Model fixes α=-0.25, while the BLD-LDAK+Alpha estimates α from the data. To construct the BLD-LDAK Model we first added the LDAK weighting to the Baseline LD Model (this increased average logl_SS_ by 11), then scaled all annotations by p_j_^0.75^ (this increased average logl_SS_ by a further 50). At this point we noted that the 10 MAF indicators had limited value (excluding them reduced average logl_SS_ by only 8) so we removed them. We were unable to improve the model further by adding features from the GCTA-LDMS-R and GCTA-LDMS-I Models (for example, if we incorporated the 4 LD or the 20 MAF-LD bins from the GCTA-LDMS-I Model, this increased the number of parameters by 3 and 19, respectively, but increased average logl_SS_ by only 2 and 13). For more details, see Supplementary Table 16.

The seven-parameter BLD-LDAK-Lite and eight-parameter BLD-LDAK-Lite+Alpha Models are reduced versions of the BLD-LDAK and BLD-LDAK+Alpha Models, respectively, obtained by removing two of the nine continuous annotations and all 57 binary annotations (Supplementary Table 8 explains how we used forward stepwise selection to decide which of the continuous annotations to retain). Supplementary Fig. 4 shows that estimates of SNP heritability and confounding bias from the BLD-LDAK-Lite Model are consistent with those from the BLD-LDAK Model, and that estimates of α from the BLD-LDAK-Lite+Alpha Model are consistent with those from the BLD-LDAK+Alpha Model. To allow α to vary based on functional annotations, we use a 16-parameter model obtained by concatenating two versions of the BLD-LDAK-Lite+Alpha Model (one where only SNPs within the category contribute heritability, and one where only SNPs outside the category contribute).

When computing the LDAK weightings, the first step is to thin SNPs so that no pair remains within 100kb with r^2^_jl_>0.98 (excluding duplicate SNPs substantially improves the efficiency of the solver used to compute the weightings^1^). The LDAK-Thin Model assumes E[h^2^_j_]=I_j_ p_j_^0.75^τ_1_, where I_j_ indicates whether SNP j remains after the thinning. To implement the LDAK-Thin Model within an existing penalized or Bayesian regression method requires two changes: firstly, thin the SNPs (for the UKBb data, this reduced the number from 4.7M to 1.4M); secondly, center and scale the genotypes so that SNP j has variance p_j_^0.75^.

### Computational demands

To run SumHer with a linear heritability model involves two steps.^4^ The first is to compute the tagging file; this takes at most one day on a single CPU and requires 20Gb memory. The second is to estimate the parameters of the heritability model; this takes at most two hours and requires 20Gb memory. To run SumHer with a non-linear heritability model, it is necessary to repeat the above process multiple times (e.g., Supplementary Fig. 8 explains how we fit the BLD-LDAK+Alpha Model by calculating tagging files and estimating parameters for 31 versions of BLD-LDAK Model). The LDAK website (see URLs) provides pre-computed tagging files for the BLD-LDAK and BLD-LDAK-Lite+Alpha Models, suitable for analyzing European, South Asian, East Asian or African populations.

## URLs

LDAK, http://www.ldak.org; LDSC, http://www.github.com/bulik/ldsc; UK Biobank https://www.ukbiobank.ac.uk

## Acknowledgements

We thank B. Shaban for help with the LDAK website, and A. Price, S. Gazal and H. Finucane for helpful discussions. D.S. is funded by the European Union’s Horizon 2020 Research and Innovation Programme under the Marie Skłodowska-Curie grant agreement no. 754513, by Aarhus University Research Foundation (AUFF) and by the Independent Research Fund Denmark under Project no. 7025-00094B. D.S. and D.J.B. are funded by the Australian Research Council under Project no. DP190103188.

## Author contributions

D.S. and J.H. performed the analyses, D.S. and D.J.B. wrote the manuscript.

## Data availability

We performed the UKBb GWAS using data applied for and downloaded via the UK Biobank website (see URLs). We obtained summary statistics for the Public GWAS from the websites of the corresponding studies. We downloaded the 1000 Genome Project data from the LDSC website (see URLs).

