## Supplementary Material for "Evaluating and improving heritability models using summary statistics"

**Supplementary Note 1: Previous comparisons of heritability models**

**Supplementary Note 2: Guidelines for using $logl_{\text{SS}}$ to evaluate and improve heritability models**

**Supplementary Figure 1: Comparison of likelihoods**

**Supplementary Figure 2: Likelihood ratio testing of heritability models**

**Supplementary Figure 3: Estimated proportions of SNP heritability**

**Supplementary Figure 4: Reduced-complexity heritability models**

**Supplementary Figure 5: Comparison with GRE**

**Supplementary Figure 6: Comparison with BayesS**

**Supplementary Figure 7: Comparison of weighted least-squares and maximum likelihood solvers**

**Supplementary Figure 8: Estimating $\alpha$**

**Supplementary Figure 9: UKBb GWAS**

**Supplementary Figure 10: Reduced quality control for UKBb GWAS**

**Supplementary Table 1: Average $logl_{\text{SS}}$ for main heritability models**

**Supplementary Table 2: $logl_{\text{SS}}$ for individual GWAS**

**Supplementary Table 3: Estimates of SNP heritability**

**Supplementary Table 4: Estimates of confounding bias**

**Supplementary Table 5: Estimates of average proportions of SNP heritability and enrichments from 14 UKBb GWAS**

**Supplementary Table 6: Estimates of average proportions of SNP heritability and enrichments from 17 Public GWAS**

**Supplementary Table 7: Genome-wide estimates of $\alpha$**

**Supplementary Table 8: Constructing the BLD-LDAK-Lite Model**

**Supplementary Table 9: Estimates of $\alpha$ across functional categories**

**Supplementary Table 10: Details of the 31 GWAS**

**Supplementary Table 11: Software options**

**Supplementary Table 12: Sensitivity of enrichment estimates to the choice of heritability SNPs**

**Supplementary Table 13: Heritability models**

**Supplementary Table 14: Baseline LD Model SNP annotations**

**Supplementary Table 15: Average $logl_{\text{SS}}$ for alternative heritability models**

**Supplementary Table 16: Constructing the BLD-LDAK Model**

**Supplementary Note 1: Previous comparisons of heritability models.** Here we compare our results with those of three previous papers. In summary, while we find errors in the work of Gazal *et al.*,[1] our results support their overall finding that estimates of functional enrichments from the Baseline LD Model are more accurate than those from the LDAK+1Fun Model. Evans *et al.*[2] argued that when estimating SNP heritability, the GCTA-LDMS-I Model performs better than the LDAK Model; our results indicate that these two models perform approximately the same, but that now the BLD-LDAK Model should be preferred. Our results support our earlier finding[3] that the LDAK Model is more realistic than the GCTA Model, but not that estimates of enrichment from the the LDAK+24Fun Model are more accurate than those from the Baseline and Baseline LD Models. For definitions of the different heritability models discussed, see Supplementary Table 13.

In their paper **"Reconciling S-LDSC and LDAK functional enrichment estimates,"** Gazal *et al.*[1] compared the Baseline LD and LDAK Models on UK Biobank data in three ways: by computing an approximate REML likelihood, using a hybrid model, and based on prediction performance. Our results support their overall conclusion, that the Baseline LD Model fits better than the LDAK and LDAK+1Fun Model. We agree with their use of a hybrid model; they showed that enrichment estimates from a model which combined the Baseline LD and LDAK+1Fun Models, were closer to those from the Baseline LD Model than those from the LDAK+1Fun Model (a minor criticism is that, in our view, it would have been more appropriate for the hybrid model to combine the Baseline LD and LDAK+24LDAK Models, but this would not have changed the overall conclusion). However we consider their other two analyses to be flawed. Specifically, the REML likelihoods and prediction accuracies they report for the Baseline LD Model in fact correspond to a truncated version of the model obtained by replacing negative estimates of $h^2_j$ with zero. This is not a trivial difference. For example, the table below shows that across the 14 UKBb GWAS, replacing negative estimates of $h^2_j$ with zero causes estimates of SNP heritability to increase on average by 37% (range 15% to 68%).

| | **6.0 M SNPs with MAF>0.05** | | | | **All 10.0 M SNPs** | | | |
|---|---|---|---|---|---|---|---|---|
| **Trait** | $h^2_-$ | $h^2_+$ | $h^2_{\text{SNP}}$ **(SD)** | $h^2_+/h^2_{\text{SNP}}$ | $h^2_-$ | $h^2_+$ | $h^2_{\text{SNP}}$ **(SD)** | $h^2_+/h^2_{\text{SNP}}$ |
| Body Mass Index | -0.05 | 0.31 | 0.27 (0.01) | 1.17 | -0.11 | 0.45 | 0.34 (0.01) | 1.31 |
| Forced Vital Capacity | -0.05 | 0.32 | 0.28 (0.01) | 1.17 | -0.12 | 0.48 | 0.37 (0.01) | 1.33 |
| Height | -0.11 | 0.64 | 0.53 (0.02) | 1.21 | -0.25 | 0.98 | 0.72 (0.03) | 1.35 |
| Impedance | -0.04 | 0.34 | 0.30 (0.01) | 1.15 | -0.12 | 0.49 | 0.38 (0.01) | 1.31 |
| Neuroticism Score | -0.04 | 0.15 | 0.12 (0.01) | 1.32 | -0.08 | 0.22 | 0.14 (0.01) | 1.56 |
| Pulse | -0.06 | 0.23 | 0.17 (0.01) | 1.37 | -0.14 | 0.36 | 0.21 (0.01) | 1.68 |
| Reaction Time | -0.03 | 0.13 | 0.09 (0.00) | 1.32 | -0.06 | 0.18 | 0.12 (0.01) | 1.56 |
| Systolic Blood Pressure | -0.04 | 0.20 | 0.17 (0.01) | 1.22 | -0.09 | 0.30 | 0.21 (0.01) | 1.43 |
| College Education | -0.04 | 0.24 | 0.20 (0.01) | 1.19 | -0.08 | 0.35 | 0.27 (0.01) | 1.29 |
| Ever Smoked | -0.03 | 0.11 | 0.08 (0.00) | 1.39 | -0.06 | 0.17 | 0.10 (0.01) | 1.62 |
| Hypertension | -0.04 | 0.16 | 0.12 (0.01) | 1.30 | -0.08 | 0.23 | 0.15 (0.01) | 1.58 |
| Snorer | -0.04 | 0.11 | 0.07 (0.00) | 1.56 | -0.08 | 0.16 | 0.08 (0.01) | 2.02 |
| Difficulty Falling Asleep | -0.05 | 0.11 | 0.07 (0.00) | 1.68 | -0.09 | 0.17 | 0.08 (0.01) | 2.06 |
| Preference for Evenings | -0.03 | 0.17 | 0.13 (0.01) | 1.26 | -0.08 | 0.25 | 0.17 (0.01) | 1.45 |
| **Average** | **-0.04** | **0.17** | **0.13 (0.00)** | **1.37** | **-0.08** | **0.25** | **0.16 (0.00)** | **1.63** |

***Truncating the Baseline LD Model.*** *We analyze the 14 UKBb GWAS using the LDSC software assuming the Baseline LD Model.* $h^2_-$ *and* $h^2_+$ *are, respectively, the sums of negative and positive* $\hat{h}^2_j$ *(the estimated heritability contributed by SNP* $j$*). Without truncating negative* $\hat{h}^2_j$*, the estimate of SNP heritability is* $h^2_{SNP} = h^2_- + h^2_+$*; with truncation, the estimate is* $h^2_+$*. For the left half of the table, the heritability SNPs (those used when summing* $\hat{h}^2_j$*) are the reference panel SNPs with MAF>0.05 (this is the recommendation of the authors of LDSC); for the right half, they are all reference panel SNPs. It is not clear how to compute the s.d. for estimates of* $h^2_-$ *and* $h^2_+$*, however, we expect that they are similar to those for* $h^2_{SNP}$*.*

Even without this error, our method of comparing heritability models has a number of advantages. While $logl_{\text{SS}}$, our approximate log likelihood, can be computed from summary statistics and for complex heritability models, to compute the REML likelihood requires individual-level data, and is only feasible for relatively small sample sizes and simple heritability models. To overcome these issues, Gazal *et al.* reduced the sample size to 20 000, while for the Baseline LD Model, instead of computing an exact likelihood, they calculated a lower bound (first they used external data to estimate the 75 heritability parameters up to a constant of proportionality, then they used REML to estimate the constant of proportionality). These restrictions substantially reduced the power of their analysis to discriminate between models. For example, if $l$ denotes the average (approximate) REML log likelihood, their first analysis (of 2.8 M SNPs) reported that $l$ for the Baseline LD Model was only 40 higher than $l$ for the LDAK Model; this means that ranked based on $l$ or $l - K/2$, where $K$ is the number of parameters, the Baseline LD Model is best, whereas ranked based on the Akaike Information Criterion ($2K - 2l$), the LDAK Model is best (in our analysis, the average difference between $logl_{\text{SS}}$ for the two models was $\sim$600, so the choice of penalization was irrelevant). To construct prediction models, Gazal *et al.* used LDpred-funct-inf,[4] a version of LDpred[5] that allows the user to specify the expected heritability of each SNP as a prior. For each trait, they trained models using 409 k samples, then tested on 25 k samples. We also compared models based on prediction (Supplementary Table 1), but in our case we measured how accurately we could predict test statistics using those from other chromosomes. The main advantages of our approach is that it is not necessary to select a prediction method (LDpred is one of many hundreds available, with no consensus on which is best), nor to identify independent test and training datasets.

Finally, Gazal *et al.* suggested that the LDAK Model suffers because many of the LDAK weightings are zero, and would be improved by using the non-sparse weightings obtained by adding the option `-quick-weights YES` (we refer to these as the LDAK-Quick weightings). With $logl_{\text{SS}}$, it is straightforward to investigate this suggestion. Supplementary Table 15 shows that replacing the LDAK weightings with the LDAK-Quick weightings does improve model fit (the LDAK-Quick and LDAK-Quick+24Fun Models have average $logl_{\text{SS}}$ 131 and 181 higher than the LDAK and LDAK+24Fun Models, respectively), but that the majority of this improvement is because they effect a model intermediate between the GCTA and LDAK Models (the LDAK-Quick weightings penalize high-LD regions less strongly than the LDAK weightings). We also find that the LDAK-Quick Model outperforms the LDAK-Thin Model (average $logl_{\text{SS}}$ is 23 higher), indicating that it should be preferred when requiring a one-parameter model. However, despite its lower model fit, we have advocated the LDAK-Thin Model owing to its simplicity; to incorporate this model in an existing regression method, it is only necessary to alter the choice of SNPs and change how genotypes are scaled (whereas to incorporate the LDAK-Quick Model, one must additionally compute the LDAK-Quick weightings).

In their paper **“Comparison of methods that use whole genome data to estimate the heritability and genetic architecture of complex traits,"** Evans *et al.*[2] compared various versions of the GCTA and LDAK Models based on how accurately each estimated the SNP heritability of simulated phenotypes. Using whole genome sequence data (8 201 individuals recorded for 38 913 048 SNPs with MAF>0.0003), they generated phenotypes each with heritability 0.5 and 1000 causal SNPs, under a variety of heritability models. Let $\mathbb{E}[h_j^2]$ denote the expected heritability contributed by the $j$th causal SNP; among others, they generated phenotypes under the GCTA Model ($\mathbb{E}[h_j^2] \propto 1$), the LDAK Model ($\mathbb{E}[h_j^2] \propto w_j[f_j(1-f_j)]^{0.75}$) and the Baseline LD Model ($\mathbb{E}[h_j^2] \propto \sum_k c_{jk}\,\tau_k$, using estimates of $\tau_k$ obtained from analyzing 31 complex traits[6]). They additionally varied whether the causal SNPs were picked at random from all available SNPs or from only a subset (e.g., those with MAF<0.0025 or those within DNase I hypersensitive sites). They then analyzed each phenotype assuming, among others, the GCTA, GCTA-LDMS-R, GCTA-LDMS-I and LDAK Models, finding that the GCTA-LDMS-I Model tended to produce estimates of SNP heritability closest to 0.5.

The two key differences between the analysis of Evans *et al.* and ours, are that they included rare SNPs and estimated heritability parameters using REML. Therefore, we perform a similar simulation study, except that we restrict to common SNPs and estimate parameters using SumHer. For this we use 130 080 UK Biobank individuals recorded for 7 558 266 SNPs with MAF>0.01 (this dataset is the same as that described in Supplementary Figure 9, except that we reduced the info score threshold from 0.99 to 0.95 so that the number of SNPs is closer to that of Evans *et al.*). We generate four sets of 100 phenotypes, each with 1000 causal SNPs and heritability 0.5; for the first three sets (which we refer to as GCTA-All, LDAK-All and BLD-LDAK-ALL Phenotypes), we pick the causal SNPs randomly from across the whole genome and sample their effect sizes according to the GCTA, LDAK or BLD-LDAK Models (for the latter, we use estimates of per-SNP heritability from our analysis of height); for the fourth set of phenotypes (which we refer to as GCTA-Rare Phenotypes), we pick the causal SNPs from those with MAF<0.05, and sample their effect sizes according to the GCTA Model. We analyze each phenotype using SumHer, assuming the GCTA, GCTA-LDMS-I, LDAK or BLD-LDAK Model, first using only 8 201 individuals when performing the single-SNP analysis (to match the number used by Evans *et al.*), then using all 130 k individuals (reflecting the typical sample size of large-scale GWAS[7–9]).

The results of our study are shown in the figure below. Like Evans *et al.*, we also find that the GCTA-LDMS-I Model performs well overall; when we analyse 8 201 individuals its average bias is 0.08, reducing to 0.04 when we analyze all 130 k individuals, in both cases approximately half the average bias of the GCTA and LDAK Models. Nonetheless, we do not advocate this method of comparing heritability models. To be confident in the results of a simulation study, we must be confident that the simulated phenotypes are sufficiently similar to real complex traits. However, for this to be possible, we would have to better understand the genetic architecture of complex traits (e.g., the type, number and location of causal variants, and the distribution of their effect sizes). Furthermore, it is difficult to avoid simulations studies being biased in favour of one of the methods. For example, we note that when Evans *et al.* varied the MAF spectrum of causal variants, the four tranches they used (MAF<0.0025, 0.0025<MAF<0.01, 0.01<MAF<0.05 and MAF>0.05) matched the tranches they used in the GCTA-LDMS-I Model, thus giving this model an advantage.

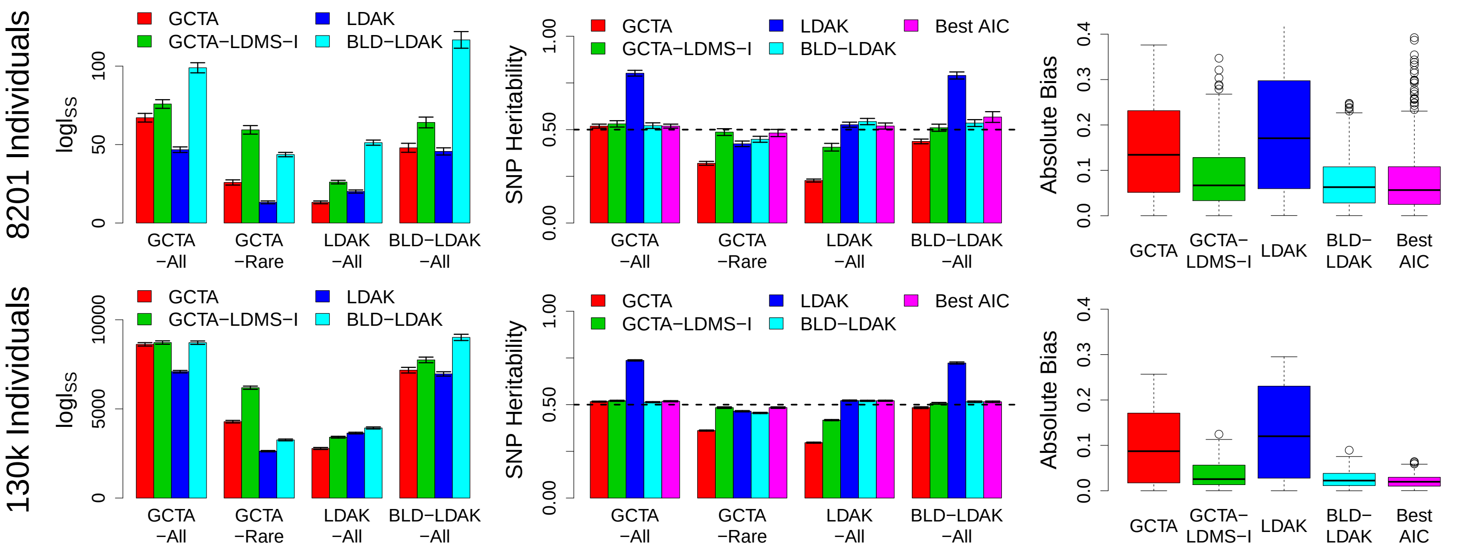


***Comparing heritability models on simulated phenotypes.*** *We generate four sets of 100 phenotypes (each with heritability 0.5): GCTA-All, LDAK-All and BLD-LDAK-All (1000 causal SNPs picked randomly from across the whole genome, with their effect sizes sampled from the GCTA, LDAK and BLD-LDAK Model, respectively) and GCTA-Rare (1000 causal SNPs picked from those with MAF<0.05, with their effect sizes sampled from the GCTA Model). We analyze each phenotype assuming either the GCTA, GCTA-LDMS-I, LDAK or BLD-LDAK Models (1, 20, 1 and 66 parameters, respectively). The left and center plots report average* $logl_{SS}$ *and SNP heritability, respectively, when analysing each set of phenotypes assuming each of the four heritability models (vertical segments indicate 95% confidence intervals). The right plots report the absolute bias of the SNP heritability estimates (difference from 0.5) across all 400 phenotypes; the horizontal lines mark the medians, the boxes indicate the interquartile range (IQR), while the whiskers extend at most 1.5 times the IQR (more extreme values are marked by circles). In the centre and right plots, the purple bars correspond to picking for each phenotype the heritability model with lowest Akaike Information Criterion (equal to* $2K - 2logl_{SS}$, *where* $K$ *is the number of parameters in the heritability model).*

We have previously explained our concerns with comparing heritability models using simulated data, and that we prefer to compare models empirically, based on how well they fit real data.[10] Evans *et al.* acknowledge our view, but in response say that they "observed multiple cases in which negligible differences in log likelihood translated into large differences in bias, as well as situations in which models with higher average log likelihoods produced more biased results than models with lower average log likelihoods." Similar to Evans *et al.*, we observe scenarios where one model fits significantly better than another and yet produces less accurate estimates of SNP heritability. For example, for the GCTA-Rare phenotypes, the GCTA Model fits significantly better than the LDAK Model, yet the average estimate of SNP heritability from the LDAK Model is closer to 0.5. However, in both our simulations and those of Evans *et al.*, it is generally the case that better fitting models produce more accurate estimates of SNP heritability. Finally, we note that were we to compare models based on simulations, then our study indicates that the BLD-LDAK Model should be preferred to the GCTA-LDSM-I Model (its average bias is lower, when using both 8 201 and 130 k individuals), but that it would be better still, for each phenotype, to report the estimate of SNP heritability from whichever of the four heritability models has lowest Akaike Information Criterion.

In our paper **SumHer better estimates the SNP heritability of complex traits,** we analyzed data from two sets of GWAS: 25 Raw GWAS (18 binary traits, 7 quantitative, average sample size 9 700) and 24 Summary GWAS (9 binary traits, 15 quantitative, average sample size 121 000; these include the 17 Public GWAS we use in this paper). We compared the GCTA and LDAK Models in three ways. First we used $logl_{\text{Old}}$, an alternative approximate model likelihood (see below); across the 25 Raw GWAS, $logl_{\text{Old}}$ was on average 17 higher when assuming the LDAK Model, while across the 24 Summary GWAS, it was on average 76 higher. Next we constructed the GCTA+LDAK Model, a concatenation of the GCTA and LDAK Models, where the parameter $p$ indicates the "LDAK proportion" (i.e., the GCTA Model is obtained by setting $p = 0$, while the LDAK Model is obtained by setting $p = 1$); across the 25 Raw GWAS our average estimate of $p$ was 1.03 (s.d. 0.02), while across the 24 Summary GWAS, it was 0.85 (s.d. 0.01). Thirdly, we constructed Bayesian polygenic risk scores (PRS) that incorporated heritability models as prior distributions on effect sizes; across five traits (body mass index, height, HLD & LDL cholesterol and triglyeride levels), the prediction accuracy was on average 5% higher (s.d. 2%) when incorporating the LDAK Model than when incorporating the GCTA Model. Based on these results, we decided not only that the LDAK Model was more realistic than the GCTA Model, but that the latter was redundant (although $p$ was substantially below one for the 24 Summary GWAS, additional analyses indicated that this could be a consequence of genotyping errors).

In this paper, we again found that the LDAK Model outperforms the GCTA Model, although to a lesser extent, and in particular, we no longer found the GCTA Model to be redundant (this is why, for example, the BLD-LDAK and BLD-LDAK+Alpha Models include both the GCTA and LDAK Models). A major reason for this difference is that we did not appreciate the potential benefit of using an extensive reference panel (Supplementary Table 1). Instead, concerned that genotyping errors would lead to biased estimates, we performed strict quality control, of the type recommended when performing REML.[10–15] Furthermore, when analysing each GWAS, we excluded from the reference panel any SNPs not present in the GWAS (equivalent to the approach taken by REML, which assumes that only GWAS SNPs contribute heritability). As a consequence, our reference panel typically contained only 2-4 M SNPs. The table below demonstrates that reducing the size of the reference panel tends to negatively impact the GCTA Model, therefore increasing the advantage of the LDAK Model and resulting in estimates of $p$ closer to one. This occurs not only for the Public GWAS, but also the UKBb GWAS, for which we performed stringent quality control (Supplementary Figure 9), indicating that the relative improvement of the GCTA Model is not an artifact of genotyping errors.

| | | **14 UKBb GWAS** | | | | | | **17 Public GWAS** | | | | | |
|---|---|---|---|---|---|---|---|---|---|---|---|---|---|
| | | **All 10.0 M Reference SNPs** | | | **4.7 M Reference SNPs** | | | **All 10.0 M Reference SNPs** | | | **4.7 M Reference SNPs** | | |
| **Heritability Model** | $K$ | $\overline{logl}_{\text{SS}}$ | $\overline{logl}_{\text{Old}}$ | $\overline{p}$ (s.d.) | $\overline{logl}_{\text{SS}}$ | $\overline{logl}_{\text{Old}}$ | $\overline{p}$ (s.d.) | $\overline{logl}_{\text{SS}}$ | $\overline{logl}_{\text{Old}}$ | $\overline{p}$ (s.d.) | $\overline{logl}_{\text{SS}}$ | $\overline{logl}_{\text{Old}}$ | $\overline{p}$ (s.d.) |
| GCTA Model | 1 | 2104 | 962 | | 1942 | 756 | | 480 | 202 | | 258 | 90 | |
| LDAK Model | 1 | 2160 | 1125 | | 2160 | 1125 | | 517 | 291 | | 329 | 174 | |
| GCTA+LDAK Model | 2 | 2244 | 1180 | 0.65 (0.01) | 2231 | 1173 | 0.80 (0.01) | 588 | 305 | 0.74 (0.01) | 367 | 175 | 0.87 (0.01) |

***Comparing the GCTA and LDAK Models.*** *We report $\overline{logl}_{SS}$ and $\overline{logl}_{Old}$, the improvement in $logl_{SS}$ and $logl_{Old}$ relative to the null model, averaged across either the 14 UKBb or 17 Public GWAS. For the GCTA+LDAK Model (constructed by concatenating the GCTA and LDAK Models), we also report the average estimate of $p$, the LDAK proportion. First we use the full reference panel (10.0 M SNPs), then we reduce to the 4.7 M SNPs in the UKBb GWAS (note that changing the reference panel does not affect the LDAK Model because this always assumes that only the 4.7 M SNPs contribute heritability). We are interested in how reducing the reference panel affects the relative performance of the GCTA and LDAK Models (i.e., changes the differences between their model fits or the estimate of $p$). Note that when analyzing the UKBb GWAS, reducing the reference panel SNPs does not change the regression SNPs, allowing us to directly compare model fits (i.e., we can directly compare log likelihoods in the two red columns with the corresponding values in the two blue columns).*

Having shown that the LDAK Model was more realistic than the GCTA Model, we then preferred estimates of functional enrichments from the LDAK+24Fun Model (an extension of the LDAK Model) to those from the Baseline Model (an extension of the GCTA Model). It was challenging to compare the Baseline and LDAK+24Fun Models directly. We were unable to reliably compute $logl_{\text{Old}}$ for the former, because SumHer originally used the weighted least-squares solver of LDSC[16] (see below) that often failed to converge (these issues are not apparent in LDSC, because when provided with a multi-parameter heritability model, it uses a one-step weighed least-squares solver that does not require convergence[17]). We showed that estimates from a hybrid model that combined the Baseline and LDAK+24Fun Models, were closer to those from the LDAK+24Fun Model than those from the Baseline Model (although again, convergence issues hindered solving this hybrid model). When we constructed Bayesian PRS assuming the Baseline and LDAK+24Fun Models, average prediction accuracy from the LDAK+24Fun Model was slightly higher, but the difference was not significant.

In this paper, we have shown that the Baseline and Baseline LD Models perform better than the LDAK+24Fun Model, both using $logl_{\text{SS}}$ and based on prediction of test statistics. Although counter-intuitive (that the LDAK Model outperforms the GCTA Model, but the Baseline and Baseline LD Models outperform the LDAK+24Fun Model), we believe that this finding provides insights into the genetic architecture of complex traits. While in general, the heritability distribution of causal variants depends on linkage disequilibrium (i.e., causal variants in regions of low LD tend to contribute more heritability than those in regions of high LD), when we condition on functional information, the relationship between heritability and LD is less pronounced. Note that our finding that the GCTA, Baseline and Baseline LD Models can all be improved by scaling based on MAF, indicates that whether or not we condition on functional information, there is always a tendency for heritability to depend on minor allele frequency.

When we proposed SumHer,[3] we estimated the parameters of the heritability model using the same iterative weighted least-squares solver that LDSC uses when provided with a one-parameter heritability model.[16] Correspondingly, we measured model fit via the approximate log likelihood

$$logl_{\text{Old}} = -\frac{U}{2}(\log(2\pi\gamma/U)+1) \quad \text{where} \quad U=\sum_j \frac{1}{u_j} \quad \text{and} \quad \gamma = \sum_j \frac{(S_j - \mathbb{E}[S_j])^2}{u_j},$$

obtained by assuming that the residuals from the least-squares regression are Gaussian distributed. Supplementary Figures 1, 2 & 7 show that $logl_{\text{SS}}$ performs better than $logl_{\text{Old}}$, reflecting that it is based on the more appropriate assumption that test statistics are Gamma distributed.

**Supplementary Note 2: Guidelines for using $logl_{SS}$ to evaluate and improve heritability models.** As a general rule, we recommend using the BLD-LDAK or BLD-LDAK+Alpha Models, as our analyses indicated that these are the best-performing across a wide range of traits. However, an alternative is to consider a variety of models, then use $logl_{SS}$ to decide which should be preferred or whether a hybrid model should be used instead. The figure below provides a demonstration of this approach. Note that for clarity, we consider only a selection of simple heritability models (we start with the GCTA, LDAK and LDAK-Thin Models); in practice, we would recommend more complex models (e.g., starting with the Baseline LD, BLD-LDAK and/or BLD-LDAK+Alpha Models).

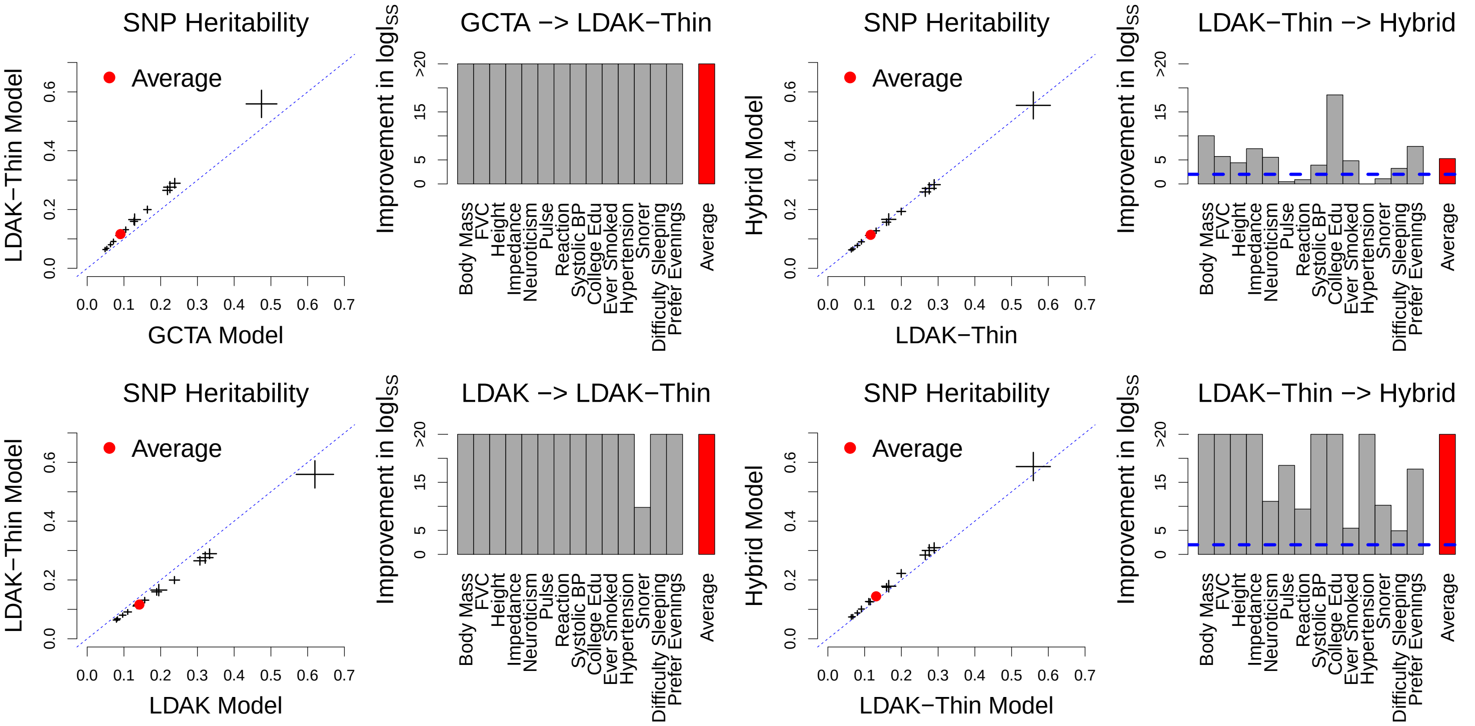


**Estimating SNP Heritability for the 14 UKBb GWAS.** Columns 1 & 3 compare estimates of SNP heritability from pairs of models (horizontal and vertical segments indicate 95% confidence intervals, while red points mark the average estimates). Columns 2 & 4 compare the difference in $logl_{SS}$ between pairs of models (note that the bars are truncated at 20). The top row considers the GCTA and LDAK-Thin Models. The first plot shows that the two models produce systematically different estimates of SNP heritability. The second plot shows that, if required to choose one set of results, we should use those from the LDAK-Thin Model, as this model consistently leads to higher model fit. Rather than choosing one model, we could instead use a hybrid model (i.e., combine the GCTA and LDAK-Thin Models). However, in this case, we find that a hybrid model is unnecessary; the third plot shows that its estimates are close to those from the LDAK-Thin Model, while the fourth plot shows that combining the two models produces only a small increase in $logl_{SS}$. The bottom row considers the LDAK and LDAK-Thin Models. Again, we see that the two models produce different estimates of SNP heritability, and that if required to choose, we should use those from the LDAK-Thin Model. However, this time we find that it is beneficial to use a hybrid model, as this substantially improves model fit. As a general rule, we would suggest increasing the complexity of a heritability model if the improvement in $logl_{SS}$ is at least two per parameter added.

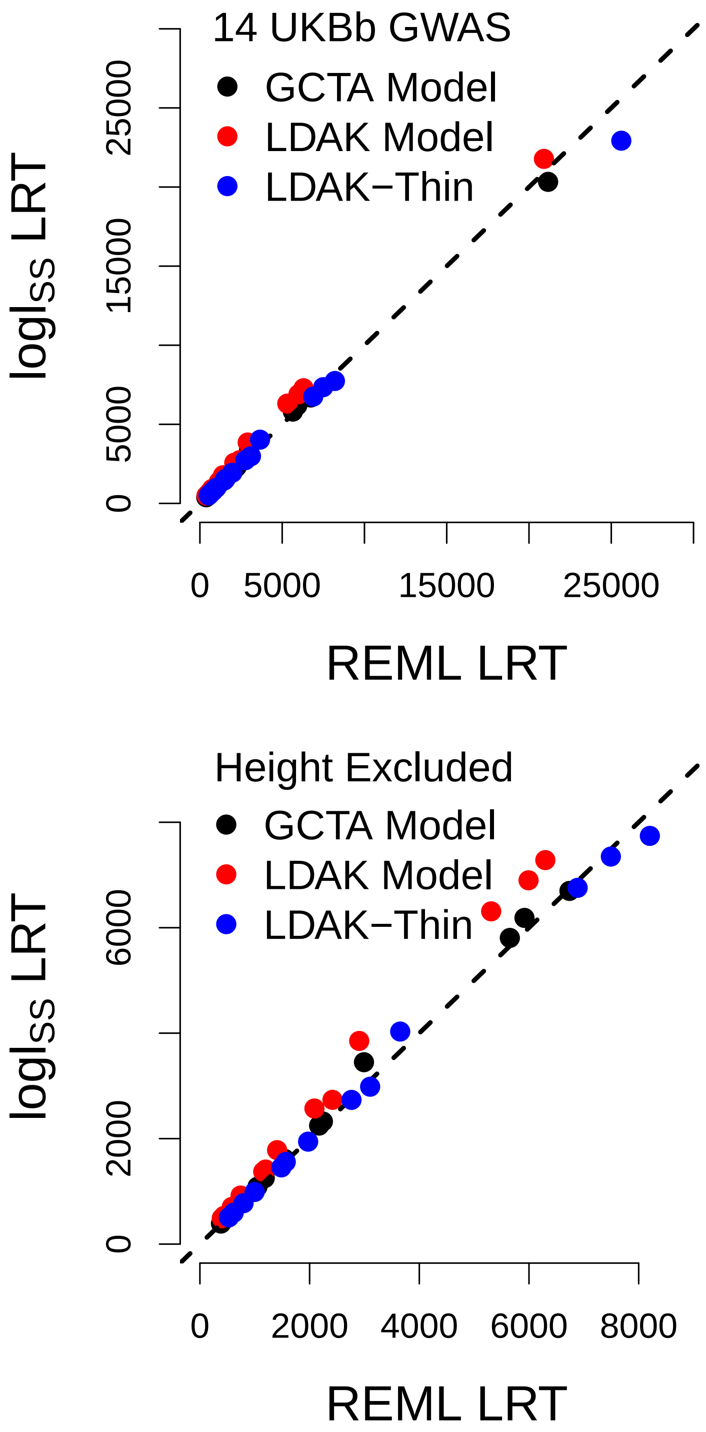


**Supplementary Figure 1: Comparison of likelihoods.** Plots compare likelihood ratio test (LRT) statistics (twice the improvement in log likelihood relative to the null model) computed using the likelihood from restricted maximum likelihood[18] (REML) with those from $logl_{\text{SS}}$, our new approximate likelihood, and $logl_{\text{Old}}$, the approximate likelihood we reported in the original version of SumHer[3] (see Supplementary Note 1 for details). We only analyze the 14 UKBb GWAS, because to perform REML requires individual-level data, and we only consider the GCTA, LDAK and LDAK-Thin Models, because REML is only feasible for simple heritability models. To ensure a fair comparison, when running SumHer we restrict the reference panel to the 4.7M GWAS SNPs. The bottom plots are zoomed versions of the top plots (obtained by excluding height, the most heritable trait). We see that the LRT statistics from $logl_{\text{SS}}$ are highly concordant with those from REML, indicating that the weights used when calculating $logl_{\text{SS}}$ perform well. We observe lower concordance between the LRT statistics from $logl_{\text{Old}}$ and those from REML, reflecting that $logl_{\text{Old}}$ was based on the assumption that test statistics were Gaussian distributed, rather than Gamma distributed.

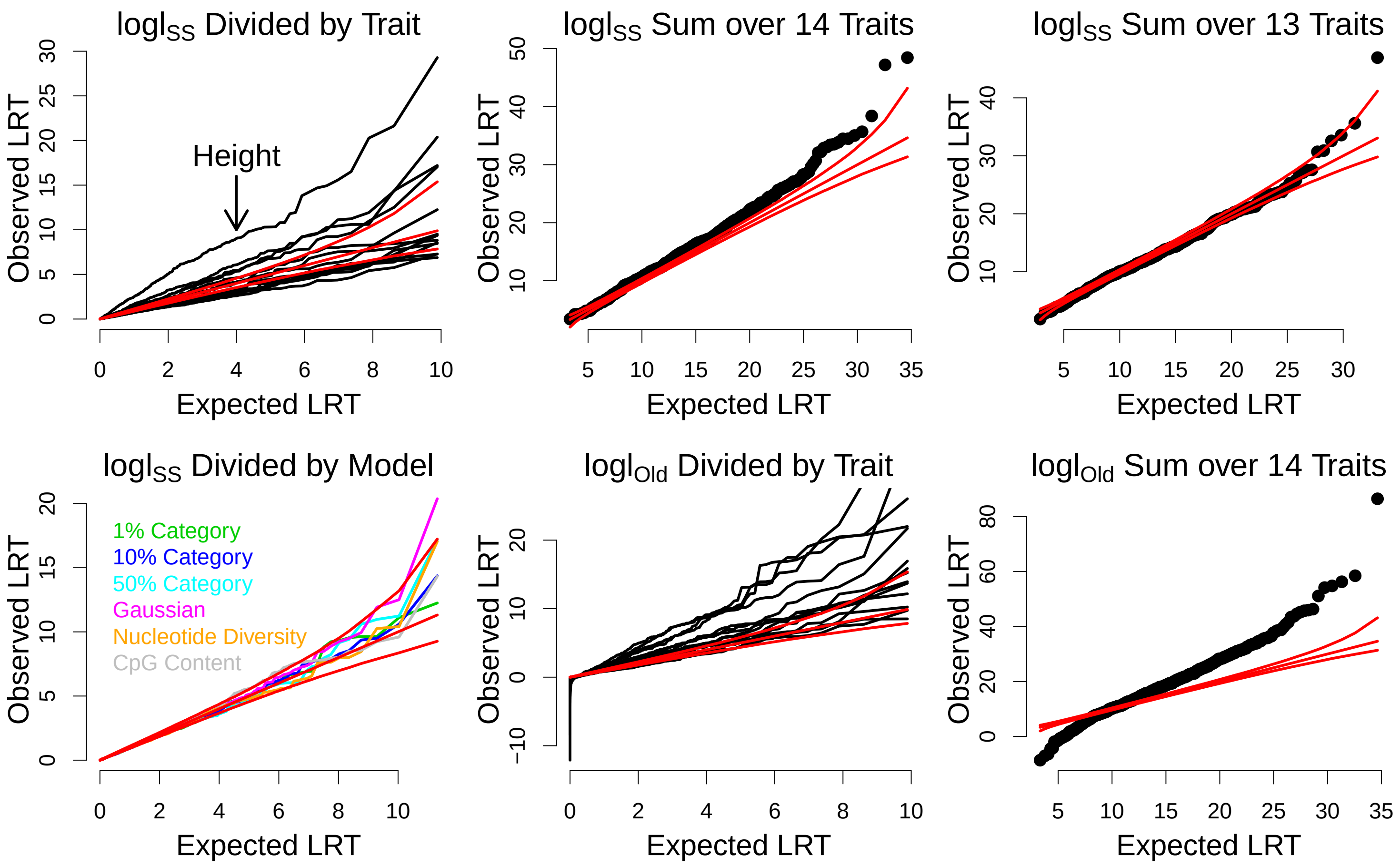


**Supplementary Figure 2: Likelihood ratio testing of heritability models.** We investigate the validity of performing a likelihood ratio test (LRT) using the approximate model likelihood $logl_{\text{SS}}$. We consider six heritability models, each of the form $\mathbb{E}[h_j^2] = \tau_1 + \tau_2 a_j$, where $a_j$ is one of three binary indicators, containing 1%, 10% or 50% of SNPs (picked at random), or one of three continuous annotations, either obtained by shuffling Annotations 4 or 6 of the Baseline LD Model (nuceleotide diversity or CpG content), or by sampling from a Gaussian distribution. For each heritability model, we construct 100 versions (i.e., we generate each $a_j$ 100 times), then test these on each of the 14 UKBb GWAS (in total, $6 \times 100 \times 14$ tests). Finally, for each test, we compute $T$, twice the improvement in log likelihood relative to the GCTA Model (obtained by setting $\tau_2 = 0$). Each of the six annotations is independent of genetic architecture, so if $T$ was computed using an exact likelihood, it would be asymptotically $\chi^2(1)$ distributed, and its sum across $D$ traits would be $\chi^2(D)$ distributed.

Each plot compares the observed values of either $T$ or its sum over traits, to their expected values were they computed from an exact likelihood (red lines mark $y = x$ and 95% confidence intervals for the expected values). For the first four plots we computed $T$ using $logl_{\text{SS}}$, our (new) approximate likelihood. In general, the observed distribution of $T$ is close to its expected distribution (first plot). However, we observe inflation for height, suggesting that $logl_{\text{SS}}$ performs worse for highly heritable traits (Supplementary Figure 3 shows that the estimated SNP heritability of height is double that for any other trait). Consequently, the sum of $T$ across 14 traits is close to its expectation (second plot), and even more so when we exclude height (third plot). The observed distribution of $T$ is similar regardless of which of the six heritability models is used (fourth plot). Note that in light of the inflation we observe for height, we confirm that the ranking of heritability models is the same when height is excluded (Supplementary Table 1). For the final two plots we compute $T$ using $logl_{\text{Old}}$, the approximate likelihood we reported in the original version of SumHer[3] (see Supplementary Note 1). Now the observed distributions of $T$ and its sum are very different to their expected distributions, reflecting that this likelihood was based on the assumption that test statistics were Gaussian distributed, rather than Gamma distributed.

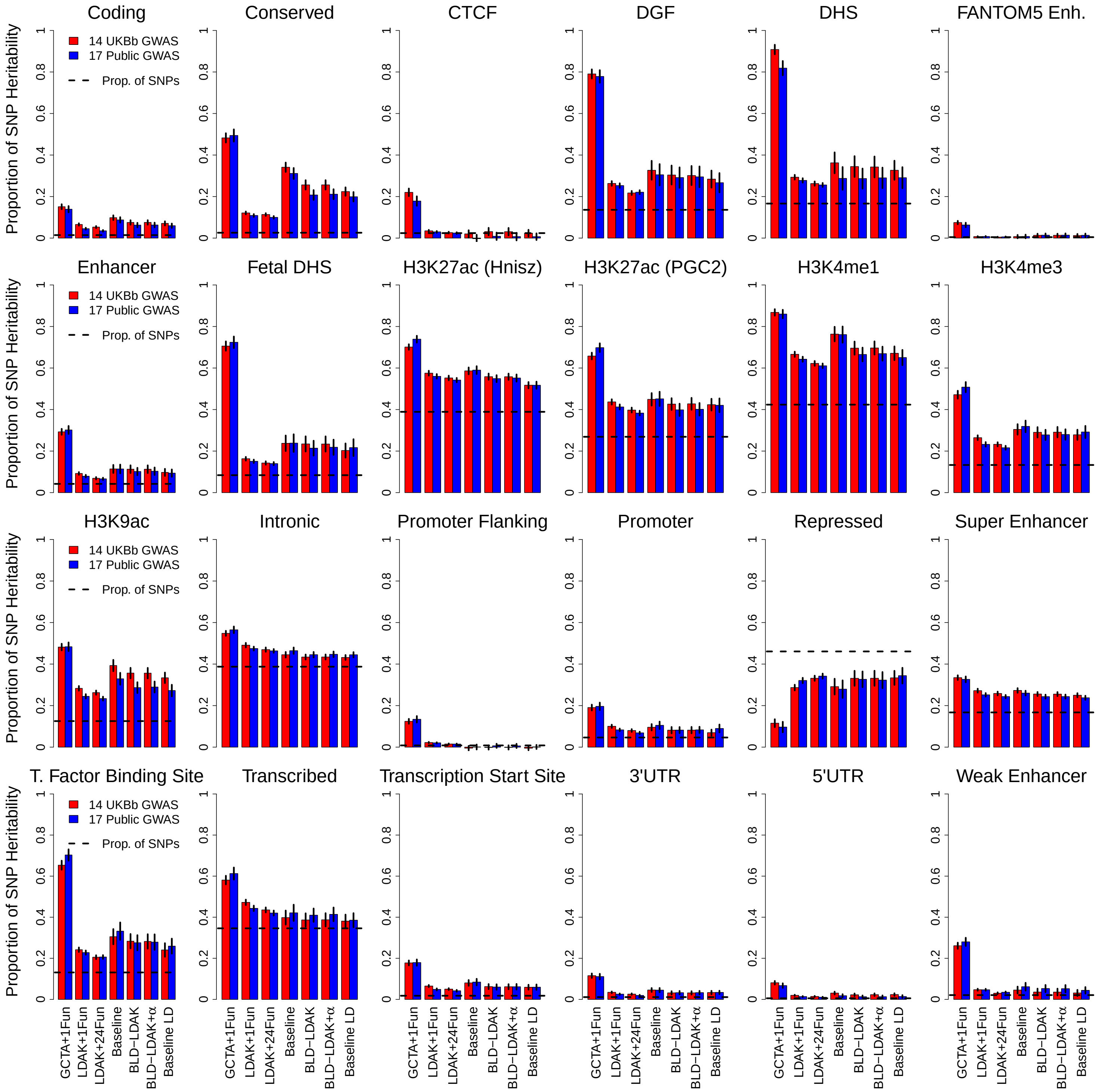


**Supplementary Figure 3: Estimated proportions of SNP heritability.** This is an expanded version of Figure 1d in the main text, and shows that estimates of functional enrichments tend to converge as the heritability model becomes more complex. Plots report the estimated proportion of SNP heritability contributed by each category of SNPs, averaged across either the 14 UKBb or 17 Public GWAS; vertical segments indicate 95% confidence intervals. Bars indicate the heritability model used and are ordered by number of parameters (see Supplementary Table 13 for definitions): GCTA+1Fun Model (two parameters, used by Gusev et al.[19]), LDAK+1Fun Model (two parameters, Speed et al.[10]), LDAK+24Fun Model (25 parameters, Speed et al.[3]), Baseline Model (53 parameters, Finucane et al.[17]), BLD-LDAK and BLD-LDAK+Alpha Models (66 and 67 parameters, this paper) and Baseline LD Model (75 parameters, Gazal et al.[16]). The estimated enrichment of a category is obtained by dividing its estimated proportions of SNP heritability by the proportion of SNPs it contains (horizontal dashed lines). Numerical values are provided in Supplementary Tables 5 & 6.

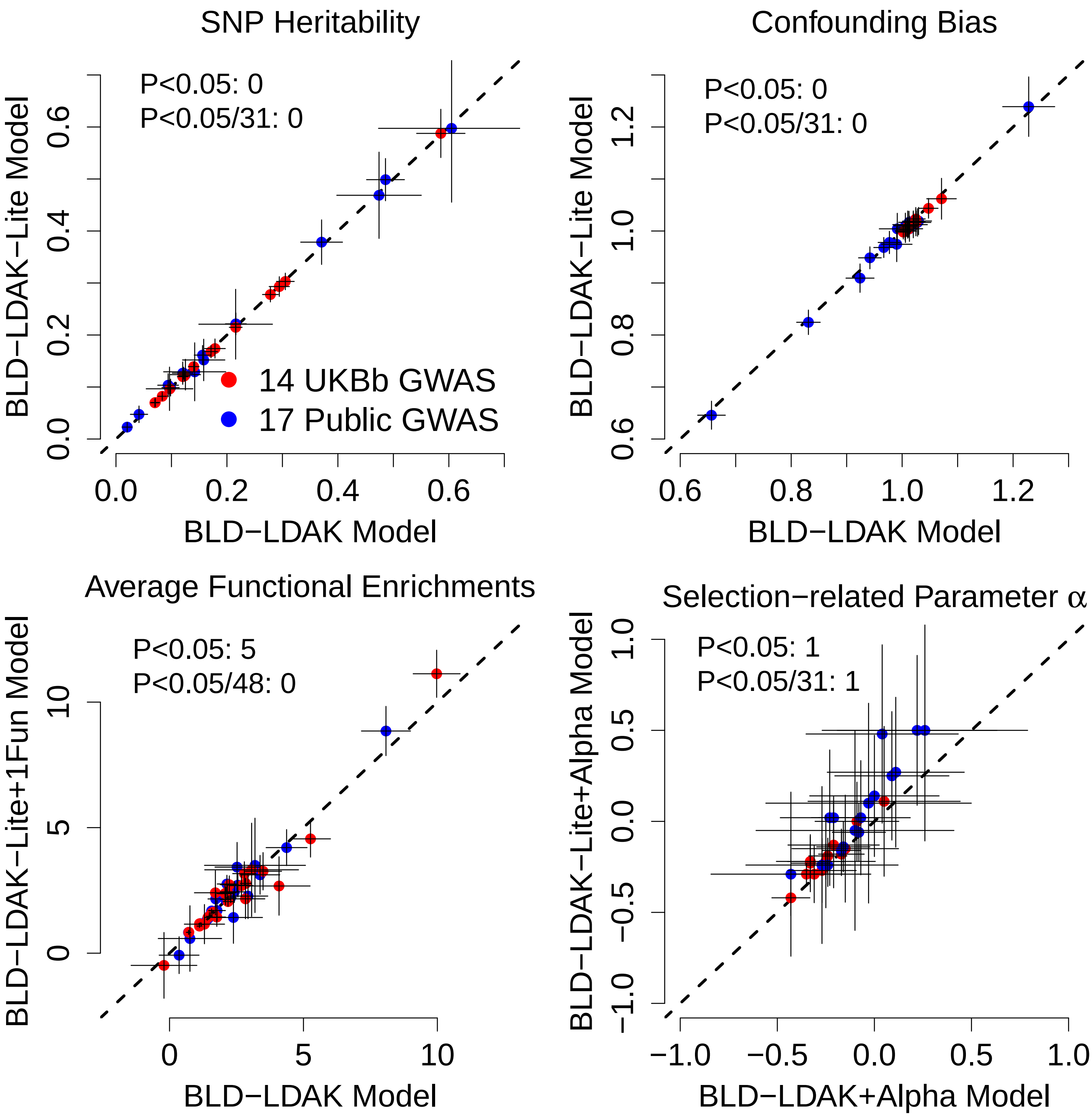


**Supplementary Figure 4: Reduced-complexity heritability models.** The seven-parameter BLD-LDAK-Lite is a reduced versions of the BLD-LDAK Model, obtained by removing two of the nine continuous annotations and all 57 binary annotations (Supplementary Table 8 explains how we used forward stepwise selection to decide which of the continuous annotations to retain). The nine-parameter BLD-LDAK-Lite+1Fun Model adds to the BLD-LDAK-Lite Model one function indicator and the corresponding 500 bp buffer, while the eight-parameter BLD-LDAK-Lite+Alpha Model is the same as the BLD-LDAK-Lite Model, except annotations are scaled by $[f_j(1-f_j]^{1+\alpha}$. These plots show that estimates of SNP heritability and confounding bias from the BLD-LDAK-Lite Model, and average estimates of functional enrichments from the BLD-LDAK-Lite+1Fun Model are close to the those from the BLD-LDAK Model, while estimates of $\alpha$ from the BLD-LDAK-Lite+Alpha Model are close to those from the BLD-LDAK+Alpha Model. Numbers indicate how many of the pairs of estimates are inconsistent either nominally or after Bonferroni correction. Numerical values are provided in Supplementary Tables 3, 4, 5, 6 & 7.

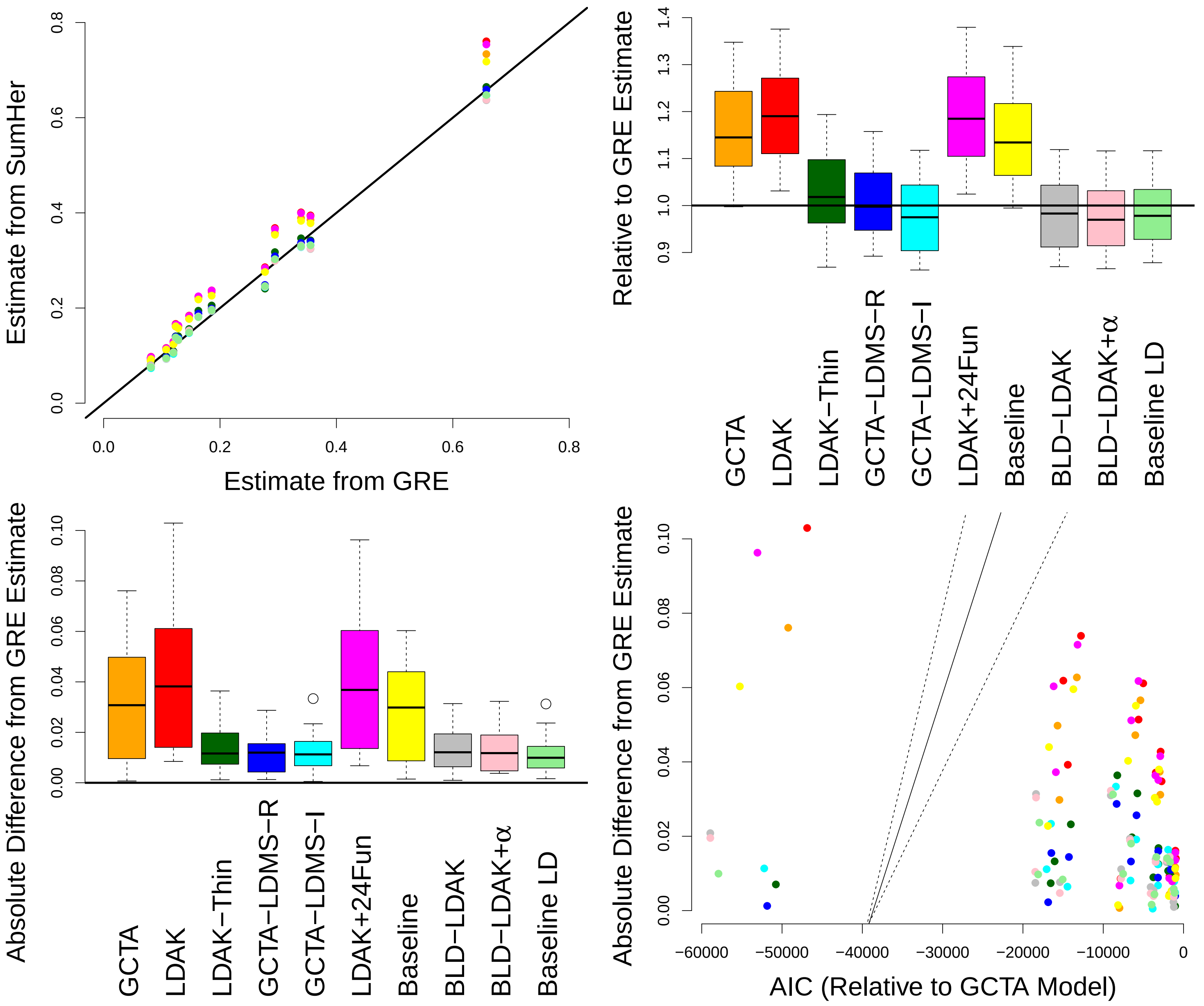


**Supplementary Figure 5: Comparison with GRE**. Hou *et al.*[20] proposed GRE, a method for estimating SNP heritability without specifying a heritability model. GRE requires individual level data and that there are more individuals than the number of SNPs on the largest chromosome. Here we compare estimates from GRE to those from SumHer for the 14 UKBb GWAS. To run GRE, we follow the instructions at www.github.com/bogdanlab/h2-GRE; to satisfy the sample size requirement, we use only the 623 k directly-genotyped SNPs (Hou *et al.* did likewise). For SumHer, we consider ten heritability models; to enable a fair comparison with GRE, we always restrict the reference panel to genotyped SNPs. The first three plots compare estimates of SNP heritability from GRE and SumHer. It is noticeable that when using only genotyped SNPs, changing the heritability model has a much smaller impact on estimates of SNP heritability than when using imputed SNPs (Supplementary Table 3); this reflects that with fewer SNPs, the impact of the prior assumptions is reduced. Nonetheless, if we consider GRE estimates to be the "gold standard", then this analysis indicates that the LDAK-Thin, GCTA-LDMS-R, GCTA-LDMS-I, BLD-LDAK, BLD-LDAK+Alpha and Baseline LD Models produce more accurate estimates of SNP heritability than the GCTA, LDAK, LDAK+24Fun and Baseline Models. In the fourth plot, the solid and dashed lines mark the point estimate and 95% confidence intervals for the gradient when regressing the absolute difference between estimates from SumHer and GRE on AIC (when performing this regression, we include an indicator for trait, to reflect that $logl_{SS}$ will tend to be higher for more heritable traits). If we again consider GRE estimates to be the gold standard, then the fact that the gradient is significantly positive ($P < 10^{-6}$) indicates that lower AIC implies more accurate estimates of SNP heritability.

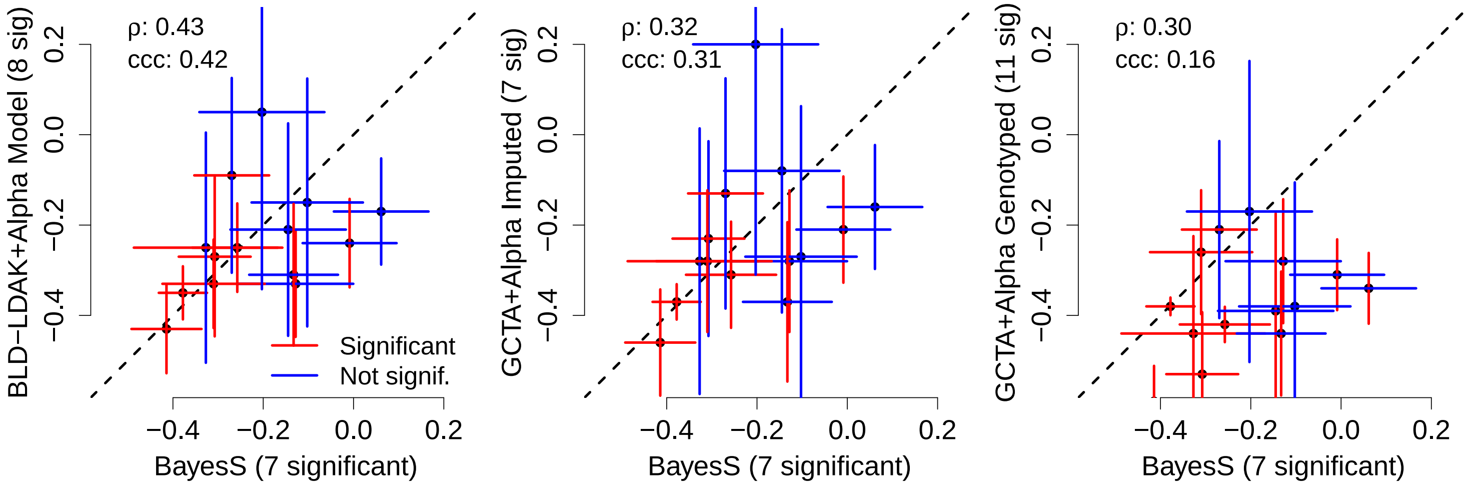


**Supplementary Figure 6: Comparison with BayesS.** Zeng *et al.*[21] proposed BayesS, a method for estimating the selection-related parameter $\alpha$ using individual-level data. It also provides estimates of $\pi$, the proportion of SNPs with non-zero effect, and $h^2_{\text{SNP}}$, the SNP heritability. BayesS assumes effect sizes have the distribution $\beta_j \sim \pi\mathbb{N}(0, [2f_j(1-f_j)]^\alpha\sigma^2_g) + (1-\pi)\delta_0$, where $f_j$ is the MAF of SNP $j$, $\sigma^2_g$ is a global variance term and $\delta_0$ is a point mass at zero. These three plots compare, for the 14 UKBb GWAS, estimates of $\alpha$ from BayesS to those from SumHer. Red lines indicate significant estimates (P<0.05/14); we summarize the similarity via the correlation ($\rho$) and concordance correlation coefficient[22] (ccc). When running BayesS, it is not computationally feasible to analyze all 130 k individuals and 4.7 M SNPs. Therefore, we restricted to the 623 k directly-genotyped SNPs (Zeng *et al.* did likewise), and analyzed only an eighth of the 130 k individuals at a time (we then report the inverse-variance weighted average of the eight estimates of $\alpha$). We first ran SumHer assuming the BLD-LDAK+Alpha Model (using the full reference panel), then twice assuming the "GCTA+Alpha Model", $\mathbb{E}[h^2_j] = [f_j(1-f_j)]^{1+\alpha}\tau_1$ (first using the full reference panel, then only the 623 k directly-genotyped SNPs). Under the BayesS model (assuming SNPs are in Hardy-Weinberg Equilibrium), $\mathbb{E}[h^2_j] = \pi[2f_j(1-f_j)]^{1+\alpha}\sigma^2_g$. Therefore, we had expected results from BayesS to be closest to those from SumHer assuming the GCTA+Alpha Model (and restricting the reference panel to genotyped SNPs). Instead, they are closest to those from SumHer assuming the BLD-LDAK+Alpha Model. This offers support for the claim of Zeng *et al.* that by using a mixture prior, their method is robust to the true relationship between $\mathbb{E}[h^2_j]$ and linkage disequilibrium (discussed in their Supplementary Note entitled "Robustness to LD heterogeneity").

Compared to BayesS, the main advantages of using SumHer to estimate $\alpha$ are computational. It requires only summary statistics, whereas BayesS needs individual-level data. It is able to consider dense SNP data, whereas BayesS is only feasible with sparse data. It uses at most 20 Gb of RAM, whereas our analyses using BayesS required 40 Gb (to have analyzed all 130 k individuals simultaneously would have required 300 Gb). Further, while both methods take approximately 300 computer hours, with SumHer most of this time is spent constructing the 31 tagging files (Supplementary Figure 8), which is required only once (for subsequent GWAS, it is only necessary to perform the 31 regressions, each of which takes about an hour). An additional advantage of SumHer is that it allows the user to specify complex heritability models, which is not currently possible in BayesS. On the other hand, BayesS provides estimates of the polygeneity of traits, which SumHer does not, and we note that BayesS estimates are slightly more precise (average standard deviation is 0.054, compared to 0.059 when running SumHer assuming the BLD-LDAK+Alpha Model).

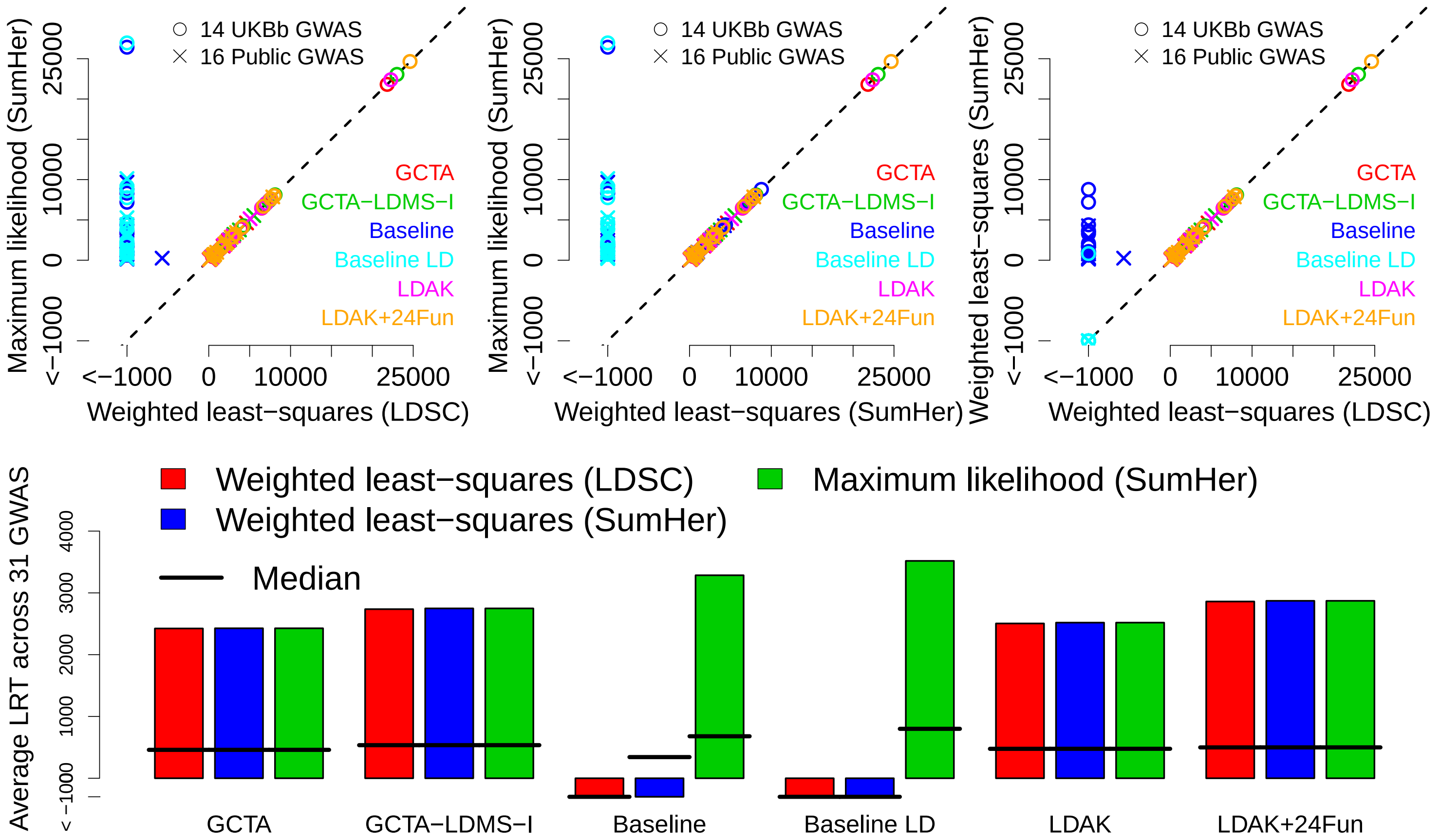


**Supplementary Figure 7: Comparison of weighted least-squares and maximum likelihood solvers.** The plots compare likelihood ratio test (LRT) statistics (twice the improvement in log likelihood relative to the null model), computed using $logl_{SS}$, our approximate model likelihood. We consider six heritability models (see Supplementary Table 13 for definitions), estimating parameters using either maximum likelihood (our recommended approach) or weighted least-squares regression (the approach used by LDSC[16] and previously by SumHer[3]). Note that when we estimate parameters for the Baseline and Baseline LD Models using weighted least-squares regression, we frequently obtain negative $E[S]$; so that we can compute $logl_{SS}$, we replace these with $10^{-6}$. These plots show that for the GCTA, GCTA-LDMS-I, LDAK and LDAK+24Fun Models (the simpler models), the two approaches give near-identical model fit. However, for the Baseline and Baseline LD Models (the more complex models), weighted least-squares regression often results in a worse fit, because it does not respect that test statistics are approximately Gamma distributed. Note that the reason we observe discordance between the weighted least-squares estimates from LDSC and SumHer (mainly evident for the Baseline Model), is because the SumHer weighted least-squares solver is always iterative,[3] whereas the LDSC solver is iterative when provided with a single-parameter heritability model,[16] but one-step when provided with a multi-parameter model.[17]

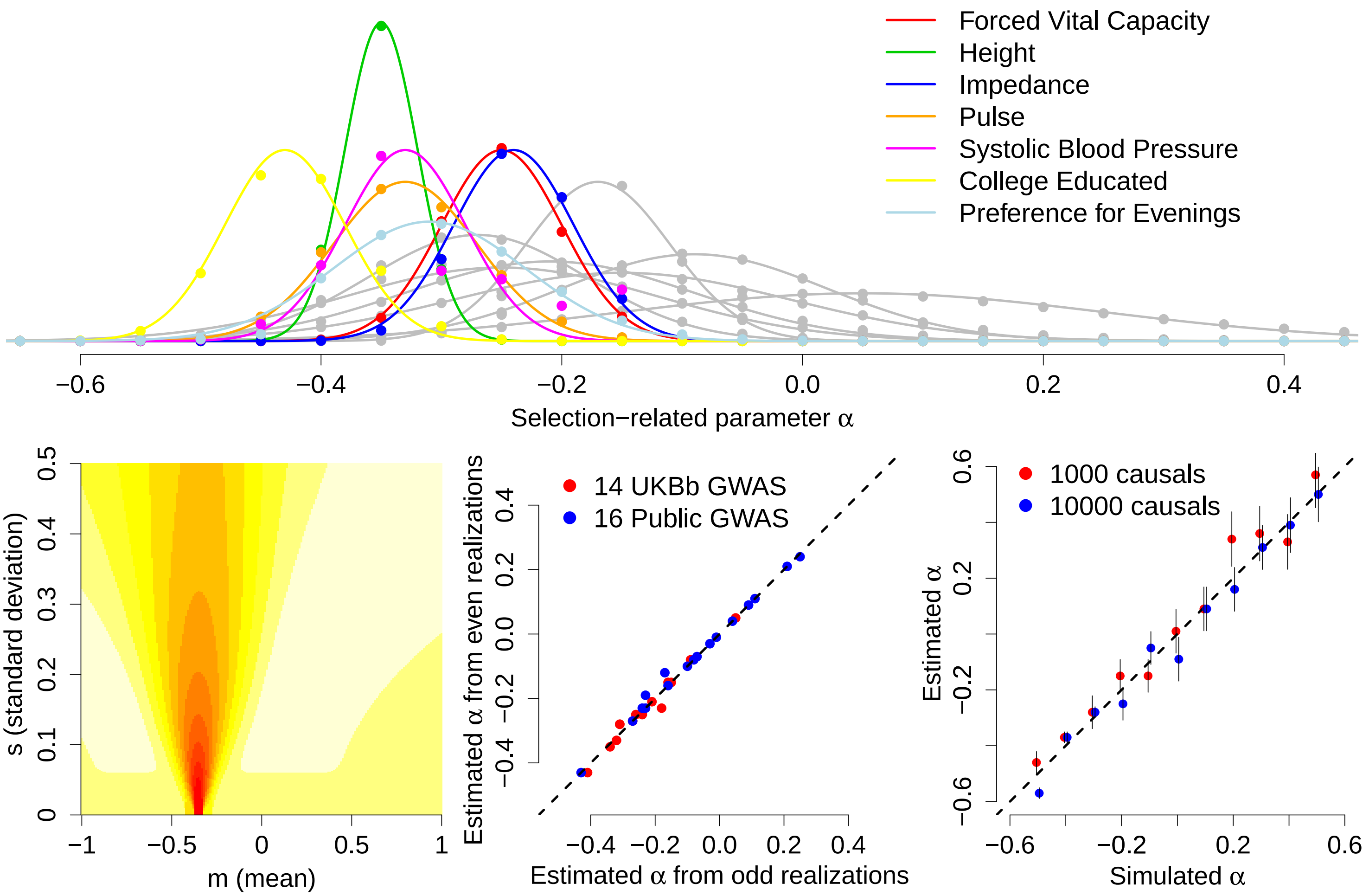


**Supplementary Figure 8: Estimating $\alpha$.** We are unable to estimate all 67 parameters of the BLD-LDAK+Alpha Model simultaneously. However, if we fix $\alpha$ (the non-linear parameter), we can solve for the $\tau_k$ (the linear parameters). Let $l_\alpha$ denote $logl_{\text{SS}}$ when the model is solved with $\alpha$ fixed. We first compute $l_{-1}, l_{-0.95}, \ldots, l_{0.45}, l_{0.5}$, then our estimate of $\alpha$ and its standard deviation are the pair $(m, s) \in \{-1, -0.99, \ldots, 0.99, 1.00\} \times \{0.01, 0.02, \ldots, 0.5\}$ that maximizes $R(m, s)$, the correlation between $\exp(l_\alpha)$ and $\exp(-\frac{(\alpha-m)^2}{2s^2})$. This corresponds to using a grid-search to identify the Gaussian distribution (with mean $m$ and standard deviation $s$) that most closely matches the 31 realizations of $l_\alpha$. The top plot compares the estimated likelihood curves to the realizations of $l_\alpha$ for each of the 14 UKBb GWAS; the seven labelled curves are those corresponding to significant estimates of $\alpha$ (P<0.05/31). The bottom left plot shows values of $R(m, s)$ for height (red indicates values closer to one); the horizontal and vertical lines mark the maximizing values of $m$ and $s$. The bottom middle plot shows that the estimates of $\alpha$ obtained using only the 16 odd-indexed realizations of $l_\alpha$ are similar to those obtained using only the 15 even-indexed realizations, indicating that our use of 31 values for $\alpha$ is sufficient. This grid-search strategy can be generalized. For example, when considering whether $\alpha$ varies across functional categories of SNPs (Supplementary Table 9), we first compute $l_{\alpha_1,\alpha_2}$ (where $\alpha_1$ and $\alpha_2$ denote, respectively, the values of $\alpha$ inside and outside the category being considered) for all $\{\alpha_1, \alpha_2\} \in \{-1, -.9, \ldots, 0.5\} \times \{-1, -.9, \ldots, 0.5\}$, then use a four-dimensional grid-search to identify the two (independent) Gaussian distributions that best fit the likelihood realizations.

For the bottom right plot, we perform simulations similar to those Zeng *et al.*[21] used to demonstrate their method BayesS (Supplementary Figure 6). Using the UKBb data (130 k individuals, 4.7 M SNPs), we generate phenotypes under the model $Y = \sum_j \beta_j X_j + e$, where $X_j$ are standardized genotypes for causal SNPs, $\beta_j \sim \mathbb{N}(0, [f_j(1-f_j)]^{1+\alpha}$, where $f_j$ is the MAF of causal SNP $j$, and $e \sim \mathbb{N}(0, \sigma_e^2)$. We consider 11 values of $\alpha$ $(-0.5, -0.4, \ldots, 0.4, 0.5)$; for each we first generate a phenotype with 1000 causal SNPs, then one with 10 000 causal SNPs (each time picking the causal SNPs at random and setting $\sigma_e^2$ so that the heritability is 0.5). We analyze each phenotype using the "GCTA+Alpha Model", $\mathbb{E}[h_j^2] = [f_j(1-f_j)]^{1+\alpha}\tau_1$, finding that for all 22 phenotypes the estimated value of $\alpha$ is close to its simulated value (as expected, considering that our assumed heritability model is consistent with the simulation model).

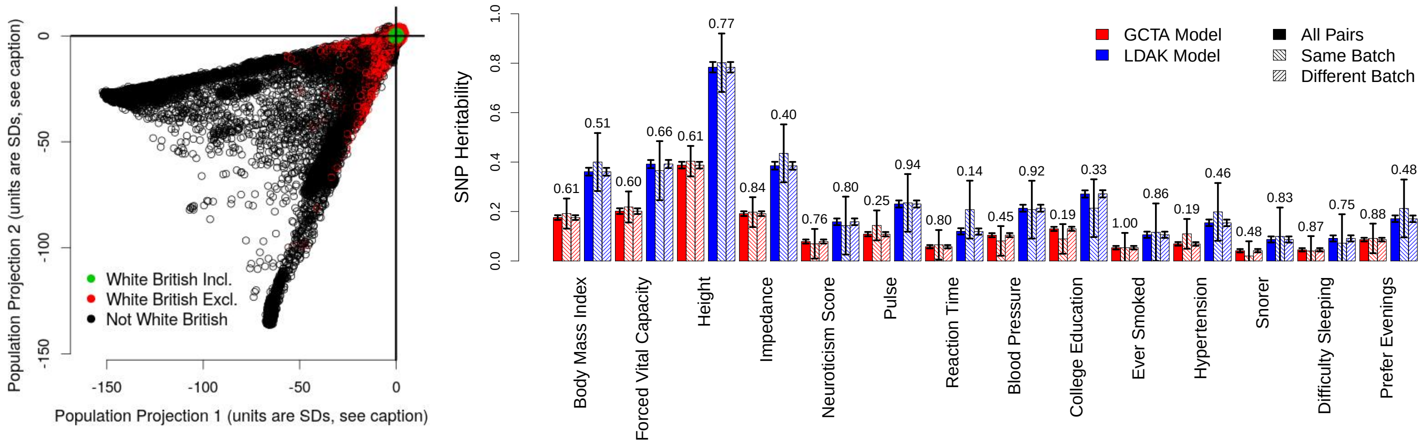


**Supplementary Figure 9: UKBb GWAS.** We first projected the 487 k UK Biobank[8,9] individuals onto the top two principal components from the 1000 Genome Project[23] data. The left plot shows the distance of each individual from the median projection, computed across the 430 k individuals recorded as being "White British" (datafield 21000, code 1001); the units are SDs (computed across the same 430 k individuals). We retained the 398 k individuals (marked in green) whose projections are consistent ($P$ >0.05) with this median (i.e., whose squared distance from the origin is less than 5.99, the 95% percentile of the $\chi^2(2)$ distribution). Next, we reduced to 225 k individuals (103 k males, 121 k females) by excluding those whose recorded sex and genetic sex did not match (datafields 31 & 21001), or who were missing values for either Townsend Deprivation Index or age (datafields 189 & 21022), or for any of the following 18 phenotypes: diastolic and systolic blood pressure (4079 & 4080), pulse rate (102), body mass index (21001), height (50), glasses (2207), basal metabolic rate (23105), impedance (23106), snorer (1210), preference for evenings (1180), difficulty falling asleep (1200), reaction time (20023), ever smoked (20160), forced vital capacity (3062), neuroticism score (20127), college education (6138), handedness (1707) and mouth problems (6149). We created two additional phenotypes by identifying which individuals were recorded as having asthma and hypertension (datafield 20002, codes 1111 & 1065). For all regressions, we used adjusted phenotypes, obtained by regressing the original phenotypes on 13 covariates: sex, age, Townsend Deprivation Index and 10 principal components (five from the data, five from the 1000 Genome Project projections).

To estimate the (total) heritability of each phenotype, we performed REML using a subset of 19 068 closely related individuals (each had estimated coefficient of relatedness $\geq$ 0.2 with at least one other individual). Based on this analysis, we decided to exclude four phenotypes, asthma, glasses, handedness and mouth problems, as each had estimated heritability below 0.1. We also excluded basal metabolic rate and diastolic blood pressure, as these were highly correlated with the more heritable phenotypes height and systolic blood pressure, respectively.

For the GWAS, we filtered so that no pair of individuals remained with allelic correlation >0.02 (the absolute value of the smallest correlation observed). This left us with 130 080 individuals (60 301 males, 69 779 females). Of the 93 M autosomal SNPs, we retained only the 4 725 151 with MAF$\geq$0.01, info score $\geq$0.99 (computed using LDAK[10]), that were present in the 1000 Genome Project reference panel provided on the LDSC website (www.github.com/bulik/ldsc), but not in the major histocompatibility complex (Chr 6:25-34 Mb). We converted genotype probabilities to hard calls using a 0.95 threshold, then performed linear association analysis for each of the 14 remaining phenotypes using PLINK 2.[24] To confirm our quality control was sufficient, we performed two tests (scripts at www.ldak.org/protocol). To estimate inflation due to population structure and relatedness, we computed $I = (H_1 + H_2 + H_3 + H_4 - H_{ALL})/3$, where $H_{All}$, $H_1$, $H_2$, $H_3$, $H_4$ are, respectively, the Haseman-Elston estimates of SNP heritabilities from the whole genome and each quarter; for all 14 Phenotypes, $0 \leq I \leq 0.001$, indicating minimal inflation. To estimate inflation due to genotyping errors, we compared $H_{\text{Same}}$ and $H_{\text{Diff}}$, Haseman-Elston estimates of the SNP heritability from only pairs of individuals in the same genotyping batch, and from only pairs of individuals in different genotyping batches (datafield 22000). The right plot reports for each phenotype the SNP heritability estimated from all individuals, $H_{\text{Same}}$ and $H_{\text{Diff}}$, first assuming the GCTA Model, then the LDAK Model (vertical segments mark 95% confidence intervals). The p-values above the bars indicate that the difference between $H_{\text{Same}}$ & $H_{\text{Diff}}$ is never significant (all $P \gg 0.05$).

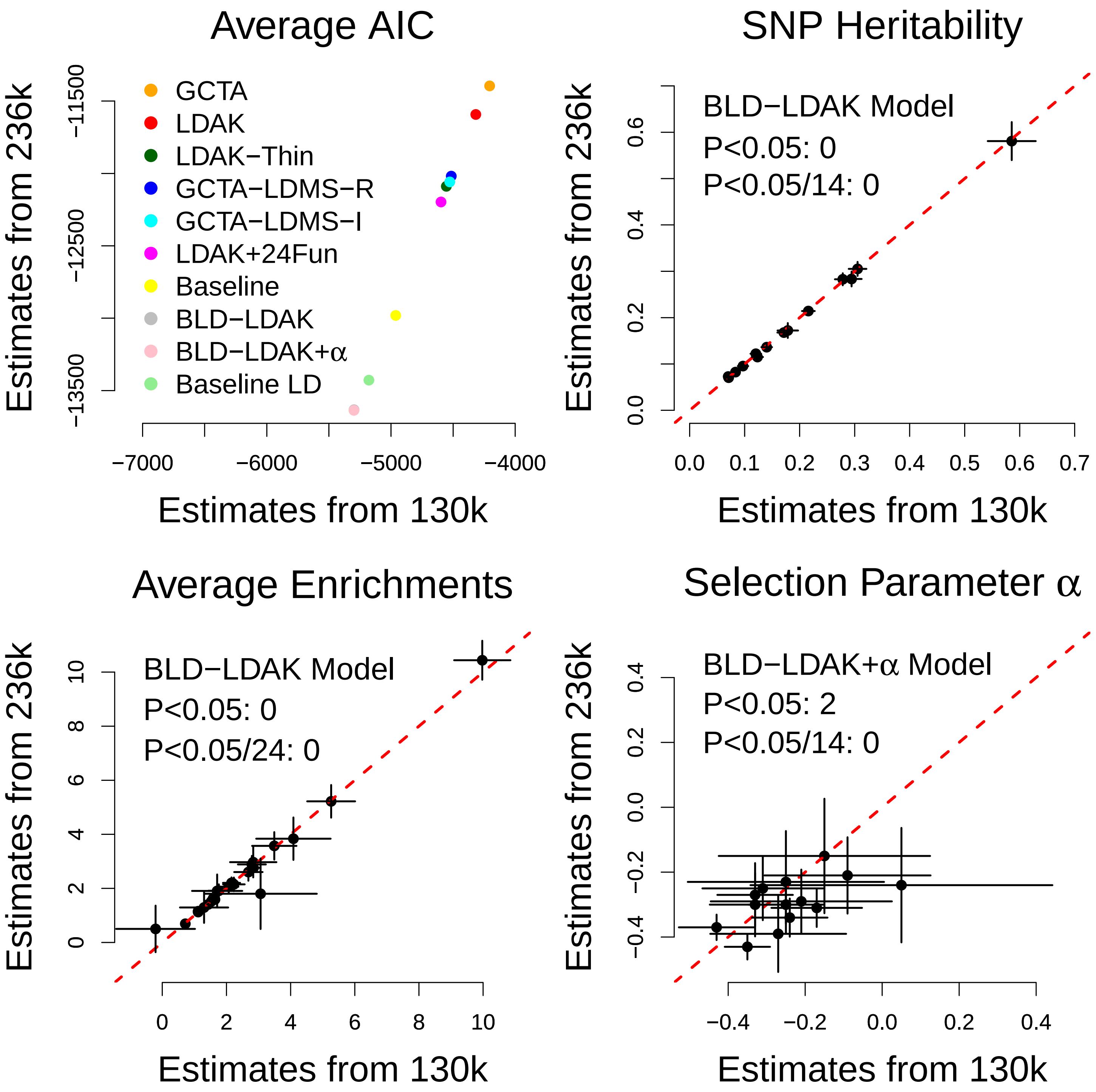


**Supplementary Figure 10: Reduced quality control for UKBb GWAS.** For our main analysis of the UKBb GWAS, we first identified individuals with values for all 14 phenotypes, then filtered so that no pair remained with allelic correlation >0.02 (Supplementary Table 9). As a secondary analysis, we instead identified individuals with values for any of the 14 phenotypes, then filtered so that no pair remained with allelic correlation >0.03125. This increased the number of individuals from 130 080 to 246 655, with on average 236 k phenotypic values per GWAS (range 201 k to 247 k). The first plot shows that increasing the sample size does not change the ranking of models based on the Akaike Information Criterion.[25] The remaining three plots shows that it does not significantly change estimates of SNP heritability or average functional enrichments from the BLD-LDAK Model, nor estimates of the selection-related parameter $\alpha$ from the BLD-LDAK+Alpha Model (horizontal and vertical segments indicate 95% confidence intervals; numbers indicate how many of the pairs of estimates are inconsistent either nominally or after Bonferroni correction).

| 1000GP Reference Panel | | All Reference SNPs (10.0 M) | | | | | | | | | Reduced Reference SNPs (4.7 M) | | | | |
|---|---|---|---|---|---|---|---|---|---|---|---|---|---|---|---|
| | | 14 UKBb GWAS | | | | | 17 Public GWAS | | | | 14 UKBb GWAS | | | | |
| Model | $K$ | $\overline{logl_{SS}}$ | $\overline{logl_{SS}}^{13}$ | $\overline{logl_{SS}}^{\dagger}$ | $\overline{\rho}$ | $\overline{\rho}'$ | $\overline{logl_{SS}}$ | $\overline{logl_{SS}}^{\dagger}$ | $\overline{\rho}$ | $\overline{\rho}'$ | $\overline{logl_{SS}}$ | $\overline{logl_{SS}}^{13}$ | $\overline{logl_{SS}}^{\dagger}$ | $\overline{\rho}$ | $\overline{\rho}'$ |
| GCTA | 1 | 2104 | 1427 | 851 | 0.059 | 0.073 | 480 | 173 | 0.038 | 0.043 | 1942 | 1300 | 767 | 0.053 | 0.067 |
| LDAK | 1 | 2160 | 1465 | 873 | 0.066 | 0.097 | 517 | 195 | 0.048 | 0.058 | 2160 | 1465 | 873 | 0.066 | 0.097 |
| LDAK-Thin | 1 | 2278 | 1557 | 931 | 0.073 | 0.104 | 617 | 245 | 0.056 | 0.070 | 2278 | 1557 | 931 | 0.073 | 0.104 |
| GCTA+1Fun | 2 | 2361 | 1569 | 972 | 0.068 | 0.091 | 695 | 261 | 0.056 | 0.077 | 2201 | 1443 | 889 | 0.061 | 0.086 |
| LDAK+1Fun | 2 | 2249 | 1505 | 909 | 0.067 | 0.106 | 607 | 228 | 0.055 | 0.070 | 2249 | 1505 | 909 | 0.067 | 0.106 |
| GCTA-LDMS-R | 20 | 2277 | 1570 | 928 | 0.071 | 0.091 | 640 | 257 | 0.052 | 0.055 | 2217 | 1521 | 895 | 0.071 | 0.094 |
| GCTA-LDMS-I | 20 | 2283 | 1572 | 940 | 0.069 | 0.090 | 625 | 255 | 0.048 | 0.052 | 2231 | 1529 | 909 | 0.068 | 0.090 |
| LDAK+24Fun | 25 | 2324 | 1554 | 959 | 0.068 | 0.117 | 704 | 283 | 0.057 | 0.077 | 2324 | 1554 | 959 | 0.068 | 0.117 |
| Baseline | 53 | 2534 | 1712 | 1079 | 0.074 | 0.093 | 909 | 377 | 0.061 | 0.088 | 2361 | 1576 | 989 | 0.066 | 0.086 |
| BLD-LDAK | 66 | **2715** | **1870** | **1167** | **0.085** | **0.121** | **1098** | **483** | **0.071** | **0.100** | **2657** | **1825** | **1134** | **0.082** | **0.122** |
| BLD-LDAK+ALPHA | 67 | **2716** | **1871** | **1169** | **0.086** | **0.120** | **1100** | **483** | **0.071** | **0.097** | **2661** | **1829** | **1134** | **0.082** | **0.123** |
| Baseline LD | 75 | 2665 | 1831 | 1155 | 0.082 | 0.120 | 1043 | 457 | 0.070 | 0.097 | 2568 | 1758 | 1106 | 0.078 | 0.120 |

| UKBb Reference Panel | | All Reference SNPs (10.0 M) | | | | | | | | | Reduced Reference SNPs (4.7 M) | | | | |
|---|---|---|---|---|---|---|---|---|---|---|---|---|---|---|---|
| | | 14 UKBb GWAS | | | | | 17 Public GWAS | | | | 14 UKBb GWAS | | | | |
| GCTA | 1 | 2019 | 1373 | 843 | 0.060 | 0.074 | 475 | 173 | 0.039 | 0.045 | 1868 | 1253 | 762 | 0.053 | 0.067 |
| LDAK | 1 | 2064 | 1404 | 861 | 0.068 | 0.100 | 505 | 191 | 0.048 | 0.059 | 2064 | 1404 | 861 | 0.068 | 0.100 |
| LDAK-Thin | 1 | 2191 | 1502 | 931 | 0.075 | 0.105 | 608 | 243 | 0.056 | 0.069 | 2191 | 1502 | 931 | 0.075 | 0.105 |
| GCTA+1Fun | 2 | 2263 | 1509 | 967 | 0.069 | 0.092 | 685 | 266 | 0.056 | 0.078 | 2114 | 1390 | 889 | 0.062 | 0.087 |
| LDAK+1Fun | 2 | 2153 | 1444 | 897 | 0.070 | 0.111 | 597 | 228 | 0.055 | 0.072 | 2153 | 1444 | 897 | 0.070 | 0.111 |
| GCTA-LDMS-R | 20 | 2198 | 1517 | 930 | 0.073 | 0.093 | 635 | 264 | 0.051 | 0.053 | 2155 | 1480 | 906 | 0.072 | 0.095 |
| GCTA-LDMS-I | 20 | 2190 | 1510 | 935 | 0.071 | 0.091 | 614 | 258 | 0.048 | 0.051 | 2152 | 1476 | 912 | 0.069 | 0.091 |
| LDAK+24Fun | 25 | 2224 | 1489 | 947 | 0.071 | 0.123 | 698 | 289 | 0.058 | 0.078 | 2224 | 1489 | 947 | 0.071 | 0.123 |
| Baseline | 53 | 2426 | 1645 | 1067 | 0.075 | 0.093 | 895 | 380 | 0.061 | 0.089 | 2265 | 1516 | 982 | 0.067 | 0.087 |
| BLD-LDAK | 66 | **2607** | **1800** | **1151** | **0.088** | **0.122** | **1081** | **492** | **0.073** | **0.103** | **2550** | **1755** | **1120** | **0.085** | **0.122** |
| BLD-LDAK+ALPHA | 67 | **2608** | **1801** | **1151** | **0.088** | **0.123** | **1083** | **492** | **0.072** | **0.097** | **2556** | **1761** | **1121** | **0.085** | **0.124** |
| Baseline LD | 75 | 2547 | 1755 | 1135 | 0.085 | 0.120 | 1026 | 462 | 0.071 | 0.098 | 2449 | 1681 | 1087 | 0.080 | 0.121 |

**Supplementary Table 1: Average $logl_{SS}$ for main heritability models.** The 12 models are defined in Supplementary Table 13. $K$ is the number of parameters. $logl_{SS}$ is the approximate log likelihood, while $logl^{\dagger}_{SS}$ is an unweighted version, obtained by first thinning the regression SNPs (those used when regressing $S_j$ on $\mathbb{E}[S_j]$) so that no pair within 1 cM has $r^2_{jl} > 0.1$, then setting the $1/u_j$ (the SNP weights) to one. We report $\overline{logl_{SS}}$ and $\overline{logl_{SS}}^{\dagger}$, the increase in $logl_{SS}$ and $logl^{\dagger}_{SS}$ relative to the null model, averaged across either the 14 UKBb or 17 Public GWAS. We also report $\overline{logl_{SS}}^{13}$, the increase in $logl_{SS}$ averaged across only 13 of the UKBb GWAS (we exclude height, as our analysis in Supplementary Figure 2 indicates that $logl_{SS}$ is less accurate for this highly-heritable trait). Whether we order based on $\overline{logl_{SS}}$, $\overline{logl_{SS}}^{13}$ or $\overline{logl_{SS}}^{\dagger}$, the BLD-LDAK and BLD-LDAK+Alpha Models (marked in **bold**) rank best. This is also the case if we order models based on the Akaike Information Criterion[25] (equivalent to ranking them based on $logl_{SS}$-$K$).

We also compare models based on prediction accuracy. For each SNP we compute $\mathbb{E}[S_j]$, its expected test statistic given the heritability model (e.g., for the GCTA Model, $\mathbb{E}[S_j] = 1 + n\sum_{l \text{ near } j} r^2_{jl}\tau_1$). This requires estimates of the heritability parameters; to avoid overfitting, we use leave-one-chromosome-out cross-validation (e.g., when computing $\mathbb{E}[S_j]$ for Chromosome 1 SNPs, we use parameter estimates obtained from Chromosomes 2-22). We calculate the weighted correlation $\rho = C(S_j, \mathbb{E}[S_j])/\sqrt{C(S_j, S_j)C(\mathbb{E}[S_j], \mathbb{E}[S_j])}$, where $C(a_j, b_j) = (\sum_j a_j b_j/u_j)(\sum_j 1/u_j) - (\sum_j a_j/u_j)(\sum_j b_j/u_j)$, then report $\overline{\rho}$, the average of $\rho$ across either the 14 UKBb or 17 Public GWAS. For completeness, we also report $\overline{\rho}'$, the average of the unweighted correlation (obtained by setting $u_j = 1$). The s.d. of $\overline{\rho}$ and $\overline{\rho}'$ is always 0.001. We see that the ranking of models based on prediction accuracy is consistent with the ranking based on model fit.

For the top table, we use the 1000 Genome Project[23] reference panel (489 European individuals, recorded for 10.0 M SNPs). The bottom table shows that the ranking of models is the same if we instead use a UK Biobank reference panel (constructed by randomly picking 2 000 of the 130 k individuals used in the UKBb GWAS, then extracting their genotypes for the 10.0 M SNPs present in the 1000 Genomes Project panel). For the UKBb GWAS, we repeat the analysis restricting the reference panel to the 4.7 M GWAS SNPs (the regression SNPs are unchanged, so values from this analysis can be directly compared with those obtained using the full reference panel). We find that model fit and prediction accuracy are generally higher when using the full reference panel, reflecting that when predicting how much heritability a SNP tags, it is beneficial to consider contributions from all neighboring SNPs, rather than just those present in the GWAS (there is no difference when assuming either the LDAK or LDAK+24Fun Models, because only the 4.7 M SNPs are used when computing the LDAK weightings).

| Heritability Model | GCTA | LDAK | LDAK-Thin | GCTA+1Fun | LDAK+1Fun | GCTA-LDMS-R | GCTA-LDMS-I | LDAK+24Fun | Baseline | BLD-LDAK | BLD-LDAK+Alpha | Baseline LD |
|---|---|---|---|---|---|---|---|---|---|---|---|---|
| Body Mass Index | 3234 (12) | 3252 (11) | 3469 (7) | 3340 (8) | 3262 (10) | 3514 (5) | 3511 (6) | 3345 (9) | 3598 (4) | 3944 (2) | 3946 (1) | 3875 (3) |
| Forced Vital Capacity | 3379 (12) | 3525 (11) | 3686 (7) | 3905 (5) | 3683 (8) | 3656 (10) | 3676 (9) | 3807 (6) | 4159 (4) | 4422 (1) | 4422 (2) | 4310 (3) |
| Height | 10908 (12) | 11191 (11) | 11653 (8) | 12658 (5) | 11916 (7) | 11469 (10) | 11531 (9) | 12324 (6) | 13214 (4) | 13704 (2) | 13709 (1) | 13506 (3) |
| Impedance | 3732 (12) | 3784 (11) | 4018 (8) | 4139 (5) | 3910 (10) | 4060 (6) | 4057 (7) | 4009 (9) | 4395 (4) | 4701 (1) | 4701 (2) | 4593 (3) |
| Neuroticism Score | 732 (9) | 726 (10) | 798 (6) | 746 (8) | 726 (11) | 813 (7) | 818 (5) | 747 (12) | 856 (4) | 968 (1) | 968 (2) | 959 (3) |
| Pulse | 1309 (12) | 1412 (11) | 1540 (7) | 1578 (6) | 1528 (8) | 1475 (10) | 1484 (9) | 1612 (5) | 1830 (4) | 1979 (1) | 1980 (2) | 1930 (3) |
| Reaction Time | 460 (12) | 477 (8) | 519 (4) | 475 (11) | 477 (9) | 535 (6) | 536 (5) | 500 (10) | 541 (7) | 629 (1) | 629 (2) | 621 (3) |
| Systolic Blood Pressure | 1289 (12) | 1330 (11) | 1418 (6) | 1527 (5) | 1383 (10) | 1434 (8) | 1434 (9) | 1441 (7) | 1641 (4) | 1780 (2) | 1781 (1) | 1748 (3) |
| College Education | 1964 (12) | 2002 (10) | 2076 (7) | 2004 (9) | 2003 (11) | 2119 (6) | 2133 (5) | 2045 (8) | 2198 (4) | 2391 (2) | 2396 (1) | 2365 (3) |
| Ever Smoked | 382 (9) | 370 (10) | 413 (5) | 390 (8) | 370 (11) | 422 (6) | 418 (7) | 388 (12) | 466 (4) | 544 (1) | 544 (2) | 525 (3) |
| Hypertension | 627 (12) | 708 (11) | 737 (9) | 803 (5) | 764 (7) | 757 (8) | 742 (10) | 802 (6) | 871 (4) | 981 (2) | 983 (1) | 947 (3) |
| Snorer | 232 (12) | 255 (9) | 265 (4) | 255 (10) | 259 (7) | 277 (8) | 278 (6) | 275 (11) | 316 (5) | 372 (1) | 372 (2) | 363 (3) |
| Difficulty Falling Asleep | 280 (9) | 275 (10) | 304 (6) | 289 (8) | 275 (11) | 331 (4) | 327 (5) | 283 (12) | 343 (7) | 407 (2) | 408 (1) | 408 (3) |
| Preference for Evenings | 926 (12) | 929 (9) | 1000 (6) | 949 (8) | 929 (10) | 1023 (4) | 1022 (5) | 951 (11) | 1048 (7) | 1187 (1) | 1187 (2) | 1160 (3) |
| **Average** | 2104 | 2160 | 2278 | 2361 | 2249 | 2277 | 2283 | 2324 | 2534 | 2715 | 2716 | 2665 |

| Heritability Model | GCTA | LDAK | LDAK-Thin | GCTA+1Fun | LDAK+1Fun | GCTA-LDMS-R | GCTA-LDMS-I | LDAK+24Fun | Baseline | BLD-LDAK | BLD-LDAK+Alpha | Baseline LD |
|---|---|---|---|---|---|---|---|---|---|---|---|---|
| Coronary Artery | 48 (11) | 60 (9) | 82 (8) | 117 (1) | 105 (4) | 74 (10) | 65 (12) | 115 (6) | 151 (5) | 171 (2) | 171 (3) | 165 (7) |
| Crohn's Disease | 105 (12) | 157 (11) | 174 (10) | 337 (6) | 260 (7) | 203 (9) | 207 (8) | 367 (5) | 587 (4) | 734 (2) | 737 (1) | 688 (3) |
| Ever Smoked? | 32 (10) | 47 (1) | 45 (3) | 34 (9) | 48 (2) | 62 (4) | 59 (6) | 58 (8) | 67 (12) | 106 (5) | 106 (7) | 99 (11) |
| Inflammatory Bowel | 119 (12) | 205 (11) | 206 (10) | 411 (6) | 358 (7) | 242 (8) | 239 (9) | 491 (5) | 680 (4) | 857 (2) | 860 (1) | 793 (3) |
| Rheumatoid Arthritis | 77 (12) | 116 (10) | 124 (9) | 242 (6) | 190 (7) | 167 (8) | 124 (11) | 308 (5) | 406 (4) | 541 (1) | 541 (2) | 521 (3) |
| Schizophrenia | 878 (12) | 913 (11) | 1117 (5) | 993 (8) | 921 (10) | 1134 (6) | 1112 (7) | 1010 (9) | 1244 (4) | 1582 (1) | 1583 (2) | 1500 (3) |
| Type 2 Diabetes | 138 (12) | 186 (11) | 255 (7) | 289 (5) | 237 (8) | 239 (9) | 225 (10) | 294 (6) | 382 (4) | 475 (2) | 476 (1) | 439 (3) |
| Bone Mineral Density | 38 (12) | 53 (11) | 65 (7) | 93 (5) | 66 (8) | 81 (9) | 72 (10) | 96 (6) | 169 (4) | 211 (2) | 213 (1) | 205 (3) |
| Body Mass Index | 1226 (12) | 1253 (11) | 1535 (7) | 1399 (9) | 1283 (10) | 1660 (5) | 1598 (6) | 1425 (8) | 1848 (4) | 2291 (1) | 2291 (2) | 2165 (3) |
| Depressive Symptoms | 65 (8) | 59 (11) | 85 (4) | 65 (9) | 62 (10) | 95 (6) | 94 (7) | 80 (12) | 129 (5) | 170 (1) | 171 (2) | 167 (3) |
| Height | 2312 (12) | 2570 (11) | 2963 (8) | 4201 (5) | 3476 (7) | 2800 (9) | 2772 (10) | 3925 (6) | 4842 (4) | 5275 (1) | 5275 (2) | 5031 (3) |
| Menarche Age | 1514 (11) | 1511 (12) | 1814 (7) | 1625 (9) | 1521 (10) | 1892 (5) | 1878 (6) | 1718 (8) | 2139 (4) | 2737 (1) | 2738 (2) | 2612 (3) |
| Menopause Age | 88 (12) | 112 (11) | 149 (8) | 239 (6) | 184 (7) | 162 (9) | 155 (10) | 294 (5) | 430 (4) | 502 (2) | 505 (1) | 474 (3) |
| Neuroticism | 261 (9) | 230 (11) | 305 (7) | 270 (8) | 230 (12) | 339 (4) | 334 (5) | 255 (10) | 360 (6) | 473 (2) | 474 (1) | 460 (3) |
| Subjective Well-Being | 77 (5) | 56 (10) | 81 (2) | 81 (4) | 56 (11) | 91 (8) | 91 (9) | 68 (12) | 128 (6) | 147 (1) | 147 (3) | 149 (7) |
| Waist-Hip Ratio | 202 (12) | 260 (11) | 313 (10) | 365 (5) | 316 (8) | 343 (7) | 332 (9) | 351 (6) | 503 (4) | 628 (2) | 629 (1) | 584 (3) |
| Years Education | 985 (12) | 1011 (10) | 1167 (7) | 1050 (9) | 1011 (11) | 1293 (5) | 1270 (6) | 1116 (8) | 1392 (4) | 1772 (2) | 1775 (1) | 1678 (3) |
| **Average** | 480 | 517 | 617 | 695 | 607 | 640 | 625 | 704 | 909 | 1098 | 1100 | 1043 |

**Supplementary Table 2: $logl_{SS}$ for individual GWAS.** We report the increase in $logl_{SS}$ relative to the null model. Numbers in brackets indicate the ranking of each heritability model based on the Akaike Information Criterion[25] (equal to $2K - 2logl_{SS}$, where $K$ is the number of parameters in the model); for each GWAS, we highlight in red, blue and green, respectively, the three heritability models ranked first, second and third. The 12 models are defined in Supplementary Table 13. When we restrict to the nine existing heritability models, the Baseline LD Model ranks first for 28 out of 31 GWAS. However, when we include the three novel heritability models, the BLD-LDAK and BLD-LDAK+Alpha Models are now the top-two ranking models for 28 out of 31 GWAS.

Although we have generally compared models across multiple GWAS, we note that there is often sufficient power to perform model comparisons for individual GWAS. For example, if we set the significance threshold to 0.01, then we can justify adding a parameter to a heritability model if its inclusion increases $logl_{SS}$ by at least 3.3 (because 2×3.3 is the 99th percentile of the $\chi^2(1)$ distribution). A potential application would be to perform stepwise selection to see which of the 24 functional annotations are most important for each trait; as well as providing insights into genetic architecture, excluding the non-significant annotations could potentially lead to improved prediction accuracy. Alternatively, we could use trait-specific versions of the BLD-LDAK Model, obtained by fixing $\alpha$ to values other than -0.25; by comparing $logl_{SS}$ for the BLD-LDAK and BLD-LDAK+Alpha Model (the latter estimates $\alpha$ from the data), we can see that this would be most beneficial for height and college education.

| Heritability Model | GCTA | LDAK | LDAK -Thin | GCTA -LDMS-R | GCTA -LDMS-I | LDAK +24Fun | Baseline | BLD-LDAK | BLD-LDAK +Alpha | Baseline LD | BLD-LDAK -LITE |
|---|---|---|---|---|---|---|---|---|---|---|---|
| Body Mass Index | 0.22 (0.01) | 0.31 (0.01) | 0.27 (0.01) | 0.26 (0.01) | 0.26 (0.01) | 0.30 (0.01) | 0.22 (0.01) | 0.28 (0.01) | 0.28 (0.01) | 0.24 (0.01) | 0.28 (0.01) |
| Forced Vital Capacity | 0.23 (0.01) | 0.32 (0.01) | 0.28 (0.01) | 0.27 (0.01) | 0.27 (0.01) | 0.32 (0.01) | 0.23 (0.01) | 0.29 (0.01) | 0.29 (0.01) | 0.25 (0.01) | 0.29 (0.01) |
| Height | 0.47 (0.02) | 0.62 (0.03) | 0.56 (0.02) | 0.54 (0.02) | 0.54 (0.02) | 0.62 (0.02) | 0.47 (0.02) | 0.59 (0.02) | 0.57 (0.02) | 0.49 (0.02) | 0.59 (0.02) |
| Impedance | 0.24 (0.01) | 0.33 (0.01) | 0.29 (0.01) | 0.28 (0.01) | 0.29 (0.01) | 0.33 (0.01) | 0.24 (0.01) | 0.31 (0.01) | 0.31 (0.01) | 0.27 (0.01) | 0.30 (0.01) |
| Neuroticism Score | 0.09 (0.00) | 0.14 (0.01) | 0.12 (0.00) | 0.11 (0.00) | 0.11 (0.00) | 0.14 (0.01) | 0.10 (0.00) | 0.12 (0.00) | 0.12 (0.00) | 0.11 (0.01) | 0.12 (0.00) |
| Pulse | 0.13 (0.01) | 0.19 (0.01) | 0.17 (0.01) | 0.16 (0.01) | 0.17 (0.01) | 0.20 (0.01) | 0.14 (0.01) | 0.18 (0.01) | 0.18 (0.01) | 0.16 (0.01) | 0.17 (0.01) |
| Reaction Time | 0.07 (0.00) | 0.11 (0.01) | 0.09 (0.00) | 0.09 (0.00) | 0.09 (0.00) | 0.11 (0.01) | 0.08 (0.00) | 0.10 (0.00) | 0.10 (0.00) | 0.09 (0.00) | 0.10 (0.00) |
| Systolic Blood Pressure | 0.13 (0.01) | 0.19 (0.01) | 0.16 (0.01) | 0.15 (0.01) | 0.16 (0.01) | 0.19 (0.01) | 0.14 (0.00) | 0.17 (0.01) | 0.17 (0.01) | 0.16 (0.01) | 0.17 (0.01) |
| College Education | 0.16 (0.01) | 0.24 (0.01) | 0.20 (0.01) | 0.19 (0.01) | 0.19 (0.01) | 0.23 (0.01) | 0.17 (0.00) | 0.22 (0.01) | 0.21 (0.01) | 0.18 (0.01) | 0.21 (0.01) |
| Ever Smoked | 0.06 (0.00) | 0.10 (0.00) | 0.08 (0.00) | 0.07 (0.00) | 0.07 (0.00) | 0.10 (0.00) | 0.07 (0.00) | 0.08 (0.00) | 0.08 (0.00) | 0.08 (0.00) | 0.08 (0.00) |
| Hypertension | 0.09 (0.00) | 0.14 (0.01) | 0.11 (0.00) | 0.11 (0.00) | 0.12 (0.00) | 0.14 (0.01) | 0.10 (0.00) | 0.12 (0.01) | 0.13 (0.01) | 0.11 (0.01) | 0.12 (0.01) |
| Snorer | 0.05 (0.00) | 0.08 (0.00) | 0.06 (0.00) | 0.06 (0.00) | 0.06 (0.00) | 0.08 (0.00) | 0.05 (0.00) | 0.07 (0.00) | 0.07 (0.00) | 0.06 (0.00) | 0.07 (0.00) |
| Difficulty Falling Asleep | 0.05 (0.00) | 0.08 (0.00) | 0.07 (0.00) | 0.07 (0.00) | 0.07 (0.00) | 0.08 (0.00) | 0.06 (0.00) | 0.07 (0.00) | 0.07 (0.00) | 0.06 (0.00) | 0.07 (0.00) |
| Preference for Evenings | 0.10 (0.00) | 0.16 (0.01) | 0.13 (0.00) | 0.13 (0.00) | 0.13 (0.00) | 0.15 (0.01) | 0.11 (0.00) | 0.14 (0.00) | 0.14 (0.00) | 0.12 (0.01) | 0.14 (0.00) |
| Average | 0.09 (0.00) | 0.14 (0.00) | 0.12 (0.00) | 0.11 (0.00) | 0.12 (0.00) | 0.14 (0.00) | 0.10 (0.00) | 0.13 (0.00) | 0.13 (0.00) | 0.11 (0.00) | 0.13 (0.00) |
| Relative to GCTA | 1 | 1.46 (0.01) | 1.24 (0.01) | 1.19 (0.01) | 1.21 (0.01) | 1.44 (0.01) | 1.04 (0.01) | 1.31 (0.01) | 1.31 (0.01) | 1.15 (0.01) | 1.31 (0.01) |
| Relative to LDAK | 0.68 (0.01) | 1 | 0.85 (0.01) | 0.82 (0.01) | 0.83 (0.01) | 0.99 (0.01) | 0.71 (0.01) | 0.90 (0.01) | 0.90 (0.01) | 0.79 (0.01) | 0.90 (0.01) |
| Relative to BLD-LDAK | 0.76 (0.01) | 1.11 (0.01) | 0.94 (0.01) | 0.90 (0.01) | 0.92 (0.01) | 1.10 (0.01) | 0.79 (0.01) | 1 | 0.99 (0.01) | 0.88 (0.01) | 0.99 (0.01) |
| Relative to Baseline LD | 0.86 (0.01) | 1.26 (0.01) | 1.07 (0.01) | 1.03 (0.01) | 1.04 (0.01) | 1.25 (0.01) | 0.90 (0.01) | 1.14 (0.01) | 1.13 (0.01) | 1 | 1.13 (0.01) |
| Rel. to UKBb Panel estimates | 1.04 (0.01) | 1.04 (0.01) | 1.03 (0.01) | 1.02 (0.01) | 1.02 (0.01) | 1.04 (0.01) | 1.04 (0.01) | 1.05 (0.01) | 1.04 (0.01) | 1.03 (0.01) | 1.05 (0.01) |

| Heritability Model | GCTA | LDAK | LDAK -Thin | GCTA -LDMS-R | GCTA -LDMS-I | LDAK +24Fun | Baseline | BLD-LDAK | BLD-LDAK +Alpha | Baseline LD | BLD-LDAK -LITE |
|---|---|---|---|---|---|---|---|---|---|---|---|
| Coronary Artery | 0.05 (0.01) | 0.14 (0.02) | 0.09 (0.01) | 0.07 (0.01) | 0.07 (0.01) | 0.16 (0.02) | 0.08 (0.01) | 0.10 (0.02) | 0.10 (0.02) | 0.09 (0.02) | 0.10 (0.02) |
| Crohn’s Disease | 0.33 (0.05) | 0.91 (0.10) | 0.54 (0.06) | 0.54 (0.06) | 0.63 (0.09) | 0.96 (0.09) | 0.53 (0.04) | 0.60 (0.07) | 0.56 (0.06) | 0.57 (0.06) | 0.60 (0.07) |
| Ever Smoked? | 0.05 (0.01) | 0.14 (0.01) | 0.07 (0.01) | 0.07 (0.01) | 0.09 (0.01) | 0.14 (0.01) | 0.06 (0.01) | 0.13 (0.01) | 0.12 (0.01) | 0.11 (0.01) | 0.12 (0.02) |
| Inflammatory Bowel | 0.23 (0.03) | 0.66 (0.06) | 0.37 (0.03) | 0.36 (0.04) | 0.40 (0.05) | 0.67 (0.05) | 0.36 (0.03) | 0.47 (0.04) | 0.44 (0.04) | 0.41 (0.04) | 0.47 (0.04) |
| Rheumatoid Arthritis | 0.10 (0.02) | 0.28 (0.04) | 0.15 (0.02) | 0.14 (0.02) | 0.13 (0.02) | 0.28 (0.03) | 0.15 (0.01) | 0.16 (0.02) | 0.16 (0.02) | 0.15 (0.02) | 0.15 (0.02) |
| Schizophrenia | 0.35 (0.02) | 0.73 (0.02) | 0.47 (0.02) | 0.42 (0.02) | 0.44 (0.02) | 0.71 (0.02) | 0.39 (0.01) | 0.49 (0.02) | 0.47 (0.02) | 0.44 (0.02) | 0.50 (0.02) |
| Type 2 Diabetes | 0.05 (0.00) | 0.14 (0.01) | 0.09 (0.01) | 0.07 (0.01) | 0.07 (0.01) | 0.14 (0.01) | 0.07 (0.01) | 0.10 (0.01) | 0.09 (0.01) | 0.09 (0.01) | 0.10 (0.01) |
| Bone Mineral Density | 0.12 (0.02) | 0.34 (0.04) | 0.20 (0.02) | 0.16 (0.02) | 0.17 (0.03) | 0.35 (0.04) | 0.20 (0.02) | 0.22 (0.03) | 0.19 (0.03) | 0.22 (0.04) | 0.22 (0.03) |
| Body Mass Index | 0.11 (0.01) | 0.23 (0.01) | 0.15 (0.01) | 0.13 (0.01) | 0.13 (0.01) | 0.22 (0.01) | 0.12 (0.01) | 0.16 (0.01) | 0.16 (0.01) | 0.15 (0.01) | 0.16 (0.01) |
| Depressive Symptoms | 0.03 (0.00) | 0.08 (0.01) | 0.05 (0.00) | 0.04 (0.00) | 0.04 (0.01) | 0.08 (0.01) | 0.04 (0.00) | 0.04 (0.01) | 0.04 (0.01) | 0.04 (0.01) | 0.05 (0.01) |
| Height | 0.27 (0.02) | 0.56 (0.03) | 0.37 (0.02) | 0.32 (0.02) | 0.33 (0.02) | 0.55 (0.03) | 0.33 (0.01) | 0.37 (0.02) | 0.37 (0.02) | 0.32 (0.02) | 0.38 (0.02) |
| Menarche Age | 0.17 (0.01) | 0.32 (0.01) | 0.22 (0.01) | 0.19 (0.01) | 0.20 (0.01) | 0.31 (0.01) | 0.18 (0.01) | 0.22 (0.01) | 0.21 (0.01) | 0.21 (0.01) | 0.22 (0.01) |
| Menopause Age | 0.09 (0.01) | 0.26 (0.03) | 0.15 (0.02) | 0.13 (0.02) | 0.12 (0.02) | 0.24 (0.03) | 0.12 (0.01) | 0.14 (0.03) | 0.14 (0.02) | 0.11 (0.03) | 0.13 (0.03) |
| Neuroticism | 0.07 (0.01) | 0.16 (0.02) | 0.09 (0.01) | 0.08 (0.00) | 0.08 (0.01) | 0.16 (0.02) | 0.08 (0.01) | 0.09 (0.01) | 0.09 (0.01) | 0.08 (0.01) | 0.10 (0.01) |
| Subjective Well-Being | 0.02 (0.00) | 0.04 (0.00) | 0.03 (0.00) | 0.02 (0.00) | 0.02 (0.00) | 0.04 (0.00) | 0.02 (0.00) | 0.02 (0.00) | 0.02 (0.00) | 0.02 (0.00) | 0.02 (0.00) |
| Waist-Hip Ratio | 0.07 (0.01) | 0.18 (0.01) | 0.10 (0.01) | 0.09 (0.01) | 0.09 (0.01) | 0.18 (0.01) | 0.09 (0.01) | 0.12 (0.01) | 0.11 (0.01) | 0.11 (0.01) | 0.13 (0.01) |
| Years Education | 0.09 (0.00) | 0.18 (0.01) | 0.12 (0.00) | 0.10 (0.00) | 0.11 (0.00) | 0.18 (0.01) | 0.10 (0.00) | 0.12 (0.01) | 0.12 (0.00) | 0.11 (0.01) | 0.13 (0.01) |
| Average | 0.05 (0.00) | 0.14 (0.00) | 0.07 (0.00) | 0.06 (0.00) | 0.08 (0.00) | 0.14 (0.00) | 0.07 (0.00) | 0.11 (0.00) | 0.11 (0.00) | 0.09 (0.00) | 0.10 (0.00) |
| Relative to GCTA | 1 | 2.18 (0.03) | 1.38 (0.02) | 1.20 (0.02) | 1.24 (0.02) | 2.13 (0.03) | 1.18 (0.02) | 1.43 (0.02) | 1.39 (0.02) | 1.32 (0.03) | 1.47 (0.03) |
| Relative to LDAK | 0.44 (0.01) | 1 | 0.62 (0.01) | 0.54 (0.01) | 0.57 (0.01) | 0.98 (0.01) | 0.53 (0.01) | 0.66 (0.01) | 0.64 (0.01) | 0.61 (0.01) | 0.68 (0.01) |
| Relative to BLD-LDAK | 0.65 (0.01) | 1.49 (0.02) | 0.93 (0.01) | 0.82 (0.01) | 0.86 (0.02) | 1.46 (0.02) | 0.80 (0.01) | 1 | 0.97 (0.02) | 0.91 (0.02) | 1.03 (0.02) |
| Relative to Baseline LD | 0.71 (0.01) | 1.62 (0.03) | 1.02 (0.02) | 0.90 (0.01) | 0.94 (0.02) | 1.59 (0.02) | 0.88 (0.01) | 1.09 (0.02) | 1.05 (0.02) | 1 | 1.12 (0.02) |
| Rel. to UKBb Panel estimates | 1.02 (0.02) | 1.05 (0.02) | 1.00 (0.02) | 0.99 (0.02) | 0.98 (0.02) | 1.05 (0.02) | 1.03 (0.01) | 1.05 (0.02) | 1.05 (0.02) | 1.07 (0.02) | 1.06 (0.02) |

**Supplementary Table 3: Estimates of SNP heritability.** We report estimates of the heritability explained by SNPs with MAF>0.05. All estimates are on the observed scale. The final column shows that estimates from the seven-parameter BLD-LDAK-Lite Model are close to those from the 66-parameter BLD-LDAK Model. These estimates were obtained using the 1000 Genome Project reference panel (489 European individuals); the final row of each table compares these estimates to those obtained using a UK Biobank reference panel (2 000 individuals randomly picked from the 130 k we used for the UKBb GWAS); for example, when assuming the BLD-LDAK Model, estimates for the 14 UKBb GWAS obtained using the 1000 Genome Project panel are on average 5% (s.d. 1%) higher than those obtained using the UK Biobank panel. We expect estimates using the UK Biobank panel to be more accurate (both due to its larger size and because it is more closely matched ancestrally). Therefore, our results indicate that using the 1000 Genome Project panel leads to a small over-estimation of SNP heritability (however, Supplementary Table 1 shows that this over-estimation does not affect the ranking of heritability models).

| Heritability Model | GCTA | LDAK | LDAK -Thin | GCTA -LDMS-R | GCTA -LDMS-I | LDAK +24Fun | Baseline | BLD-LDAK | BLD-LDAK +Alpha | Baseline LD | BLD-LDAK -LITE |
|---|---|---|---|---|---|---|---|---|---|---|---|
| Body Mass Index | 1.06 (0.01) | 0.91 (0.01) | 1.03 (0.01) | 1.06 (0.01) | 1.05 (0.01) | 0.91 (0.01) | 1.03 (0.01) | 1.03 (0.01) | 1.04 (0.01) | 1.00 (0.02) | 1.02 (0.01) |
| Forced Vital Capacity | 1.08 (0.01) | 0.91 (0.01) | 1.04 (0.01) | 1.07 (0.01) | 1.05 (0.01) | 0.91 (0.01) | 1.03 (0.01) | 1.02 (0.01) | 1.03 (0.01) | 1.02 (0.02) | 1.02 (0.01) |
| Height | 1.13 (0.01) | 0.85 (0.02) | 1.07 (0.01) | 1.13 (0.01) | 1.10 (0.01) | 0.87 (0.02) | 1.05 (0.01) | 1.07 (0.01) | 1.07 (0.01) | 1.04 (0.03) | 1.06 (0.02) |
| Impedance | 1.06 (0.01) | 0.89 (0.01) | 1.03 (0.01) | 1.06 (0.01) | 1.04 (0.01) | 0.89 (0.01) | 1.02 (0.01) | 1.02 (0.01) | 1.03 (0.01) | 0.99 (0.02) | 1.02 (0.01) |
| Neuroticism Score | 1.03 (0.01) | 0.96 (0.01) | 1.02 (0.01) | 1.03 (0.01) | 1.03 (0.01) | 0.97 (0.01) | 1.02 (0.01) | 1.02 (0.01) | 1.03 (0.01) | 1.00 (0.01) | 1.02 (0.01) |
| Pulse | 1.05 (0.01) | 0.93 (0.01) | 1.01 (0.01) | 1.04 (0.01) | 1.03 (0.01) | 0.93 (0.01) | 1.01 (0.01) | 1.01 (0.01) | 1.00 (0.01) | 0.99 (0.02) | 1.00 (0.01) |
| Reaction Time | 1.03 (0.01) | 0.96 (0.01) | 1.01 (0.01) | 1.02 (0.01) | 1.02 (0.01) | 0.97 (0.01) | 1.01 (0.01) | 1.01 (0.01) | 1.02 (0.01) | 1.01 (0.01) | 1.00 (0.01) |
| Systolic Blood Pressure | 1.04 (0.01) | 0.94 (0.01) | 1.02 (0.01) | 1.03 (0.01) | 1.02 (0.01) | 0.93 (0.01) | 1.00 (0.01) | 1.01 (0.01) | 1.01 (0.01) | 1.01 (0.01) | 1.01 (0.01) |
| College Education | 1.07 (0.01) | 0.96 (0.01) | 1.05 (0.01) | 1.06 (0.01) | 1.05 (0.01) | 0.97 (0.01) | 1.05 (0.01) | 1.05 (0.01) | 1.05 (0.01) | 1.03 (0.01) | 1.04 (0.01) |
| Ever Smoked | 1.02 (0.00) | 0.97 (0.01) | 1.01 (0.00) | 1.02 (0.00) | 1.03 (0.01) | 0.97 (0.01) | 1.01 (0.00) | 1.01 (0.01) | 1.02 (0.01) | 1.00 (0.01) | 1.01 (0.01) |
| Hypertension | 1.04 (0.01) | 0.96 (0.01) | 1.02 (0.01) | 1.03 (0.01) | 1.02 (0.01) | 0.95 (0.01) | 1.01 (0.01) | 1.01 (0.01) | 1.01 (0.01) | 1.01 (0.01) | 1.00 (0.01) |
| Snorer | 1.02 (0.00) | 0.98 (0.01) | 1.01 (0.00) | 1.02 (0.00) | 1.01 (0.01) | 0.98 (0.01) | 1.01 (0.00) | 1.01 (0.01) | 1.01 (0.01) | 1.02 (0.01) | 1.00 (0.01) |
| Difficulty Falling Asleep | 1.01 (0.00) | 0.96 (0.01) | 1.00 (0.00) | 1.01 (0.00) | 1.00 (0.01) | 0.97 (0.01) | 1.01 (0.00) | 1.00 (0.01) | 1.01 (0.01) | 1.00 (0.01) | 1.00 (0.01) |
| Preference for Evenings | 1.03 (0.00) | 0.95 (0.01) | 1.01 (0.00) | 1.03 (0.01) | 1.02 (0.01) | 0.95 (0.01) | 1.02 (0.01) | 1.01 (0.01) | 1.01 (0.01) | 0.99 (0.01) | 1.00 (0.01) |
| Average | 1.04 (0.00) | 0.95 (0.00) | 1.02 (0.00) | 1.03 (0.00) | 1.03 (0.00) | 0.95 (0.00) | 1.02 (0.00) | 1.02 (0.00) | 1.02 (0.00) | 1.01 (0.00) | 1.01 (0.00) |
| Relative to GCTA | 1 | 0.92 (0.00) | 0.98 (0.00) | 1.00 (0.00) | 0.99 (0.00) | 0.91 (0.00) | 0.98 (0.00) | 0.98 (0.00) | 0.98 (0.00) | 0.97 (0.00) | 0.97 (0.00) |
| Relative to LDAK | 1.09 (0.00) | 1 | 1.07 (0.00) | 1.09 (0.00) | 1.08 (0.00) | 1.00 (0.00) | 1.07 (0.00) | 1.07 (0.00) | 1.08 (0.00) | 1.06 (0.00) | 1.06 (0.00) |
| Relative to BLD-LDAK | 1.02 (0.00) | 0.94 (0.00) | 1.00 (0.00) | 1.02 (0.00) | 1.01 (0.00) | 0.93 (0.00) | 1.00 (0.00) | 1 | 1.00 (0.00) | 0.99 (0.00) | 1.00 (0.00) |
| Relative to Baseline LD | 1.03 (0.00) | 0.94 (0.00) | 1.01 (0.00) | 1.03 (0.00) | 1.02 (0.00) | 0.94 (0.00) | 1.01 (0.00) | 1.01 (0.00) | 1.01 (0.00) | 1 | 1.00 (0.00) |
| Rel. to UKBb Panel estimates | 1.00 (0.00) | 1.02 (0.00) | 1.01 (0.00) | 1.01 (0.00) | 1.01 (0.00) | 1.02 (0.00) | 1.01 (0.00) | 1.01 (0.00) | 1.01 (0.00) | 1.00 (0.00) | 1.01 (0.00) |

| Heritability Model | GCTA | LDAK | LDAK -Thin | GCTA -LDMS-R | GCTA -LDMS-I | LDAK +24Fun | Baseline | BLD-LDAK | BLD-LDAK +Alpha | Baseline LD | BLD-LDAK -LITE |
|---|---|---|---|---|---|---|---|---|---|---|---|
| Coronary Artery | 1.04 (0.01) | 0.99 (0.01) | 1.01 (0.01) | 1.03 (0.01) | 1.03 (0.01) | 0.99 (0.01) | 1.01 (0.01) | 1.01 (0.01) | 1.01 (0.01) | 1.01 (0.02) | 1.01 (0.01) |
| Crohn’s Disease | 1.05 (0.01) | 0.97 (0.01) | 1.02 (0.01) | 1.01 (0.01) | 1.00 (0.01) | 0.96 (0.01) | 1.00 (0.01) | 1.01 (0.01) | 1.02 (0.01) | 1.00 (0.01) | 1.01 (0.01) |
| Ever Smoked? | 1.02 (0.01) | 0.96 (0.01) | 1.00 (0.01) | 1.00 (0.01) | 0.99 (0.01) | 0.97 (0.01) | 1.01 (0.01) | 0.97 (0.01) | 0.97 (0.01) | 0.97 (0.01) | 0.97 (0.01) |
| Inflammatory Bowel | 1.08 (0.01) | 0.98 (0.01) | 1.04 (0.01) | 1.04 (0.01) | 1.03 (0.02) | 0.97 (0.01) | 1.03 (0.01) | 1.01 (0.01) | 1.03 (0.01) | 1.02 (0.01) | 1.02 (0.01) |
| Rheumatoid Arthritis | 0.97 (0.01) | 0.90 (0.01) | 0.95 (0.01) | 0.95 (0.01) | 0.96 (0.01) | 0.89 (0.01) | 0.93 (0.01) | 0.94 (0.01) | 0.94 (0.01) | 0.94 (0.02) | 0.95 (0.01) |
| Schizophrenia | 1.10 (0.01) | 0.91 (0.01) | 1.05 (0.01) | 1.07 (0.01) | 1.05 (0.01) | 0.92 (0.01) | 1.06 (0.01) | 1.03 (0.01) | 1.04 (0.01) | 1.03 (0.02) | 1.02 (0.01) |
| Type 2 Diabetes | 1.04 (0.01) | 0.95 (0.01) | 0.99 (0.01) | 1.01 (0.01) | 1.01 (0.01) | 0.95 (0.01) | 1.00 (0.01) | 0.98 (0.01) | 0.99 (0.01) | 0.97 (0.02) | 0.98 (0.01) |
| Bone Mineral Density | 1.05 (0.01) | 1.00 (0.01) | 1.03 (0.01) | 1.04 (0.01) | 1.04 (0.01) | 0.99 (0.01) | 1.02 (0.01) | 1.02 (0.01) | 1.03 (0.01) | 1.01 (0.01) | 1.02 (0.01) |
| Body Mass Index | 0.74 (0.01) | 0.56 (0.01) | 0.68 (0.01) | 0.71 (0.01) | 0.70 (0.01) | 0.57 (0.01) | 0.69 (0.01) | 0.66 (0.01) | 0.66 (0.01) | 0.63 (0.02) | 0.65 (0.01) |
| Depressive Symptoms | 1.02 (0.01) | 0.96 (0.01) | 1.01 (0.01) | 1.01 (0.01) | 1.01 (0.01) | 0.97 (0.01) | 1.01 (0.01) | 1.01 (0.01) | 1.02 (0.01) | 1.02 (0.02) | 1.00 (0.01) |
| Height | 1.41 (0.04) | 0.95 (0.04) | 1.24 (0.03) | 1.34 (0.03) | 1.31 (0.05) | 1.00 (0.03) | 1.20 (0.02) | 1.23 (0.02) | 1.24 (0.02) | 1.26 (0.04) | 1.24 (0.03) |
| Menarche Age | 1.09 (0.01) | 0.88 (0.02) | 1.04 (0.01) | 1.08 (0.01) | 1.06 (0.02) | 0.90 (0.02) | 1.05 (0.01) | 1.03 (0.01) | 1.04 (0.01) | 0.99 (0.02) | 1.02 (0.01) |
| Menopause Age | 1.02 (0.02) | 0.94 (0.02) | 0.99 (0.01) | 1.00 (0.01) | 1.01 (0.02) | 0.95 (0.02) | 1.00 (0.01) | 0.99 (0.02) | 1.00 (0.01) | 1.01 (0.02) | 1.00 (0.02) |
| Neuroticism | 1.03 (0.01) | 0.91 (0.02) | 1.00 (0.01) | 1.02 (0.01) | 1.02 (0.01) | 0.91 (0.02) | 1.00 (0.01) | 0.99 (0.01) | 1.00 (0.01) | 1.01 (0.02) | 0.97 (0.02) |
| Subjective Well-Being | 1.02 (0.01) | 0.97 (0.01) | 1.01 (0.01) | 1.02 (0.01) | 1.02 (0.01) | 0.98 (0.01) | 1.01 (0.01) | 1.02 (0.01) | 1.02 (0.01) | 1.02 (0.01) | 1.01 (0.01) |
| Waist-Hip Ratio | 0.90 (0.01) | 0.79 (0.01) | 0.86 (0.01) | 0.87 (0.01) | 0.87 (0.01) | 0.79 (0.01) | 0.85 (0.01) | 0.83 (0.01) | 0.84 (0.01) | 0.83 (0.02) | 0.82 (0.01) |
| Years Education | 1.00 (0.01) | 0.80 (0.02) | 0.95 (0.01) | 0.98 (0.01) | 0.96 (0.01) | 0.81 (0.02) | 0.96 (0.01) | 0.92 (0.01) | 0.94 (0.01) | 0.93 (0.02) | 0.91 (0.01) |
| Average | 1.01 (0.00) | 0.91 (0.00) | 0.98 (0.00) | 1.00 (0.00) | 0.99 (0.00) | 0.92 (0.00) | 0.98 (0.00) | 0.97 (0.00) | 0.98 (0.00) | 0.97 (0.00) | 0.97 (0.00) |
| Relative to GCTA | 1 | 0.90 (0.00) | 0.97 (0.00) | 0.98 (0.00) | 0.98 (0.00) | 0.90 (0.00) | 0.97 (0.00) | 0.95 (0.00) | 0.96 (0.00) | 0.95 (0.00) | 0.95 (0.00) |
| Relative to LDAK | 1.09 (0.00) | 1 | 1.06 (0.00) | 1.08 (0.00) | 1.08 (0.00) | 1.00 (0.00) | 1.07 (0.00) | 1.06 (0.00) | 1.07 (0.00) | 1.06 (0.00) | 1.06 (0.00) |
| Relative to BLD-LDAK | 1.04 (0.00) | 0.95 (0.00) | 1.01 (0.00) | 1.03 (0.00) | 1.02 (0.00) | 0.95 (0.00) | 1.01 (0.00) | 1 | 1.01 (0.00) | 1.00 (0.00) | 1.00 (0.00) |
| Relative to Baseline LD | 1.04 (0.00) | 0.95 (0.00) | 1.01 (0.00) | 1.03 (0.00) | 1.02 (0.00) | 0.95 (0.00) | 1.01 (0.00) | 1.00 (0.00) | 1.01 (0.00) | 1 | 1.00 (0.00) |
| Rel. to UKBb Panel estimates | 1.00 (0.00) | 1.00 (0.00) | 1.01 (0.00) | 1.01 (0.00) | 1.01 (0.00) | 1.00 (0.00) | 1.00 (0.00) | 1.00 (0.00) | 1.00 (0.00) | 0.99 (0.00) | 1.00 (0.00) |

**Supplementary Table 4: Estimates of confounding bias.** We report estimates of confounding bias ($1 + A$, where $A$ is the average additive inflation of test statistics). The final column shows that estimates from the seven-parameter BLD-LDAK-Lite Model are close to those from the 66-parameter BLD-LDAK Model. These estimates were obtained using the 1000 Genome Project reference panel (489 European individuals); the final row of each table compares these estimates to those obtained using a UK Biobank reference panel (2 000 individuals randomly picked from the 130 k we used for the UKBb GWAS). In general, the estimates are very similar; for example, when assuming the BLD-LDAK Model, estimates for the 14 UKBb GWAS obtained using the 1000 Genome Project reference panel are on average only 1% (s.d. 0) higher than those obtained using the UKBb panel.

| Category / Prop. of SNPs | GCTA+1Fun | LDAK+1Fun | LDAK+24Fun | Baseline | BLD-LDAK | BLD-LDAK+Alpha | Baseline LD | BLD-LDAK-Lite+1Fun |
|---|---|---|---|---|---|---|---|---|
| Coding | 0.15 (0.01) | 0.07 (0.00) | 0.05 (0.00) | 0.10 (0.01) | 0.08 (0.01) | 0.08 (0.01) | 0.07 (0.01) | 0.07 (0.01) |
| 1.4% | 10.6 (0.4) | 4.6 (0.2) | 3.7 (0.2) | 6.9 (0.4) | 5.3 (0.4) | 5.3 (0.4) | 4.9 (0.4) | 4.6 (0.4) |
| Conserved | 0.48 (0.01) | 0.12 (0.00) | 0.11 (0.00) | 0.34 (0.01) | 0.26 (0.01) | 0.26 (0.01) | 0.22 (0.01) | 0.29 (0.01) |
| 2.6% | 18.8 (0.4) | 4.7 (0.1) | 4.4 (0.1) | 13.3 (0.4) | 10.0 (0.4) | 10.0 (0.4) | 8.7 (0.4) | 11.1 (0.5) |
| CTCF | 0.22 (0.01) | 0.03 (0.00) | 0.03 (0.00) | 0.02 (0.01) | 0.03 (0.01) | 0.03 (0.01) | 0.02 (0.01) | 0.03 (0.01) |
| 2.4% | 9.2 (0.4) | 1.5 (0.1) | 1.1 (0.1) | 0.8 (0.4) | 1.3 (0.4) | 1.3 (0.4) | 1.0 (0.3) | 1.2 (0.4) |
| DGF | 0.79 (0.01) | 0.26 (0.01) | 0.22 (0.00) | 0.33 (0.02) | 0.30 (0.02) | 0.30 (0.02) | 0.28 (0.02) | 0.37 (0.02) |
| 13.6% | 5.8 (0.1) | 1.9 (0.0) | 1.6 (0.0) | 2.4 (0.2) | 2.2 (0.2) | 2.2 (0.2) | 2.1 (0.2) | 2.7 (0.2) |
| DHS | 0.91 (0.01) | 0.29 (0.01) | 0.26 (0.01) | 0.36 (0.03) | 0.34 (0.03) | 0.34 (0.03) | 0.33 (0.02) | 0.39 (0.03) |
| 16.6% | 5.5 (0.1) | 1.8 (0.0) | 1.6 (0.0) | 2.2 (0.2) | 2.1 (0.2) | 2.1 (0.2) | 2.0 (0.1) | 2.4 (0.2) |
| FANTOM5 Enhancer | 0.07 (0.00) | 0.01 (0.00) | 0.00 (0.00) | 0.01 (0.00) | 0.01 (0.00) | 0.01 (0.00) | 0.01 (0.00) | 0.01 (0.00) |
| 0.4% | 17.3 (1.0) | 1.6 (0.2) | 1.1 (0.2) | 1.6 (0.9) | 3.1 (0.9) | 3.1 (0.9) | 2.8 (0.9) | 3.3 (0.9) |
| Enhancer | 0.29 (0.01) | 0.09 (0.00) | 0.07 (0.00) | 0.11 (0.01) | 0.11 (0.01) | 0.11 (0.01) | 0.10 (0.01) | 0.11 (0.01) |
| 4.2% | 7.0 (0.2) | 2.2 (0.1) | 1.6 (0.1) | 2.7 (0.2) | 2.7 (0.2) | 2.7 (0.2) | 2.3 (0.2) | 2.7 (0.2) |
| Fetal DHS | 0.71 (0.01) | 0.16 (0.00) | 0.14 (0.00) | 0.24 (0.02) | 0.23 (0.02) | 0.23 (0.02) | 0.20 (0.02) | 0.27 (0.02) |
| 8.4% | 8.4 (0.1) | 1.9 (0.1) | 1.7 (0.0) | 2.8 (0.2) | 2.8 (0.2) | 2.8 (0.2) | 2.4 (0.2) | 3.2 (0.2) |
| H3K27ac (Hnisz) | 0.70 (0.01) | 0.58 (0.01) | 0.55 (0.01) | 0.59 (0.01) | 0.56 (0.01) | 0.56 (0.01) | 0.52 (0.01) | 0.55 (0.01) |
| 38.9% | 1.8 (0.0) | 1.5 (0.0) | 1.4 (0.0) | 1.5 (0.0) | 1.4 (0.0) | 1.4 (0.0) | 1.3 (0.0) | 1.4 (0.0) |
| H3K27ac (PGC2) | 0.66 (0.01) | 0.44 (0.01) | 0.40 (0.01) | 0.45 (0.02) | 0.43 (0.01) | 0.43 (0.01) | 0.42 (0.01) | 0.43 (0.01) |
| 26.9% | 2.4 (0.0) | 1.6 (0.0) | 1.5 (0.0) | 1.7 (0.1) | 1.6 (0.1) | 1.6 (0.1) | 1.6 (0.1) | 1.6 (0.1) |
| H3K4me1 | 0.87 (0.01) | 0.67 (0.01) | 0.62 (0.01) | 0.76 (0.02) | 0.70 (0.02) | 0.70 (0.02) | 0.67 (0.02) | 0.69 (0.02) |
| 42.4% | 2.0 (0.0) | 1.6 (0.0) | 1.5 (0.0) | 1.8 (0.0) | 1.6 (0.0) | 1.6 (0.0) | 1.6 (0.0) | 1.6 (0.0) |
| H3K4me3 | 0.47 (0.01) | 0.26 (0.01) | 0.23 (0.01) | 0.30 (0.01) | 0.29 (0.01) | 0.29 (0.01) | 0.28 (0.01) | 0.27 (0.01) |
| 13.3% | 3.5 (0.1) | 2.0 (0.0) | 1.7 (0.0) | 2.3 (0.1) | 2.2 (0.1) | 2.2 (0.1) | 2.1 (0.1) | 2.1 (0.1) |
| H3K9ac | 0.48 (0.01) | 0.28 (0.01) | 0.26 (0.01) | 0.39 (0.01) | 0.36 (0.01) | 0.36 (0.01) | 0.33 (0.01) | 0.35 (0.01) |
| 12.5% | 3.8 (0.1) | 2.3 (0.0) | 2.1 (0.0) | 3.1 (0.1) | 2.8 (0.1) | 2.8 (0.1) | 2.7 (0.1) | 2.8 (0.1) |
| Intronic | 0.55 (0.01) | 0.49 (0.01) | 0.47 (0.01) | 0.45 (0.01) | 0.43 (0.01) | 0.43 (0.01) | 0.43 (0.01) | 0.45 (0.01) |
| 38.7% | 1.4 (0.0) | 1.3 (0.0) | 1.2 (0.0) | 1.1 (0.0) | 1.1 (0.0) | 1.1 (0.0) | 1.1 (0.0) | 1.2 (0.0) |
| Promoter Flanking | 0.12 (0.01) | 0.02 (0.00) | 0.02 (0.00) | -0.00 (0.01) | -0.00 (0.01) | -0.00 (0.01) | -0.01 (0.00) | -0.00 (0.01) |
| 0.8% | 15.0 (0.7) | 2.7 (0.2) | 1.8 (0.2) | -0.6 (0.7) | -0.2 (0.6) | -0.2 (0.6) | -0.6 (0.6) | -0.5 (0.7) |
| Promoter | 0.19 (0.01) | 0.10 (0.00) | 0.08 (0.00) | 0.10 (0.01) | 0.08 (0.01) | 0.08 (0.01) | 0.07 (0.01) | 0.07 (0.01) |
| 4.6% | 4.1 (0.2) | 2.2 (0.1) | 1.7 (0.1) | 2.1 (0.2) | 1.8 (0.2) | 1.8 (0.2) | 1.5 (0.2) | 1.4 (0.2) |
| Repressed | 0.12 (0.01) | 0.29 (0.01) | 0.33 (0.01) | 0.29 (0.02) | 0.33 (0.02) | 0.33 (0.02) | 0.33 (0.02) | 0.38 (0.02) |
| 46.1% | 0.3 (0.0) | 0.6 (0.0) | 0.7 (0.0) | 0.6 (0.0) | 0.7 (0.0) | 0.7 (0.0) | 0.7 (0.0) | 0.8 (0.0) |
| Super Enhancer | 0.33 (0.01) | 0.27 (0.00) | 0.26 (0.00) | 0.27 (0.01) | 0.26 (0.00) | 0.26 (0.00) | 0.25 (0.00) | 0.25 (0.01) |
| 16.7% | 2.0 (0.0) | 1.6 (0.0) | 1.5 (0.0) | 1.6 (0.0) | 1.5 (0.0) | 1.5 (0.0) | 1.5 (0.0) | 1.5 (0.0) |
| T. Factor Binding Site | 0.65 (0.01) | 0.24 (0.01) | 0.21 (0.00) | 0.31 (0.02) | 0.28 (0.02) | 0.28 (0.02) | 0.24 (0.02) | 0.32 (0.02) |
| 13.1% | 5.0 (0.1) | 1.8 (0.0) | 1.6 (0.0) | 2.3 (0.1) | 2.2 (0.1) | 2.2 (0.1) | 1.8 (0.1) | 2.4 (0.1) |
| Transcribed | 0.58 (0.01) | 0.47 (0.01) | 0.44 (0.01) | 0.40 (0.02) | 0.39 (0.02) | 0.39 (0.02) | 0.38 (0.02) | 0.37 (0.01) |
| 34.6% | 1.7 (0.0) | 1.4 (0.0) | 1.3 (0.0) | 1.2 (0.1) | 1.1 (0.0) | 1.1 (0.0) | 1.1 (0.0) | 1.1 (0.0) |
| Transcription Start Site | 0.18 (0.01) | 0.07 (0.00) | 0.05 (0.00) | 0.08 (0.01) | 0.06 (0.01) | 0.06 (0.01) | 0.06 (0.01) | 0.06 (0.01) |
| 1.8% | 10.0 (0.3) | 3.7 (0.1) | 2.8 (0.1) | 4.5 (0.4) | 3.5 (0.4) | 3.5 (0.4) | 3.3 (0.3) | 3.3 (0.4) |
| 3'UTR | 0.12 (0.01) | 0.03 (0.00) | 0.03 (0.00) | 0.05 (0.00) | 0.03 (0.00) | 0.03 (0.00) | 0.03 (0.00) | 0.02 (0.00) |
| 1.1% | 10.3 (0.5) | 3.1 (0.2) | 2.3 (0.2) | 4.1 (0.4) | 2.8 (0.4) | 2.8 (0.4) | 3.0 (0.4) | 2.2 (0.4) |
| 5'UTR | 0.08 (0.00) | 0.02 (0.00) | 0.01 (0.00) | 0.03 (0.00) | 0.02 (0.00) | 0.02 (0.00) | 0.02 (0.00) | 0.01 (0.00) |
| 0.5% | 14.8 (0.8) | 3.6 (0.3) | 2.6 (0.2) | 5.6 (0.6) | 4.1 (0.6) | 4.2 (0.6) | 4.3 (0.6) | 2.7 (0.6) |
| Weak Enhancer | 0.26 (0.01) | 0.05 (0.00) | 0.03 (0.00) | 0.05 (0.01) | 0.04 (0.01) | 0.04 (0.01) | 0.03 (0.01) | 0.05 (0.01) |
| 2.1% | 12.4 (0.4) | 2.2 (0.1) | 1.4 (0.1) | 2.2 (0.4) | 1.7 (0.4) | 1.7 (0.4) | 1.5 (0.4) | 2.4 (0.5) |
| Concordance with | 1.000 | 0.998 | 0.999 | 1.000 | 1.000 | 1.000 | 1.000 | 1.000 |
| UKBb Panel estimates | 0.999 | 0.985 | 0.992 | 1.000 | 0.999 | 0.999 | 0.999 | 0.998 |

**Supplementary Table 5: Estimates of average proportions of SNP heritability and enrichments from 14 UKBb GWAS.** Pairs of rows report the estimated proportion of SNP heritability and enrichment for each of the 24 functional categories of SNPs in the Baseline Model, averaged across the 14 UKBb GWAS. Enrichments significantly different from one ($P$<0.05/24) are marked in red. The final column shows that estimates from the nine-parameter BLD-LDAK-Lite+1Fun Model are close to those from the 66-parameter BLD-LDAK Model. These estimates were obtained using the 1000 Genome Project reference panel (489 European individuals); the final pair of rows shows that the concordance correlation coefficient[22] between these estimates and those obtained using a UK Biobank reference panel (2 000 individuals randomly picked from the 130 k we used for the UKBb GWAS) is always close to one.

| Category / Prop. of SNPs | GCTA+1Fun | LDAK+1Fun | LDAK+24Fun | Baseline | BLD-LDAK | BLD-LDAK+Alpha | Baseline LD | BLD-LDAK-Lite+1Fun |
|---|---|---|---|---|---|---|---|---|
| Coding | 0.15 (0.01) | 0.07 (0.00) | 0.05 (0.00) | 0.10 (0.01) | 0.08 (0.01) | 0.08 (0.01) | 0.07 (0.01) | 0.07 (0.01) |
| 1.4% | 10.6 (0.4) | 4.6 (0.2) | 3.7 (0.2) | 6.9 (0.4) | 5.3 (0.4) | 5.3 (0.4) | 4.9 (0.4) | 4.6 (0.4) |
| Conserved | 0.48 (0.01) | 0.12 (0.00) | 0.11 (0.00) | 0.34 (0.01) | 0.26 (0.01) | 0.26 (0.01) | 0.22 (0.01) | 0.29 (0.01) |
| 2.6% | 18.8 (0.4) | 4.7 (0.1) | 4.4 (0.1) | 13.3 (0.4) | 10.0 (0.4) | 10.0 (0.4) | 8.7 (0.4) | 11.1 (0.5) |
| CTCF | 0.22 (0.01) | 0.03 (0.00) | 0.03 (0.00) | 0.02 (0.01) | 0.03 (0.01) | 0.03 (0.01) | 0.02 (0.01) | 0.03 (0.01) |
| 2.4% | 9.2 (0.4) | 1.5 (0.1) | 1.1 (0.1) | 0.8 (0.4) | 1.3 (0.4) | 1.3 (0.4) | 1.0 (0.3) | 1.2 (0.4) |
| DGF | 0.79 (0.01) | 0.26 (0.01) | 0.22 (0.00) | 0.33 (0.02) | 0.30 (0.02) | 0.30 (0.02) | 0.28 (0.02) | 0.37 (0.02) |
| 13.6% | 5.8 (0.1) | 1.9 (0.0) | 1.6 (0.0) | 2.4 (0.2) | 2.2 (0.2) | 2.2 (0.2) | 2.1 (0.2) | 2.7 (0.2) |
| DHS | 0.91 (0.01) | 0.29 (0.01) | 0.26 (0.01) | 0.36 (0.03) | 0.34 (0.03) | 0.34 (0.03) | 0.33 (0.02) | 0.39 (0.03) |
| 16.6% | 5.5 (0.1) | 1.8 (0.0) | 1.6 (0.0) | 2.2 (0.2) | 2.1 (0.2) | 2.1 (0.2) | 2.0 (0.1) | 2.4 (0.2) |
| FANTOM5 Enhancer | 0.07 (0.00) | 0.01 (0.00) | 0.00 (0.00) | 0.01 (0.00) | 0.01 (0.00) | 0.01 (0.00) | 0.01 (0.00) | 0.01 (0.00) |
| 0.4% | 17.3 (1.0) | 1.6 (0.2) | 1.1 (0.2) | 1.6 (0.9) | 3.1 (0.9) | 3.1 (0.9) | 2.8 (0.9) | 3.3 (0.9) |
| Enhancer | 0.29 (0.01) | 0.09 (0.00) | 0.07 (0.00) | 0.11 (0.01) | 0.11 (0.01) | 0.11 (0.01) | 0.10 (0.01) | 0.11 (0.01) |
| 4.2% | 7.0 (0.2) | 2.2 (0.1) | 1.6 (0.1) | 2.7 (0.2) | 2.7 (0.2) | 2.7 (0.2) | 2.3 (0.2) | 2.7 (0.2) |
| Fetal DHS | 0.71 (0.01) | 0.16 (0.00) | 0.14 (0.00) | 0.24 (0.02) | 0.23 (0.02) | 0.23 (0.02) | 0.20 (0.02) | 0.27 (0.02) |
| 8.4% | 8.4 (0.1) | 1.9 (0.1) | 1.7 (0.0) | 2.8 (0.2) | 2.8 (0.2) | 2.8 (0.2) | 2.4 (0.2) | 3.2 (0.2) |
| H3K27ac (Hnisz) | 0.70 (0.01) | 0.58 (0.01) | 0.55 (0.01) | 0.59 (0.01) | 0.56 (0.01) | 0.56 (0.01) | 0.52 (0.01) | 0.55 (0.01) |
| 38.9% | 1.8 (0.0) | 1.5 (0.0) | 1.4 (0.0) | 1.5 (0.0) | 1.4 (0.0) | 1.4 (0.0) | 1.3 (0.0) | 1.4 (0.0) |
| H3K27ac (PGC2) | 0.66 (0.01) | 0.44 (0.01) | 0.40 (0.01) | 0.45 (0.02) | 0.43 (0.01) | 0.43 (0.01) | 0.42 (0.01) | 0.43 (0.01) |
| 26.9% | 2.4 (0.0) | 1.6 (0.0) | 1.5 (0.0) | 1.7 (0.1) | 1.6 (0.1) | 1.6 (0.1) | 1.6 (0.1) | 1.6 (0.1) |
| H3K4me1 | 0.87 (0.01) | 0.67 (0.01) | 0.62 (0.01) | 0.76 (0.02) | 0.70 (0.02) | 0.70 (0.02) | 0.67 (0.02) | 0.69 (0.02) |
| 42.4% | 2.0 (0.0) | 1.6 (0.0) | 1.5 (0.0) | 1.8 (0.0) | 1.6 (0.0) | 1.6 (0.0) | 1.6 (0.0) | 1.6 (0.0) |
| H3K4me3 | 0.47 (0.01) | 0.26 (0.01) | 0.23 (0.01) | 0.30 (0.01) | 0.29 (0.01) | 0.29 (0.01) | 0.28 (0.01) | 0.27 (0.01) |
| 13.3% | 3.5 (0.1) | 2.0 (0.0) | 1.7 (0.0) | 2.3 (0.1) | 2.2 (0.1) | 2.2 (0.1) | 2.1 (0.1) | 2.1 (0.1) |
| H3K9ac | 0.48 (0.01) | 0.28 (0.01) | 0.26 (0.01) | 0.39 (0.01) | 0.36 (0.01) | 0.36 (0.01) | 0.33 (0.01) | 0.35 (0.01) |
| 12.5% | 3.8 (0.1) | 2.3 (0.0) | 2.1 (0.0) | 3.1 (0.1) | 2.8 (0.1) | 2.8 (0.1) | 2.7 (0.1) | 2.8 (0.1) |
| Intronic | 0.55 (0.01) | 0.49 (0.01) | 0.47 (0.01) | 0.45 (0.01) | 0.43 (0.01) | 0.43 (0.01) | 0.43 (0.01) | 0.45 (0.01) |
| 38.7% | 1.4 (0.0) | 1.3 (0.0) | 1.2 (0.0) | 1.1 (0.0) | 1.1 (0.0) | 1.1 (0.0) | 1.1 (0.0) | 1.2 (0.0) |
| Promoter Flanking | 0.12 (0.01) | 0.02 (0.00) | 0.02 (0.00) | -0.00 (0.01) | -0.00 (0.01) | -0.00 (0.01) | -0.01 (0.00) | -0.00 (0.01) |
| 0.8% | 15.0 (0.7) | 2.7 (0.2) | 1.8 (0.2) | -0.6 (0.7) | -0.2 (0.6) | -0.2 (0.6) | -0.6 (0.6) | -0.5 (0.7) |
| Promoter | 0.19 (0.01) | 0.10 (0.00) | 0.08 (0.00) | 0.10 (0.01) | 0.08 (0.01) | 0.08 (0.01) | 0.07 (0.01) | 0.07 (0.01) |
| 4.6% | 4.1 (0.2) | 2.2 (0.1) | 1.7 (0.1) | 2.1 (0.2) | 1.8 (0.2) | 1.8 (0.2) | 1.5 (0.2) | 1.4 (0.2) |
| Repressed | 0.12 (0.01) | 0.29 (0.01) | 0.33 (0.01) | 0.29 (0.02) | 0.33 (0.02) | 0.33 (0.02) | 0.33 (0.02) | 0.38 (0.02) |
| 46.1% | 0.3 (0.0) | 0.6 (0.0) | 0.7 (0.0) | 0.6 (0.0) | 0.7 (0.0) | 0.7 (0.0) | 0.7 (0.0) | 0.8 (0.0) |
| Super Enhancer | 0.33 (0.01) | 0.27 (0.00) | 0.26 (0.00) | 0.27 (0.01) | 0.26 (0.00) | 0.26 (0.00) | 0.25 (0.00) | 0.25 (0.01) |
| 16.7% | 2.0 (0.0) | 1.6 (0.0) | 1.5 (0.0) | 1.6 (0.0) | 1.5 (0.0) | 1.5 (0.0) | 1.5 (0.0) | 1.5 (0.0) |
| T. Factor Binding Site | 0.65 (0.01) | 0.24 (0.01) | 0.21 (0.00) | 0.31 (0.02) | 0.28 (0.02) | 0.28 (0.02) | 0.24 (0.02) | 0.32 (0.02) |
| 13.1% | 5.0 (0.1) | 1.8 (0.0) | 1.6 (0.0) | 2.3 (0.1) | 2.2 (0.1) | 2.2 (0.1) | 1.8 (0.1) | 2.4 (0.1) |
| Transcribed | 0.58 (0.01) | 0.47 (0.01) | 0.44 (0.01) | 0.40 (0.02) | 0.39 (0.02) | 0.39 (0.02) | 0.38 (0.02) | 0.37 (0.01) |
| 34.6% | 1.7 (0.0) | 1.4 (0.0) | 1.3 (0.0) | 1.2 (0.1) | 1.1 (0.0) | 1.1 (0.0) | 1.1 (0.0) | 1.1 (0.0) |
| Transcription Start Site | 0.18 (0.01) | 0.07 (0.00) | 0.05 (0.00) | 0.08 (0.01) | 0.06 (0.01) | 0.06 (0.01) | 0.06 (0.01) | 0.06 (0.01) |
| 1.8% | 10.0 (0.3) | 3.7 (0.1) | 2.8 (0.1) | 4.5 (0.4) | 3.5 (0.4) | 3.5 (0.4) | 3.3 (0.3) | 3.3 (0.4) |
| 3'UTR | 0.12 (0.01) | 0.03 (0.00) | 0.03 (0.00) | 0.05 (0.00) | 0.03 (0.00) | 0.03 (0.00) | 0.03 (0.00) | 0.02 (0.00) |
| 1.1% | 10.3 (0.5) | 3.1 (0.2) | 2.3 (0.2) | 4.1 (0.4) | 2.8 (0.4) | 2.8 (0.4) | 3.0 (0.4) | 2.2 (0.4) |
| 5'UTR | 0.08 (0.00) | 0.02 (0.00) | 0.01 (0.00) | 0.03 (0.00) | 0.02 (0.00) | 0.02 (0.00) | 0.02 (0.00) | 0.01 (0.00) |
| 0.5% | 14.8 (0.8) | 3.6 (0.3) | 2.6 (0.2) | 5.6 (0.6) | 4.1 (0.6) | 4.2 (0.6) | 4.3 (0.6) | 2.7 (0.6) |
| Weak Enhancer | 0.26 (0.01) | 0.05 (0.00) | 0.03 (0.00) | 0.05 (0.01) | 0.04 (0.01) | 0.04 (0.01) | 0.03 (0.01) | 0.05 (0.01) |
| 2.1% | 12.4 (0.4) | 2.2 (0.1) | 1.4 (0.1) | 2.2 (0.4) | 1.7 (0.4) | 1.7 (0.4) | 1.5 (0.4) | 2.4 (0.5) |
| Concordance with | 1.000 | 0.998 | 0.999 | 1.000 | 1.000 | 1.000 | 1.000 | 1.000 |
| UKBb Panel estimates | 0.999 | 0.985 | 0.992 | 1.000 | 0.999 | 0.999 | 0.999 | 0.998 |

**Supplementary Table 6: Estimates of average proportions of SNP heritability and enrichments from 17 Public GWAS.** This is the same as Supplementary Table 5, except averages are across the 17 Public GWAS.

| 14 UKBb GWAS | 1000GP Reference Panel | | | | | | UKBb Reference Panel | | | | | |
|---|---|---|---|---|---|---|---|---|---|---|---|---|
| | BLD-LDAK+Alpha | | BLD-LDAK-Lite+Alpha | | GCTA+Alpha | | BLD-LDAK+Alpha | | BLD-LDAK-Lite+Alpha | | GCTA+Alpha | |
| | Mode | Mean (SD) | Mode | Mean (SD) | Mode | Mean (SD) | Mode | Mean (SD) | Mode | Mean (SD) | Mode | Mean (SD) |
| Body Mass Index | -0.15 | -0.17 (0.06) | -0.20 | -0.18 (0.06) | -0.15 | -0.16 (0.07) | -0.10 | -0.11 (0.06) | -0.10 | -0.11 (0.06) | -0.10 | -0.11 (0.07) |
| Forced Vital Capacity | -0.25 | -0.25 (0.05) | -0.20 | -0.22 (0.05) | -0.30 | -0.31 (0.06) | -0.20 | -0.20 (0.05) | -0.15 | -0.16 (0.05) | -0.25 | -0.24 (0.06) |
| Height | -0.35 | -0.35 (0.03) | -0.30 | -0.29 (0.03) | -0.35 | -0.37 (0.02) | -0.25 | -0.27 (0.03) | -0.20 | -0.20 (0.03) | -0.30 | -0.28 (0.04) |
| Impedance | -0.25 | -0.24 (0.05) | -0.20 | -0.19 (0.05) | -0.20 | -0.21 (0.06) | -0.20 | -0.21 (0.05) | -0.15 | -0.15 (0.05) | -0.20 | -0.19 (0.06) |
| Neuroticism Score | -0.30 | -0.27 (0.09) | -0.30 | -0.27 (0.09) | -0.25 | -0.23 (0.11) | -0.20 | -0.22 (0.10) | -0.20 | -0.21 (0.10) | -0.20 | -0.20 (0.12) |
| Pulse | -0.35 | -0.33 (0.06) | -0.20 | -0.22 (0.07) | -0.30 | -0.28 (0.08) | -0.30 | -0.29 (0.07) | -0.15 | -0.16 (0.08) | -0.25 | -0.25 (0.08) |
| Reaction Time | -0.20 | -0.21 (0.12) | -0.15 | -0.13 (0.12) | -0.10 | -0.08 (0.16) | -0.15 | -0.16 (0.13) | -0.10 | -0.08 (0.13) | -0.10 | -0.07 (0.16) |
| Systolic Blood Pressure | -0.35 | -0.33 (0.05) | -0.25 | -0.23 (0.08) | -0.30 | -0.28 (0.08) | -0.20 | -0.20 (0.05) | -0.20 | -0.20 (0.08) | -0.25 | -0.25 (0.08) |
| College Education | -0.45 | -0.43 (0.05) | -0.40 | -0.42 (0.05) | -0.45 | -0.46 (0.06) | -0.40 | -0.39 (0.06) | -0.40 | -0.38 (0.06) | -0.45 | -0.43 (0.06) |
| Ever Smoked | -0.30 | -0.25 (0.13) | -0.25 | -0.22 (0.13) | -0.30 | -0.28 (0.15) | -0.25 | -0.25 (0.13) | -0.25 | -0.22 (0.14) | -0.30 | -0.29 (0.14) |
| Hypertension | -0.10 | -0.09 (0.11) | 0.00 | 0.00 (0.11) | -0.15 | -0.13 (0.13) | -0.05 | -0.03 (0.12) | 0.05 | 0.08 (0.13) | -0.10 | -0.08 (0.14) |
| Snorer | -0.15 | -0.15 (0.14) | -0.15 | -0.15 (0.15) | -0.30 | -0.27 (0.17) | -0.15 | -0.12 (0.14) | -0.10 | -0.07 (0.16) | -0.25 | -0.23 (0.17) |
| Difficulty Falling Asleep | 0.05 | 0.05 (0.20) | 0.10 | 0.11 (0.21) | 0.20 | 0.20 (0.26) | 0.05 | 0.06 (0.21) | 0.10 | 0.12 (0.21) | 0.20 | 0.21 (0.26) |
| Preference for Evenings | -0.30 | -0.31 (0.08) | -0.30 | -0.29 (0.08) | -0.40 | -0.37 (0.09) | -0.25 | -0.26 (0.09) | -0.25 | -0.23 (0.09) | -0.35 | -0.33 (0.09) |
| Average | | -0.30 (0.02) | | -0.25 (0.02) | | -0.33 (0.02) | | -0.23 (0.02) | | -0.19 (0.02) | | -0.25 (0.02) |

| 17 Public GWAS | 1000GP Reference Panel | | | | | | UKBb Reference Panel | | | | | |
|---|---|---|---|---|---|---|---|---|---|---|---|---|
| | BLD-LDAK+Alpha | | BLD-LDAK-Lite+Alpha | | GCTA+Alpha | | BLD-LDAK+Alpha | | BLD-LDAK-Lite+Alpha | | GCTA+Alpha | |
| | Mode | Mean (SD) | Mode | Mean (SD) | Mode | Mean (SD) | Mode | Mean (SD) | Mode | Mean (SD) | Mode | Mean (SD) |
| Coronary Artery | -0.30 | -0.27 (0.20) | -0.30 | -0.24 (0.22) | 0.00 | 0.04 (0.37) | -0.30 | -0.27 (0.20) | -0.25 | -0.24 (0.22) | -0.05 | 0.00 (0.37) |
| Crohn's Disease | 0.10 | 0.11 (0.18) | 0.25 | 0.27 (0.21) | 0.35 | 0.35 (0.25) | 0.15 | 0.18 (0.20) | 0.35 | 0.34 (0.21) | 0.40 | 0.37 (0.26) |
| Ever Smoked? | -0.15 | -0.10 (0.26) | -0.10 | -0.05 (0.28) | 0.50 | 0.46 (0.38) | 0.00 | 0.06 (0.30) | 0.05 | 0.08 (0.29) | 0.50 | 0.46 (0.38) |
| Inflammatory Bowel | 0.10 | 0.09 (0.15) | 0.25 | 0.25 (0.18) | 0.50 | 0.50 (0.22) | 0.05 | 0.08 (0.15) | 0.25 | 0.25 (0.19) | 0.50 | 0.49 (0.23) |
| Rheumatoid Arthritis | -0.25 | -0.23 (0.13) | 0.00 | 0.02 (0.19) | 0.25 | 0.24 (0.26) | -0.25 | -0.26 (0.12) | -0.10 | -0.09 (0.17) | 0.10 | 0.14 (0.25) |
| Schizophrenia | -0.15 | -0.16 (0.07) | -0.15 | -0.14 (0.07) | 0.10 | 0.11 (0.10) | -0.15 | -0.18 (0.07) | -0.15 | -0.15 (0.07) | 0.05 | 0.05 (0.09) |
| Type 2 Diabetes | 0.00 | 0.04 (0.20) | 0.50 | 0.48 (0.25) | 0.50 | 0.50 (0.18) | 0.10 | 0.11 (0.20) | 0.50 | 0.49 (0.23) | 0.50 | 0.50 (0.18) |
| Bone Mineral Density | 0.25 | 0.26 (0.27) | 0.50 | 0.50 (0.31) | 0.50 | 0.50 (0.20) | 0.20 | 0.22 (0.29) | 0.50 | 0.45 (0.31) | 0.50 | 0.50 (0.22) |
| Body Mass Index | -0.25 | -0.24 (0.06) | -0.25 | -0.24 (0.06) | 0.25 | 0.25 (0.10) | -0.20 | -0.21 (0.06) | -0.20 | -0.19 (0.06) | 0.25 | 0.26 (0.10) |
| Depressive Symptoms | -0.05 | -0.03 (0.27) | 0.05 | 0.10 (0.28) | 0.50 | 0.47 (0.32) | -0.10 | -0.05 (0.25) | 0.05 | 0.08 (0.26) | 0.45 | 0.41 (0.31) |
| Height | -0.20 | -0.21 (0.04) | 0.05 | 0.02 (0.06) | 0.10 | 0.13 (0.07) | -0.20 | -0.19 (0.05) | 0.00 | 0.02 (0.06) | 0.10 | 0.13 (0.07) |
| Menarche Age | -0.15 | -0.17 (0.05) | -0.15 | -0.16 (0.06) | 0.05 | 0.03 (0.07) | -0.20 | -0.18 (0.05) | -0.20 | -0.17 (0.06) | 0.00 | -0.01 (0.07) |
| Menopause Age | 0.20 | 0.22 (0.21) | 0.50 | 0.50 (0.21) | 0.50 | 0.50 (0.22) | 0.25 | 0.27 (0.21) | 0.50 | 0.49 (0.23) | 0.50 | 0.50 (0.23) |
| Neuroticism | 0.00 | 0.00 (0.17) | 0.15 | 0.14 (0.17) | 0.50 | 0.50 (0.18) | -0.05 | -0.01 (0.16) | 0.10 | 0.12 (0.16) | 0.50 | 0.48 (0.20) |
| Subjective Well-Being | -0.45 | -0.43 (0.21) | -0.35 | -0.29 (0.23) | -0.20 | -0.14 (0.34) | -0.45 | -0.42 (0.22) | -0.35 | -0.30 (0.24) | -0.25 | -0.20 (0.32) |
| Waist-Hip Ratio | -0.10 | -0.07 (0.13) | 0.00 | 0.02 (0.16) | 0.50 | 0.50 (0.15) | -0.05 | -0.03 (0.13) | 0.05 | 0.06 (0.16) | 0.50 | 0.50 (0.15) |
| Years Education | -0.10 | -0.08 (0.07) | -0.05 | -0.06 (0.08) | 0.30 | 0.30 (0.11) | -0.05 | -0.04 (0.08) | 0.00 | -0.01 (0.08) | 0.25 | 0.28 (0.11) |
| Average | | -0.16 (0.02) | | -0.07 (0.03) | | 0.21 (0.03) | | -0.15 (0.02) | | -0.06 (0.03) | | 0.18 (0.03) |

**Supplementary Table 7: Genome-wide estimates of $\alpha$.** The BLD-LDAK+Alpha Model contains 67 parameters: 66 $\tau_k$ and $\alpha$. As explained in Supplementary Figure 8, we do not estimate all parameters simultaneously, but instead repeatedly fix $\alpha$ then solve for the $\tau_k$. In total, we consider 31 values of $\alpha$ (from -1 to 0.5, with separation 0.05). In this table, the mode estimate is the value of $\alpha$ which results in highest $logl_{\text{SS}}$; the mean estimate is that obtained by fitting a Gaussian distribution to the 31 realizations of $logl_{\text{SS}}$. Estimates significantly different from zero ($P$<0.05/31) are marked in red.

| 1000G Reference Panel | Average of $logl_{\text{SS}}$ across 14 UKBb GWAS (relative to the null model) | | | | | | | |
|---|---|---|---|---|---|---|---|---|
| **Annotation** | **Step 1** | **Step 2** | **Step 3** | **Step 4** | **Step 5** | **Step 6** | **Step 7** | **Step 8** |
| GCTA | 2124 | 2223 | 2395 | 2406 | **2451** | | | |
| MAF_Adj_Predicted_Allele_Age | 18 | 2253 | 2377 | 2410 | 2446 | 2460 | 2473 | 2478 |
| MAF_Adj_LLD_AFR | 229 | 2230 | 2354 | 2407 | 2446 | 2452 | 2469 | 2476 |
| Recomb_Rate_10kb | 1555 | 2220 | 2341 | 2413 | 2435 | 2452 | **2475** | |
| Nucleotide_Diversity_10kb | 1860 | 2230 | **2406** | | | | | |
| Backgrd_Selection_Stat | 2019 | 2315 | 2375 | **2434** | | | | |
| CpG_Content_50kb | 2214 | **2330** | | | | | | |
| GERP.NS | **2219** | | | | | | | |
| LDAK | 2064 | 2272 | 2346 | 2417 | 2450 | **2468** | | |
| **Maximum improvement** | **2219** | **111** | **76** | **28** | **17** | **17** | **7** | **2** |
| GCTA | 2124 | 2223 | 2395 | 2406 | **2451** | | | |
| MAF_Adj_Predicted_Allele_Age | 18 | 2253 | 2377 | 2410 | 2446 | 2460 | 2473 | 2478 |
| MAF_Adj_LLD_AFR | 229 | 2230 | 2354 | 2407 | 2446 | 2452 | 2469 | 2476 |
| Recomb_Rate_10kb | 1555 | 2220 | 2341 | 2413 | 2435 | 2452 | **2475** | |
| Nucleotide_Diversity_10kb | 1860 | 2230 | **2406** | | | | | |
| Backgrd_Selection_Stat | 2019 | 2315 | 2375 | **2434** | | | | |
| CpG_Content_50kb | 2214 | **2330** | | | | | | |
| GERP.NS | **2219** | | | | | | | |
| LDAK | 2064 | 2272 | 2346 | 2417 | 2450 | **2468** | | |
| **Maximum improvement** | **2219** | **111** | **76** | **28** | **17** | **17** | **7** | **2** |

**Supplementary Table 8: Constructing the BLD-LDAK-Lite Model.** The seven-parameter BLD-LDAK-Lite Model is a reduced version of the 66-parameter BLD-LDAK Model. We started with the nine continuous annotations of the BLD-LDAK Model: the GCTA Annotation ($a_j = 1$), Annotations 1-6 & 73 from Supplementary Table 13, and the LDAK Annotation ($a_j = w_j$), each scaled by $[f_j(1 - f_j)]^{0.75}$. To decide which of these to retain, we used forward stepwise selection, at each step adding to the model the annotation that most improved $logl_{\text{SS}}$ across the 14 UKBb GWAS. The tables above show which annotation was added at each step, and how much this improved $logl_{\text{SS}}$, when using either the 1000 Genome Project reference panel (top) or UKBb panel (bottom). For example, at Step 1, we decided to add the annotation “GERP.NS", while at Step 2, we added the annotation “CpG_Content_50kb". We stopped after Step 7, because it was not possible to substantially improve $logl_{\text{SS}}$ by adding either of the final two annotations.

| Category | Inside Category Mode | Inside Category Mean (SD) | Outside Category Mode | Outside Category Mode | $P$ |
|---|---|---|---|---|---|
| Coding | -0.67 | -0.64 (0.03) | -0.18 | -0.18 (0.02) | $< 10^{-4}$ |
| Conserved | -0.23 | -0.24 (0.03) | -0.23 | -0.21 (0.02) | 0.2077 |
| CTCF | -0.41 | -0.40 (0.03) | -0.23 | -0.22 (0.02) | $< 10^{-4}$ |
| DGF | -0.37 | -0.37 (0.02) | -0.14 | -0.15 (0.02) | $< 10^{-4}$ |
| DHS | -0.40 | -0.40 (0.02) | -0.14 | -0.14 (0.03) | $< 10^{-4}$ |
| Fantom5 Enhancer | -0.63 | -0.52 (0.03) | -0.25 | -0.25 (0.01) | $< 10^{-4}$ |
| Enhancer | -0.27 | -0.30 (0.03) | -0.26 | -0.25 (0.02) | 0.0453 |
| Fetal DHS | -0.49 | -0.47 (0.03) | -0.19 | -0.22 (0.02) | $< 10^{-4}$ |
| H3K27ac (Hnisz) | -0.30 | -0.32 (0.02) | -0.08 | -0.09 (0.03) | $< 10^{-4}$ |
| H3K27ac (PGC2) | -0.32 | -0.33 (0.02) | -0.10 | -0.09 (0.03) | $< 10^{-4}$ |
| H3K4me1 | -0.29 | -0.31 (0.01) | -0.01 | -0.02 (0.03) | $< 10^{-4}$ |
| H3K4me3 | -0.30 | -0.27 (0.02) | -0.16 | -0.18 (0.02) | 0.0045 |
| H3K9ac | -0.37 | -0.37 (0.02) | -0.08 | -0.10 (0.02) | $< 10^{-4}$ |
| Intronic | -0.23 | -0.23 (0.02) | -0.20 | -0.18 (0.02) | 0.0574 |
| Promoter Flanking | -0.31 | -0.30 (0.03) | -0.24 | -0.25 (0.02) | 0.0652 |
| Promoter | -0.60 | -0.60 (0.03) | -0.17 | -0.15 (0.02) | $< 10^{-4}$ |
| Repressed | -0.06 | -0.07 (0.03) | -0.26 | -0.25 (0.02) | $< 10^{-4}$ |
| Super Enhancer | -0.30 | -0.31 (0.02) | -0.19 | -0.17 (0.02) | $< 10^{-4}$ |
| T. Factor Binding Site | -0.40 | -0.38 (0.02) | -0.16 | -0.16 (0.02) | $< 10^{-4}$ |
| Transcribed | -0.22 | -0.23 (0.02) | -0.15 | -0.14 (0.03) | 0.0089 |
| Transcription Start Site | -0.55 | -0.54 (0.03) | -0.22 | -0.21 (0.02) | $< 10^{-4}$ |
| 3-prime UTR | -0.59 | -0.57 (0.03) | -0.20 | -0.21 (0.01) | $< 10^{-4}$ |
| 5-prime UTR | -0.54 | -0.51 (0.03) | -0.24 | -0.22 (0.02) | $< 10^{-4}$ |
| Weak Enhancer | -0.51 | -0.51 (0.03) | -0.22 | -0.22 (0.02) | $< 10^{-4}$ |
| All SNPs | -0.25 | -0.26 (0.01) | | | |

**Supplementary Table 9: Estimates of $\alpha$ across functional categories**. To investigate whether $\alpha$ varies across a functional category, we use the 16-parameter BLD-LDAK-Lite+2Alpha Model (Supplementary Table 13). Within this model are $\alpha_1$ and $\alpha_2$, selection-related parameters corresponding to SNPs inside and outside the annotation, respectively. To estimate $\alpha_1$ and $\alpha_2$, we first solve the BLD-LDAK-Lite+2Alpha Model with $\alpha_1$ and $\alpha_2$ fixed, then we use a grid-search to identify the pair of Gaussian distributions for $\alpha_1$ and $\alpha_2$ that best fit the realizations of $logl_{\text{SS}}$ (Supplementary Figure 8). This table reports, averaged across the 14 UKBb GWAS, the mode estimates of $\alpha_1$ and $\alpha_2$ (the pair of values resulting in highest $logl_{\text{SS}}$), and the mean estimates (those obtained from the grid-search). Functional categories where the estimates of $\alpha_1$ and $\alpha_2$ are significantly different ($P$<0.05/31) are marked in red.

| | Trait | Average Sample Size | Proportion of Cases | Number of of SNPs | Number of Regression SNPs | Unweighted GIF | Weighted GIF |
|---|---|---|---|---|---|---|---|
| 14 UKBb GWAS | Body Mass Index | 129817 | | 4725151 | 4725151 | 1.48 | 1.21 |
| | Forced Vital Capacity | 129817 | | 4725151 | 4725151 | 1.45 | 1.21 |
| | Height | 129817 | | 4725151 | 4725151 | 1.65 | 1.32 |
| | Impedance | 129817 | | 4725151 | 4725151 | 1.49 | 1.23 |
| | Neuroticism Score | 129817 | | 4725151 | 4725151 | 1.24 | 1.10 |
| | Pulse | 129817 | | 4725151 | 4725151 | 1.28 | 1.12 |
| | Reaction Time | 129817 | | 4725151 | 4725151 | 1.20 | 1.09 |
| | Systolic Blood Pressure | 129817 | | 4725151 | 4725151 | 1.31 | 1.13 |
| | College Education | 129817 | 0.35 | 4725151 | 4725151 | 1.41 | 1.19 |
| | Ever Smoked | 129817 | 0.60 | 4725151 | 4725151 | 1.20 | 1.08 |
| | Hypertension | 129817 | 0.22 | 4725151 | 4725151 | 1.22 | 1.10 |
| | Snorer | 129817 | 0.37 | 4725151 | 4725151 | 1.18 | 1.07 |
| | Difficulty Falling Asleep | 129817 | | 4725151 | 4725151 | 1.19 | 1.06 |
| | Preference for Evenings | 129817 | | 4725151 | 4725151 | 1.29 | 1.12 |
| 17 Public GWAS | Coronary Artery[26] | 71445 | 0.25 | 2379111 | 1043550 | 1.14 | 1.06 |
| | Crohn's Disease[27] | 20883 | 0.28 | 8911739 | 1184323 | 1.17 | 1.09 |
| | Ever Smoked?[28] | 74053 | 0.56 | 2413086 | 1060155 | 1.12 | 1.05 |
| | Inflammatory Bowel[27] | 34652 | 0.37 | 9062653 | 1185667 | 1.20 | 1.12 |
| | Rheumatoid Arthritis[29] | 58284 | 0.25 | 8077930 | 1170949 | 1.05 | 1.00 |
| | Schizophrenia[30] | 82315 | 0.43 | 7950070 | 1175491 | 1.60 | 1.35 |
| | Type 2 Diabetes[31] | 152388 | 0.17 | 9289970 | 1186777 | 1.18 | 1.10 |
| | Bone Mineral Density[32] | 32965 | | 7669910 | 1090690 | 1.15 | 1.07 |
| | Body Mass Index[33] | 218799 | | 2491877 | 1078119 | 1.07 | 0.91 |
| | Depressive Symptoms[34] | 161460 | | 6438078 | 1115393 | 1.17 | 1.08 |
| | Height[35] | 230971 | | 2485283 | 1075510 | 2.00 | 1.69 |
| | Menarche Age[36] | 252514 | | 8937573 | 1185550 | 1.65 | 1.37 |
| | Menopause Age[37] | 69360 | | 2380803 | 1044382 | 1.10 | 1.05 |
| | Neuroticism[34] | 170911 | | 6438028 | 1115393 | 1.28 | 1.14 |
| | Subjective Well-Being[34] | 298420 | | 2242512 | 1007956 | 1.17 | 1.08 |
| | Waist-Hip Ratio[38] | 135725 | | 2475648 | 1074208 | 1.05 | 0.95 |
| | Years Education[39] | 328917 | | 7949176 | 1167609 | 1.52 | 1.26 |

**Supplementary Table 10: Details of the 31 GWAS.** We report the average sample size (per-SNP sample sizes were not available for the GWAS marked in red, so for these we report total sample size), and for binary traits, the proportion of cases. Two of the traits are categorical: for difficulty falling asleep, we coded individuals 0, 1 or 2, corresponding to never/rarely, sometimes or always, respectively (with proportions 0.26, 0.47 and 0.27); while for preference for evenings, we coded individuals 0, 1, 2 or 3, corresponding to definitely morning, more morning, more evening, definitely evening, respectively (with proportions 0.26, 0.37, 0.28 and 0.09). We report the number of SNPs (to include a SNP, it must be present in the reference panel) and the number of regression SNPs (for the 14 UKBb GWAS, we used all GWAS SNPs, while for the 17 Public GWAS, we restricted to those present in HapMap 3[40], but not in the major histocompatibility complex). Finally, we report the genomic inflation factor, first calculated using the median test statistic (across the regression SNPs), then using a weighted median test statistic (using SNP weights $1/u_j$, where $u_j = \sum_{l \text{ near } j} r_{jl}^2$). The GWAS marked in red are those which reported using either genomic control or mixed model association analysis, and thus their test statistics are likely to be deflated. Given this likely deflation, and because we had no control over the quality control steps used by the original authors, we always allowed for confounding bias when analysing the 17 Public GWAS.

| | LDSC Recommendation | SumHer Recommendation |
|---|---|---|
| Heritability model | GCTA Model (when estimating SNP heritability, confounding bias or genetic correlations) or Baseline LD Model (when estimating functional enrichments) | Now: BLD-LDAK or BLD-LDAK+Alpha Model<br>Previously: LDAK Model (SNP heritability, confounding bias or genetic correlations) and LDAK+24Fun Model (enrichments) |
| Confounding model | Assume additive inflation (estimate the intercept, $1+A$) | Assume multiplicative inflation (estimate the scaling factor, $C$) |
| Reference panel | Extensive panel, e.g., 10.0 M SNPs with MAF $\geq 0.005$ from 1000 Genomes Project[23] or 13.3 M SNPs with MAC $\geq 5$ (MAF$\geq 0.0007$) from UK10K[41] | Now: Extensive panel (same as LDSC)<br>Previously: High-quality common SNPs, e.g., 4.6 M SNPs with MAF $\geq 0.01$ and info $\geq 0.95$ from Health Retirement Study[42] |
| Regression SNPs | High-quality, MAF $\geq 0.01$ SNPs (if info scores not available, reduce to SNPs present in HapMap 3[40]) | High-quality, MAF $\geq 0.01$ SNPs (if info scores not available, use scores from an alternative GWAS) |
| Heritability SNPs | Reference panel SNPs with MAF $\geq 0.05$ | All reference panel SNPs |
| Enrichment denominator | Proportion of SNPs in category | Expected proportion of SNP heritability under LDAK Model |
| Outlier removal | Exclude MHC (Chr6:25-34 Mb), and SNPs with[6,17] $S_j > \max(80, n_j/1000)$ or $S_j > \max(80, n_j/10000)$ | Exclude MHC (Chr6:25-34 Mb), SNPs with $S_j > n_j/99$ and SNPs in LD with these (within 1 cM and $r^2_{jl} > 0.1$) |

**Supplementary Table 11: Software options.** This table lists the different choices required when running an analysis using LDSC[16] or SumHer,[3] and the recommendations of the respective authors. For our main analysis, we varied the choice of heritability model (the most important option), but followed the recommendations of LDSC for the others. In particular, we used the 1000 Genome Project reference panel provided on the LDSC website (www.github.com/bulik/ldsc), and when regressing $S_j$ on $\mathbb{E}[S_j]$ we either restricted to high-quality MAF $\geq$0.01 SNPs (UKBb GWAS) or HapMap 3 SNPs (Public GWAS). When computing heritability estimates, we restricted to SNPs with MAF $\geq$0.05, and when computing enrichments, we divided the estimated proportion of SNP heritability contributed by a category by the proportion of SNPs the category contained.

Based on the analyses in this paper (Supplementary Table 1), we have changed two of our recommendations: previously, we advised using the LDAK Model and restricting the reference panel to high-quality SNPs[3] (the latter mirrors guidelines when performing heritability analyses using REML[10,12,14,43]); we now advise assuming the BLD-LDAK or BLD-LDAK+Alpha Model and using an extensive reference panel. With regard to the confounding model, we previously argued that it is more accurate to model inflation as multiplicative rather than additive.[3] While this remains our view, one should always prefer to analyze summary statistics from GWAS with only slight confounding, than rely on adjustment for confounding bias.[44]

When calculating enrichments, LDSC compares the estimated proportions of SNP heritability contributed by each category to their expected proportions under the GCTA Model (the latter equal the proportions of SNPs each category contains). We instead preferred to compare estimated proportions to expected proportions under the LDAK Model (to reflect that high-MAF and low-LD SNPs tend to contribute more than low-MAF and high-LD SNPs). While we still consider the latter more appropriate, it generally has only a small impact on estimates of functional enrichments (across the 24 categories, LDAK expectations were typically within 20% of GCTA expectations[3]). Now that we have shown that the BLD-LDAK Model is more realistic than the LDAK Model, it may be preferable to compute expected proportions under the BLD-LDAK-Lite Model (obtained by removing the binary functional annotations from the BLD-LDAK Model). However, considering that BLD-LDAK-Lite expectations will be intermediate between GCTA and LDAK expectations, it should again suffice (and be simpler) to copy LDSC and use GCTA expectations.

Our recommendations regarding regression SNPs and outlier removal are similar to those of LDSC. However, we differ with respect to heritability SNPs (those used when computing heritability estimates). The authors of LDSC[17,45] recommend using only SNPs with MAF $\geq$0.05, because this avoids "extrapolating that $h^2$ per SNP among common variants is the same as $h^2$ per SNP among rare variants." We believe that this is a sensible precaution when using the GCTA, Baseline or Baseline LD Model, each of which assumes that $\mathbb{E}[h^2_j]$ is constant for SNPs with MAF<0.05. However, we consider it unnecessary when using the LDAK, LDAK+24Fun or BLD-LDAK Model, as these better model how heritability varies with MAF (effected by scaling annotations by $[f_j(1-f_j)]^{0.75}$, where $f_j$ is the MAF of SNP $j$). This is evidenced in Supplementary Table 12, which shows that including MAF<0.05 SNPs has less impact on enrichment estimates when assuming the LDAK+24Fun and BLD-LDAK Model than when assuming the Baseline LD Model.

| | 14 UKBb GWAS | | | | | | 17 Public GWAS | | | | | |
|---|---|---|---|---|---|---|---|---|---|---|---|---|
| **Function** | **Baseline LD** | | **LDAK+24Fun** | | **BLD-LDAK** | | **Baseline LD** | | **LDAK+24Fun** | | **BLD-LDAK** | |
| **Proportions of SNPs** | **>0.05** | **ALL** | **>0.05** | **ALL** | **>0.05** | **ALL** | **>0.05** | **ALL** | **>0.05** | **ALL** | **>0.05** | **ALL** |
| Coding | 0.07 | 0.09 | 0.05 | 0.06 | 0.07 | 0.07 | 0.06 | 0.08 | 0.04 | 0.04 | 0.06 | 0.06 |
| 1.4% / 1.6% | 4.9 | 6.3 | 3.6 | 4.0 | 5.2 | 5.2 | 4.2 | 5.3 | 2.5 | 2.7 | 4.4 | 4.4 |
| Conserved | 0.22 | 0.27 | 0.11 | 0.12 | 0.26 | 0.24 | 0.20 | 0.24 | 0.10 | 0.10 | 0.21 | 0.20 |
| 2.6% / 2.9% | 8.7 | 10.4 | 4.4 | 4.6 | 10.0 | 9.4 | 7.9 | 9.2 | 3.9 | 4.1 | 8.0 | 7.7 |
| CTCF | 0.02 | 0.02 | 0.03 | 0.03 | 0.03 | 0.03 | 0.01 | 0.01 | 0.02 | 0.02 | 0.01 | 0.01 |
| 2.4% / 2.4% | 1.0 | 1.0 | 1.1 | 1.1 | 1.3 | 1.3 | 0.3 | 0.3 | 1.0 | 1.0 | 0.4 | 0.4 |
| DGF | 0.28 | 0.30 | 0.21 | 0.21 | 0.30 | 0.29 | 0.26 | 0.28 | 0.22 | 0.22 | 0.28 | 0.28 |
| 13.6% / 13.8% | 2.1 | 2.2 | 1.6 | 1.6 | 2.2 | 2.1 | 1.9 | 2.0 | 1.6 | 1.6 | 2.1 | 2.1 |
| DHS | 0.32 | 0.35 | 0.26 | 0.26 | 0.34 | 0.33 | 0.29 | 0.31 | 0.25 | 0.25 | 0.28 | 0.28 |
| 16.6% / 16.8% | 1.9 | 2.1 | 1.6 | 1.5 | 2.0 | 2.0 | 1.7 | 1.8 | 1.5 | 1.5 | 1.7 | 1.7 |
| FANTOM5 Enhancer | 0.01 | 0.01 | 0.00 | 0.00 | 0.01 | 0.01 | 0.01 | 0.01 | 0.01 | 0.01 | 0.01 | 0.01 |
| 0.4% / 0.4% | 2.8 | 3.0 | 1.1 | 1.0 | 3.1 | 2.9 | 3.1 | 3.4 | 1.4 | 1.3 | 3.2 | 3.1 |
| Enhancer | 0.10 | 0.11 | 0.07 | 0.07 | 0.11 | 0.11 | 0.10 | 0.10 | 0.07 | 0.06 | 0.10 | 0.10 |
| 4.2% / 4.3% | 2.3 | 2.5 | 1.6 | 1.6 | 2.7 | 2.5 | 2.3 | 2.5 | 1.6 | 1.5 | 2.5 | 2.4 |
| Fetal DHS | 0.20 | 0.22 | 0.14 | 0.14 | 0.23 | 0.22 | 0.22 | 0.23 | 0.14 | 0.14 | 0.21 | 0.21 |
| 8.4% / 8.6% | 2.4 | 2.7 | 1.7 | 1.7 | 2.8 | 2.7 | 2.6 | 2.7 | 1.7 | 1.6 | 2.6 | 2.5 |
| H3K27ac (Hnisz) | 0.50 | 0.52 | 0.52 | 0.51 | 0.54 | 0.53 | 0.53 | 0.53 | 0.52 | 0.51 | 0.53 | 0.53 |
| 38.9% / 39.3% | 1.3 | 1.3 | 1.3 | 1.3 | 1.4 | 1.4 | 1.4 | 1.4 | 1.3 | 1.3 | 1.4 | 1.4 |
| H3K27ac (PGC2) | 0.42 | 0.44 | 0.39 | 0.38 | 0.42 | 0.41 | 0.42 | 0.45 | 0.39 | 0.38 | 0.40 | 0.40 |
| 26.9% / 27.3% | 1.5 | 1.6 | 1.4 | 1.4 | 1.6 | 1.5 | 1.6 | 1.7 | 1.4 | 1.4 | 1.5 | 1.5 |
| H3K4me1 | 0.66 | 0.69 | 0.60 | 0.59 | 0.68 | 0.66 | 0.64 | 0.66 | 0.61 | 0.60 | 0.64 | 0.64 |
| 42.4% / 42.9% | 1.6 | 1.6 | 1.4 | 1.4 | 1.6 | 1.6 | 1.5 | 1.6 | 1.4 | 1.4 | 1.5 | 1.5 |
| H3K4me3 | 0.28 | 0.30 | 0.23 | 0.22 | 0.29 | 0.28 | 0.29 | 0.31 | 0.22 | 0.22 | 0.28 | 0.27 |
| 13.3% / 13.7% | 2.1 | 2.3 | 1.7 | 1.7 | 2.2 | 2.1 | 2.2 | 2.4 | 1.7 | 1.6 | 2.1 | 2.0 |
| H3K9ac | 0.33 | 0.36 | 0.25 | 0.25 | 0.35 | 0.33 | 0.28 | 0.30 | 0.24 | 0.23 | 0.29 | 0.28 |
| 12.5% / 12.9% | 2.6 | 2.9 | 2.0 | 2.0 | 2.8 | 2.7 | 2.2 | 2.4 | 1.9 | 1.8 | 2.3 | 2.3 |
| Intronic | 0.43 | 0.44 | 0.47 | 0.47 | 0.43 | 0.43 | 0.44 | 0.45 | 0.47 | 0.47 | 0.45 | 0.45 |
| 38.7% / 39.4% | 1.1 | 1.1 | 1.2 | 1.2 | 1.1 | 1.1 | 1.1 | 1.2 | 1.2 | 1.2 | 1.2 | 1.2 |
| Promoter Flanking | -0.01 | -0.01 | 0.02 | 0.02 | -0.00 | -0.00 | 0.00 | 0.00 | 0.01 | 0.01 | 0.01 | 0.01 |
| 0.8% / 0.9% | -0.6 | -0.7 | 1.8 | 1.8 | -0.2 | -0.0 | 0.1 | 0.1 | 1.8 | 1.8 | 0.8 | 0.8 |
| Promoter | 0.07 | 0.08 | 0.08 | 0.08 | 0.08 | 0.08 | 0.09 | 0.10 | 0.07 | 0.07 | 0.08 | 0.08 |
| 4.6% / 4.8% | 1.5 | 1.7 | 1.7 | 1.7 | 1.8 | 1.7 | 1.9 | 2.1 | 1.5 | 1.5 | 1.8 | 1.8 |
| Repressed | 0.34 | 0.31 | 0.33 | 0.32 | 0.33 | 0.34 | 0.34 | 0.32 | 0.33 | 0.32 | 0.32 | 0.33 |
| 46.1% / 45.3% | 0.7 | 0.7 | 0.7 | 0.7 | 0.7 | 0.7 | 0.7 | 0.7 | 0.7 | 0.7 | 0.7 | 0.7 |
| Super Enhancer | 0.25 | 0.26 | 0.25 | 0.24 | 0.25 | 0.25 | 0.25 | 0.26 | 0.25 | 0.24 | 0.25 | 0.25 |
| 16.7% / 17.0% | 1.5 | 1.5 | 1.5 | 1.5 | 1.5 | 1.5 | 1.5 | 1.5 | 1.5 | 1.4 | 1.5 | 1.5 |
| T. Factor Binding Site | 0.24 | 0.26 | 0.20 | 0.20 | 0.28 | 0.27 | 0.26 | 0.28 | 0.21 | 0.20 | 0.27 | 0.27 |
| 13.1% / 13.3% | 1.8 | 2.0 | 1.5 | 1.5 | 2.1 | 2.1 | 2.0 | 2.1 | 1.6 | 1.5 | 2.1 | 2.0 |
| Transcribed | 0.38 | 0.39 | 0.43 | 0.44 | 0.39 | 0.39 | 0.39 | 0.40 | 0.42 | 0.43 | 0.41 | 0.41 |
| 34.6% / 35.3% | 1.1 | 1.1 | 1.2 | 1.3 | 1.1 | 1.1 | 1.1 | 1.2 | 1.2 | 1.2 | 1.2 | 1.2 |
| Transcription Start Site | 0.06 | 0.07 | 0.05 | 0.05 | 0.06 | 0.06 | 0.06 | 0.07 | 0.04 | 0.04 | 0.06 | 0.06 |
| 1.8% / 1.9% | 3.4 | 3.7 | 2.8 | 2.7 | 3.5 | 3.3 | 3.4 | 3.7 | 2.4 | 2.3 | 3.4 | 3.3 |
| 3'UTR | 0.03 | 0.04 | 0.03 | 0.03 | 0.03 | 0.03 | 0.03 | 0.04 | 0.02 | 0.02 | 0.03 | 0.03 |
| 1.1% / 1.2% | 3.0 | 3.5 | 2.3 | 2.4 | 2.8 | 2.7 | 3.1 | 3.6 | 1.7 | 1.8 | 2.9 | 2.9 |
| 5'UTR | 0.02 | 0.03 | 0.01 | 0.02 | 0.02 | 0.02 | 0.01 | 0.02 | 0.01 | 0.01 | 0.01 | 0.01 |
| 0.5% / 0.6% | 4.3 | 5.1 | 2.6 | 2.8 | 4.1 | 3.9 | 2.6 | 3.1 | 1.9 | 2.1 | 2.4 | 2.4 |
| Weak Enhancer | 0.03 | 0.03 | 0.03 | 0.03 | 0.04 | 0.04 | 0.04 | 0.05 | 0.03 | 0.03 | 0.05 | 0.05 |
| 2.1% / 2.1% | 1.5 | 1.6 | 1.4 | 1.4 | 1.7 | 1.7 | 2.1 | 2.2 | 1.6 | 1.6 | 2.5 | 2.4 |
| Concordance correlation | 0.994 | | 1.000 | | 0.999 | | 0.996 | | 1.000 | | 1.000 | |
| coefficient between pairs | 0.968 | | 0.994 | | 0.997 | | 0.970 | | 0.992 | | 0.999 | |

**Supplementary Table 12: Sensitivity of enrichment estimates to the choice of heritability SNPs.** In general, the heritability SNPs (those used when computing heritability estimates[17]) are the 6.0 M reference panel SNPs with MAF>0.05. Here we investigate the impact on the estimated proportions of SNP heritability and enrichments (top and bottom values in each cell, respectively) of the 24 functional categories if we instead use all 10.0 M SNPs in the reference panel. We find that the change affects estimates from the LDAK+24Fun and BLD-LDAK Models only a little, but can affect estimates from the Baseline LD Model substantially. For more discussion, see Supplementary Table 11.

| | Heritability Model | $K$ | Defintion | Notes |
|---|---|---|---|---|
| Existing Models | GCTA | 1 | $\mathbb{E}[h_j^2] = \tau_1$ | |
| | GCTA-LDMS-R | 20 | $\mathbb{E}[h_j^2] = \sum_{k=1}^{20} I_{jk}\,\tau_k$ | $I_{jk}$ indicates whether SNP j belongs to Bin $k$ |
| | GCTA-LDMS-I | 20 | $\mathbb{E}[h_j^2] = \sum_{k=1}^{20} I'_{jk}\,\tau_k$ | $I'_{jk}$ indicates whether SNP j belongs to Bin $k$ |
| | GCTA+1Fun$_g$ | 2 | $\mathbb{E}[h_j^2] = \tau_1 + c_{j(16+g)}\,\tau_2$ | Contains one function indicator from Baseline LD Model |
| | Baseline | 53 | $\mathbb{E}[h_j^2] = \tau_1 + \sum_{k=2}^{53} c_{j(15+k)}\,\tau_k$ | Contains Annotations 17-68 of Baseline LD Model |
| | Baseline LD | 75 | $\mathbb{E}[h_j^2] = \tau_1 + \sum_{k=2}^{75} c_{j(k-1)}\,\tau_k$ | |
| | LDAK | 1 | $\mathbb{E}[h_j^2] = w_j p_j^{0.75}\,\tau_1$ | |
| | LDAK-OneFun$_g$ | 2 | $\mathbb{E}[h_j^2] = w_j p_j^{0.75}(\tau_1 + c_{j(16+g)}\,\tau_2)$ | Contains one function indicator from Baseline LD Model |
| | LDAK+24Fun | 25 | $\mathbb{E}[h_j^2] = w_j p_j^{0.75}(\tau_1 + \sum_{k=2}^{25} c_{j(15+k)}\,\tau_k)$ | Contains 24 function indicators of Baseline LD Model |
| Novel Models | LDAK-Thin | 1 | $\mathbb{E}[h_j^2] = I_j p_j^{0.75}\,\tau_1$ | $I_j$ indicates whether SNP $j$ remains after thinning duplicates |
| | BLD-LDAK | 66 | $\mathbb{E}[h_j^2] = p_j^{0.75}(\tau_1 + \sum_{k=2}^{7} c_{j(k-1)}\,\tau_k$ $+ \sum_{k=8}^{65} c_{j(k+9)}\,\tau_k + w_j\,\tau_{66})$ | Contains Annotations 1-6 & 17-74 from Baseline LD Model |
| | BLD-LDAK+Alpha | 67 | $\mathbb{E}[h_j^2] = p_j^{1+\alpha}(\tau_1 + \sum_{k=2}^{7} c_{j(k-1)}\,\tau_k$ $+ \sum_{k=8}^{65} c_{j(k+9)}\,\tau_k + w_j\,\tau_{66})$ | Adds $\alpha$ to BLD-LDAK Model |
| Reduced Models | BLD-LDAK-Lite | 7 | $\mathbb{E}[h_j^2] = p_j^{0.75}(\tau_1 + \sum_{k=2}^{6} c_{jS_{k-1}}\,\tau_k + w_j\,\tau_7)$ | Contains Annotations 3-6 & 73 from Baseline LD Model |
| | BLD-LDAK-Lite+1Fun$_g$ | 9 | $\mathbb{E}[h_j^2] = p_j^{0.75}(\tau_1 + \sum_{k=2}^{6} c_{jS_{k-1}}\,\tau_k + w_j\,\tau_7$ $+ c_{j(16+g)}\,\tau_8 + c_{j(40+g)}\,\tau_9)$ | Adds to BLD-LDAK-Lite Model one function indicator and the corresponding 500 bp buffer from Baseline LD Model |
| | BLD-LDAK-Lite+Alpha | 8 | $\mathbb{E}[h_j^2] = p_j^{1+\alpha}(\tau_1 + \sum_{k=2}^{6} c_{jS_{k-1}}\,\tau_k + w_j\,\tau_7)$ | Adds $\alpha$ to BLD-LDAK-Lite Model |
| | BLD-LDAK-Lite+2Alpha$_g$ | 16 | $\mathbb{E}[h_j^2] = c_{j(16+g)}\,p_j^{1+\alpha_1}(\tau_1 + \sum_{k=2}^{6} c_{jS_{k-1}}\,\tau_k + w_j\,\tau_7)$ $+ (1 - c_{j(16+g)})\,p_j^{1+\alpha_2}(\tau_8 + \sum_{k=9}^{13} c_{jS_{k-8}}\,\tau_k + w_j\,\tau_{14})$ | Stratifies BLD-LDAK-Lite+Alpha based on one function indicator from Baseline LD Model |

**Supplementary Table 13: Heritability models.** $K$ is the number of parameters, $c_{j1}, c_{j2}, \ldots, c_{j74}$ are the 74 annotations of the Baseline LD Model (Supplementary Table 14), $w_j$ is the LDAK weighting of SNP $j$, $f_j$ is its MAF, $p_j = f_j(1-f)$ and the set $S = \{3, 4, 5, 6, 73\}$. LDAK weightings should be computed using only high-quality SNPs (therefore $w_j = 0$ for low- and moderate-quality SNPs). There are 24 versions of the GCTA+1Fun, LDAK+1Fun and BLD-LDAK-Lite+1Fun Models (one for each functional category of SNPs). Both the GCTA-LDMS-R and GCTA-LDMS-I Models divide SNPs 20 ways based on LD and MAF; when dividing based on LD, the GCTA-LDMS-R Model uses regional LD scores,[46] while the GCTA-LDMS-I Model uses per-SNP LD scores.[2] In Supplementary Table 15 we show that using 20 bins (4 LD tranches and 5 MAF tranches) results in significantly higher $logl_{\mathrm{SS}}$ than using 8 bins (4 LD tranches and 2 MAF tranches). The LDAK-Thin Model is a simplified version of the LDAK Model; after thinning the SNPs to ensure no pair remains within 100 kb with $r_{jl}^2 > 0.98$, those that remain are given weighting one. The BLD-LDAK-Lite Model uses only seven of the 66 annotations of the BLD-LDAK Model (Supplementary Table 8 explains how we used forward stepwise selection to decide which of the annotations to retain); the BLD-LDAK-Lite+1Fun Model then adds back in one function indicator and its corresponding 500 bp buffer, while the BLD-LDAK-Lite+Alpha Model instead adds in $\alpha$. The BLD-LDAK-Lite+2Alpha Model concatenates two versions of the BLD-LDAK-Lite+Alpha Model: one where only SNPs within a functional category contribute heritability, and one where only SNPs outside the category contribute.

| | | Annotation | % SNPs | | Annotation | % SNPs |
|---|---|---|---|---|---|---|
| LD-Rel. | 1 | MAF_Adj_Predicted_Allele_Age | NA | 2 | MAF_Adj_LLD_AFR | NA |
| | 3 | Recomb_Rate_10kb | NA | 4 | Nucleotide_Diversity_10kb | NA |
| | 5 | Backgrd_Selection_Stat | NA | 6 | CpG_Content_50kb | NA |
| MAF Indicators | 7 | MAFbin1 ($0.05 < MAF \leq 0.07$) | 6.1 | 8 | MAFbin2 ($0.07 < MAF \leq 0.10$) | 6.0 |
| | 9 | MAFbin3 ($0.10 < MAF \leq 0.13$) | 5.9 | 10 | MAFbin4 ($0.13 < MAF \leq 0.17$) | 6.0 |
| | 11 | MAFbin5 ($0.17 < MAF \leq 0.21$) | 5.9 | 12 | MAFbin6 ($0.21 < MAF \leq 0.26$) | 5.9 |
| | 13 | MAFbin7 ($0.26 < MAF \leq 0.32$) | 5.9 | 14 | MAFbin8 ($0.32 < MAF \leq 0.38$) | 6.0 |
| | 15 | MAFbin9 ($0.38 < MAF \leq 0.44$) | 6.0 | 16 | MAFbin10 ($0.44 < MAF \leq 0.50$) | 5.9 |
| Function Indicators | 17 | Coding_UCSC | 1.6 | 18 | Conserved_LindbladToh | 2.9 |
| | 19 | CTCF_Hoffman | 2.4 | 20 | DGF_ENCODE | 13 |
| | 21 | DHS_Trynka | 16 | 22 | Enhancer_Andersson | 0.4 |
| | 23 | Enhancer_Hoffman | 4.3 | 24 | FetalDHS_Trynka | 8.6 |
| | 25 | H3K27ac_Hnisz | 39 | 26 | H3K27ac_PGC2 | 27 |
| | 27 | H3K4me1_Trynka | 42 | 28 | H3K4me3_Trynka | 13 |
| | 29 | H3K9ac_Trynka | 12 | 30 | Intron_UCSC | 39 |
| | 31 | PromoterFlanking_Hoffman | 0.9 | 32 | Promoter_UCSC | 4.8 |
| | 33 | Repressed_Hoffman | 45 | 34 | SuperEnhancer_Hnisz | 16 |
| | 35 | TFBS_ENCODE | 13 | 36 | Transcr_Hoffman | 35 |
| | 37 | TSS_Hoffman | 1.9 | 38 | UTR_3_UCSC | 1.2 |
| | 39 | UTR_5_UCSC | 0.6 | 40 | WeakEnhancer_Hoffman | 2.1 |
| Other Annotations | 41 | Coding_UCSC.extend.500 | 6.7 | 42 | Conserved_LindbladToh.extend.500 | 33 |
| | 43 | CTCF_Hoffman.extend.500 | 7.1 | 44 | DGF_ENCODE.extend.500 | 54 |
| | 45 | DHS_Trynka.extend.500 | 49 | 46 | Enhancer_Andersson.extend.500 | 1.9 |
| | 47 | Enhancer_Hoffman.extend.500 | 9.1 | 48 | FetalDHS_Trynka.extend.500 | 28 |
| | 49 | H3K27ac_Hnisz.extend.500 | 42 | 50 | H3K27ac_PGC2.extend.500 | 33 |
| | 51 | H3K4me1_Trynka.extend.500 | 60 | 52 | H3K4me3_Trynka.extend.500 | 25 |
| | 53 | H3K9ac_Trynka.extend.500 | 23 | 54 | Intron_UCSC.extend.500 | 40 |
| | 55 | PromoterFlanking_Hoffman.extend.500 | 3.4 | 56 | Promoter_UCSC.extend.500 | 5.9 |
| | 57 | Repressed_Hoffman.extend.500 | 70 | 58 | SuperEnhancer_Hnisz.extend.500 | 17 |
| | 59 | TFBS_ENCODE.extend.500 | 34 | 60 | Transcr_Hoffman.extend.500 | 76 |
| | 61 | TSS_Hoffman.extend.500 | 3.6 | 62 | UTR_3_UCSC.extend.500 | 2.8 |
| | 63 | UTR_5_UCSC.extend.500 | 2.8 | 64 | WeakEnhancer_Hoffman.extend.500 | 8.9 |
| | 65 | DHS_peaks_Trynka | 11 | 66 | H3K4me1_peaks_Trynka | 17 |
| | 67 | H3K4me3_peaks_Trynka | 4.3 | 68 | H3K9ac_peaks_Trynka | 4.0 |
| | 69 | Super_Enhancer_Vahedi | 2.2 | 70 | Super_Enhancer_Vahedi.extend.500 | 2.2 |
| | 71 | Typical_Enhancer_Vahedi | 2.2 | 72 | Typical_Enhancer_Vahedi.extend.500 | 2.7 |
| | 73 | GERP.NS | NA | 74 | GERP.RSsup4 | 0.9 |

**Supplementary Table 14: Baseline LD Model SNP annotations** The Baseline LD Model[6] contains 74 annotations, which can be divided into 6 LD-related annotations, 10 MAF indicators, 24 function indicators and 34 auxiliary annotations (predominantly indicators of buffer regions for the functional categories). The names are those provided in the annotation files on the LDSC website (www.github.com/bulik/ldsc). For binary annotations, we report the percentage of SNPs with value 1 (i.e., the number of SNPs within the corresponding category). For more details of each annotation, see the correspondence of Gazal et al. and earlier publications.[6,17] Note that in the main text, to be consistent with previous publications,[3,10,17] we refer to there being 24 functional categories, when there are in fact 26 (Annotations 69 & 71 are included in the Baseline LD Model,[6] but were not in the earlier Baseline Model[17]).

| Heritability Model | K | 14 UKBb GWAS | 17 Public GWAS | Heritability Model | K | 14 UKBb GWAS | 17 Public GWAS |
|---|---|---|---|---|---|---|---|
| GCTA-LDMS-R | 20 | 2277 | 640 | GCTA-LDMS-R (8 partitions) | 8 | 2233 | 592 |
| GCTA-LDMS-I | 20 | 2283 | 625 | GCTA-LDMS-I (8 partitions) | 8 | 2238 | 576 |
| LDAK | 1 | 2160 | 517 | LDAK+24Fun | 25 | 2324 | 704 |
| LDAK-Spread | 1 | 2160 | 518 | LDAK-Spread+24Fun | 25 | 2333 | 716 |
| LDAK-Thin | 1 | 2278 | 617 | LDAK-Thin+24Fun | 25 | 2508 | 861 |
| LDAK-Thin-Spread | 1 | 2278 | 617 | LDAK-Thin-Spread+24Fun | 25 | 2522 | 874 |
| LDAK-Quick | 1 | 2297 | 643 | LDAK-Quick+24Fun | 25 | 2509 | 881 |

**Supplementary Table 15: Average $logl_{\text{SS}}$ for alternative heritability models.** We report the increase in $logl_{\text{SS}}$ relative to the null model, averaged across the 14 UKBb or 17 Public GWAS. The top two rows show that for the GCTA-LDMS Models, it is better to use 5 MAF tranches[46] (in total 20 LD-MAF bins) than 2 MAF tranches[2] (in total 8 LD-MAF bins); for example, when assuming the GCTA-LDMS-I Model, the average increase in $logl_{\text{SS}}$ is 47 (justifying the inclusion of 12 extra parameters).

The bottom five rows consider alternative versions of the LDAK weightings $w_j$. The standard weightings are computed by first thinning the SNPs to exclude duplicates (SNP pairs within 100 kb with $r^2_{jl} > 0.98$), then seeking an approximate solution to $Cw = 1$, where $C$ is a matrix of SNP-SNP correlations.[14] These weightings tend to be sparse; for example, of the 4.7 M UKBb SNPs, only 800 k get non-zero weighting. The LDAK-Thin weightings are obtained by thinning duplicate SNPs, then giving those that remain weighting one. These weightings are less sparse than the standard weightings; of the UKBb SNPs, 1.4 M get non-zero weighting. The LDAK-Quick weightings are obtained by no longer thinning duplicates, but adding `-quick-weights YES` when calculating weightings (they are equal to $1/u_j$, where $u_j = \sum_{l \text{ near } j} r^2_{jl}$). These weightings will be non-zero for all SNPs.

Gazal *et al.*[1] suggested that when estimating functional enrichments, the LDAK-Quick weightings should be preferred to the standard weightings (this was because they found that switching the weightings resulted in estimates closer to those from the Baseline LD Model). With $logl_{\text{SS}}$, it is straightforward to test their suggestion; in support, we find that the LDAK-Quick+24Fun Model fits better than the LDAK+24Fun Model (it has average $logl_{\text{SS}}$ 174 higher). Gazal *et al.* thought that the improvement was because the LDAK-Quick weightings are non-sparse. However, this is only a small factor, evidenced by the fact that the sparse LDAK-Thin weightings perform approximately as well as the LDAK-Quick weightings. Further, we can reduce the sparsity of the LDAK weightings by adding the option `-spread YES` (LDAK will then identify SNPs with non-zero weighting, and "spread" their weight evenly across all duplicate SNPs). Spreading the standard weightings increases the number of SNPs with non-zero weighting from 800 k to 2.1 M, while spreading the LDAK-Thin weightings ensures that all SNPs have non-zero weighting. We see that spreading the weightings only improves the LDAK+24Fun and LDAK-Thin+24Fun Models by a small amount (average $logl_{\text{SS}}$ increases by 11 and 14, respectively). Instead, we believe that the primary reason why the LDAK-Quick weightings are superior to the standard weightings is because the former penalize high-LD SNPs less harshly, resulting in a heritability model intermediate between the GCTA and LDAK Models.

| Heritability Model | K | 14 UKBb | 17 Public |
|---|---|---|---|
| Baseline LD Model | 75 | 0 | 0 |
| Add LDAK Model annotation, $w_j\,[f_j(1-f_j)]^{0.75}$ | 76 | 9 | 12 |
| Add $w_j$, scale all annotations by $[f_j(1-f_j)]^{0.75}$ | 76 | 57 | 64 |
| Add $w_j$, scale annotations by $[f_j(1-f_j)]^{0.75}$, remove 10 MAF indicators (this is the BLD-LDAK Model) | 66 | 50 | 55 |
| BLD-LDAK+Alpha Model | 67 | 51 | 57 |
| BLD-LDAK Model + 4 LD-R partitions | 69 | 53 | 59 |
| BLD-LDAK Model + 20 LD-R partitions | 85 | 64 | 72 |
| BLD-LDAK Model + 4 LD-I partitions | 69 | 52 | 58 |
| BLD-LDAK Model + 20 LD-I partitions | 85 | 62 | 70 |

**Supplementary Table 16: Constructing the BLD-LDAK Model.** Values report the increase in $logl_{\text{SS}}$ relative to the Baseline LD Model, averaged across either the 14 UKBb or 17 Public GWAS. The BLD-LDAK Model is obtained by adding to the Baseline LD Model the LDAK weighting $w_j$, scaling all annotations by $p^{0.75}$, then removing the ten MAF indicators. The first two steps increase average $logl_{\text{SS}}$ by 61 (justifying the inclusion of the extra parameter), whereas removing the MAF indicators reduces average $logl_{\text{SS}}$ by only 8 (indicating that there is little reason to keep the corresponding ten parameters). The final four rows show that the improvement in $logl_{\text{SS}}$ is insufficient to justify including either the 4 LD or 20 LD-MAF bins from the GCTA-LDMS-R and GCTA-LDMS-I Models (note that to avoid colinearity, we only add in 3 and 19 of the bins, respectively).
